# Microbes with higher metabolic independence are enriched in human gut microbiomes under stress

**DOI:** 10.1101/2023.05.10.540289

**Authors:** Iva Veseli, Yiqun T. Chen, Matthew S. Schechter, Chiara Vanni, Emily C. Fogarty, Andrea R. Watson, Bana Jabri, Ran Blekhman, Amy D. Willis, Michael K. Yu, Antonio Fernàndez-Guerra, Jessika Füssel, A. Murat Eren

## Abstract

A wide variety of human diseases are associated with loss of microbial diversity in the human gut, inspiring a great interest in the diagnostic or therapeutic potential of the microbiota. However, the ecological forces that drive diversity reduction in disease states remain unclear, rendering it difficult to ascertain the role of the microbiota in disease emergence or severity. One hypothesis to explain this phenomenon is that microbial diversity is diminished as disease states select for microbial populations that are more fit to survive environmental stress caused by inflammation or other host factors. Here, we tested this hypothesis on a large scale, by developing a software framework to quantify the enrichment of microbial metabolisms in complex metagenomes as a function of microbial diversity. We applied this framework to over 400 gut metagenomes from individuals who are healthy or diagnosed with inflammatory bowel disease (IBD). We found that high metabolic independence (HMI) is a distinguishing characteristic of microbial communities associated with individuals diagnosed with IBD. A classifier we trained using the normalized copy numbers of 33 HMI-associated metabolic modules not only distinguished states of health versus IBD, but also tracked the recovery of the gut microbiome following antibiotic treatment, suggesting that HMI is a hallmark of microbial communities in stressed gut environments.

## Introduction

The human gut is home to a diverse assemblage of microbial cells that form complex communities (Coyte, Schluter, and Foster 2015). This gut microbial ecosystem is established almost immediately after birth and plays a lifelong role in human wellbeing by contributing to immune system maturation and functioning (Belkaid and Hand 2014; Maynard et al. 2012), extracting dietary nutrients (Hijova 2019), providing protection against pathogens (Khosravi and Mazmanian 2013), metabolizing drugs (M. Zimmermann et al. 2019), and more (Knight et al. 2017). There is no universal definition of a healthy gut microbiome (Fan and Pedersen 2021), but associations between host disease states and changes in microbial community composition have sparked great interest in the therapeutic potential of gut microbes (Cani 2018; Sorbara and Pamer 2022) and led to the emergence of hypotheses that directly link disruptions of the gut microbiome to non-communicable diseases of complex etiology (Byndloss and Bäumler 2018).

Inflammatory bowel diseases (IBDs), which describe a heterogeneous group of chronic inflammatory disorders (Shan, Lee, and Chang 2022), represent an increasingly common health risk around the globe (Kaplan 2015). Understanding the role of gut microbiota in IBD has been a major area of focus in human microbiome research. Studies focusing on individual microbial taxa that typically change in relative abundance in IBD patients have proposed a range of host-microbe interactions that may contribute to disease manifestation and progression (Joossens et al. 2011; Schirmer et al. 2019; Henke et al. 2019; Machiels et al. 2014). However, even within well-constrained cohorts, a large proportion of variability in the taxonomic composition of the microbiota is unexplained, and the proportion of variability explained by disease status is low (Gevers et al. 2014; Schirmer et al. 2018; Lloyd-Price et al. 2019; Khan et al. 2019). As neither individual taxa nor broad changes in microbial community composition yield effective predictors of disease (Knox et al. 2019; M. Lee and Chang 2021), the role of gut microbes in the etiology of IBD – or the extent to which they are bystanders to disease – remains unclear (Khan et al. 2019).

The marked decrease in microbial diversity in IBD is often associated with the loss of Firmicutes populations and an increased representation of a relatively small number of taxa, such as *Bacteroides*, *Enterococcaceae*, and others (Prindiville et al. 2000; Saitoh et al. 2002; Sartor 2006; Rhodes 2007; Devkota et al. 2012; Machiels et al. 2014; Vineis et al. 2016; Lloyd-Price et al. 2019). Why a handful of taxa that also typically occur in healthy individuals in lower abundances (M. Lee and Chang 2021; Nishida et al. 2018) tend to dominate the IBD microbiome is a fundamental but open question to gain insights into the ecological underpinnings of the gut microbial ecosystem under IBD. Going beyond taxonomic summaries, a recent metagenome-wide metabolic modeling study revealed a significant loss of cross-feeding partners as a hallmark of IBD, where microbial interactions were disrupted in IBD-associated microbial communities compared to those found in healthy individuals (Marcelino et al. 2023). This observation is in line with another recent work that proposed that the extent of ‘metabolic independence’ (characterized by the genomic presence of a set of key metabolic modules for the synthesis of essential nutrients) is a determinant of microbial survival in IBD (Watson et al. 2023). It is conceivable that the disrupted metabolic interactions among microbes observed in IBD (Marcelino et al. 2023) indicates an environment that lacks the ecosystem services provided by a complex network of microbial interactions, and selects for those organisms that harness high metabolic independence (HMI) (Watson et al. 2023). This interpretation offers an ecological mechanism to explain the dominance of populations with specific metabolic features in IBD. However, this proposed mechanism warrants further investigation.

Here we implemented a high-throughput, taxonomy-independent strategy to estimate metabolic capabilities of microbial communities directly from metagenomes and investigate whether the enrichment of populations with high metabolic independence predicts IBD in the human gut. We benchmarked our findings using representative genomes associated with the human gut and their distribution in healthy individuals and those who have been diagnosed with IBD. Our results suggest that high metabolic potential (indicated by a set of 33 largely biosynthetic metabolic pathways) provides enough signal to consistently distinguish gut microbiomes under stress from those that are in homeostasis, providing deeper insights into adaptive processes initiated by stress conditions that promote rare members of gut microbiota to dominance during disease.

## Results and Discussion

We compiled 2,893 publicly-available stool metagenomes from 13 different studies, 5 of which explicitly studied the IBD gut microbiome (Supplementary Table 1a-c). The average sequencing depth varied across individual datasets (4.2 Mbp to 60.3 Mbp, with a median value of 21.4 Mbp, Supplementary Table 1c). To improve the sensitivity and accuracy of our downstream analyses that depend on metagenomic assembly, we excluded samples with less than 25 million reads, resulting in a set of 408 relatively deeply-sequenced metagenomes from 10 studies (26.4 Mbp to 61.9 Mbp, with a median value of 37.0 Mbp, Supplementary Table 1b, Supplementary File 1, Methods), which we *de novo* assembled individually. The final dataset included individuals who were healthy (n=229), diagnosed with IBD (n=101), or suffered from other gastrointestinal conditions ("non-IBD", n=78). In accordance with previous observations of reduced microbial diversity in IBD (Kostic, Xavier, and Gevers 2014; Nagalingam and Lynch 2012; Knox et al. 2019), the estimated number of populations based on the occurrence of bacterial single-copy core genes present in these metagenomes was higher in healthy individuals than those diagnosed with IBD (Supplementary Figure 1, Supplementary Table 1).

### Estimating normalized copy numbers of metabolic pathways from metagenomic assemblies

Gaining insights into microbial metabolism requires accurate estimates of pathway presence/absence and completion. While a myriad of tools address this task for single genomes (Machado et al. 2018; Aziz et al. 2008; Arkin et al. 2018; Palù et al. 2022; Shaffer et al. 2020; Geller-McGrath et al. 2023; Zorrilla et al. 2021; Zhou et al., n.d.; J. Zimmermann, Kaleta, and Waschina 2021), working with complex environmental metagenomes poses additional challenges due to the large number of organisms that are present in metagenomic assemblies. A few tools can estimate community-level metabolic potential from metagenomes without relying on the reconstruction of individual population genomes or reference-based approaches (Ye and Doak 2009; Karp et al. 2021) (Supplementary Table 5). These high-level summaries of pathway presence and redundancy in a given environment are suitable for most surveys of metabolic capacity, particularly for microbial communities of similar richness. However, since the frequency of observed metabolic modules will increase as the number of distinct microbial populations in a habitat increases, investigations of metabolic determinants of survival across environmental conditions with substantial differences in microbial richness may suffer from ambiguous observations from quantitative data. For instance, the estimated copy number of a given metabolic module may be identical between two metagenomes but its enrichment may be relatively higher in the metagenome with a lower alpha diversity, revealing its potential role in overcoming environment-specific selective pressures that influence an entire community. Working solely with raw copy numbers of metabolic modules without a normalization step that considers the microbial richness will thus shroud potentially critical insights. To quantify the differential enrichment of metabolic modules between metagenomes generated from healthy individuals and those from individuals diagnosed with IBD, we implemented a new software framework (https://anvio.org/m/anvi-estimate-metabolism) that reconstructs metabolic modules from genomes and metagenomes and then calculates the per-population copy number (PPCN) of modules in metagenomes to account for potential differences in microbial richness (Methods, Supplementary File 1 and 2). Briefly, the PPCN estimates the proportion of microbes in a community with a particular metabolic capacity (Figure 1, Supplementary Figure 2) by estimating the number of microbial populations in a given sample using single-copy core genes (SCGs) without having to reconstruct individual genomes first, thus maximizing the *de novo* recovery of gene content. Our validation of this method using simulated metagenomic data demonstrated that it is accurate in capturing metagenome-level metabolic capacity relative to genome-level metabolic capacity estimated from the same data (Supplementary File 2, Supplementary Table 6).

**Figure 1.**
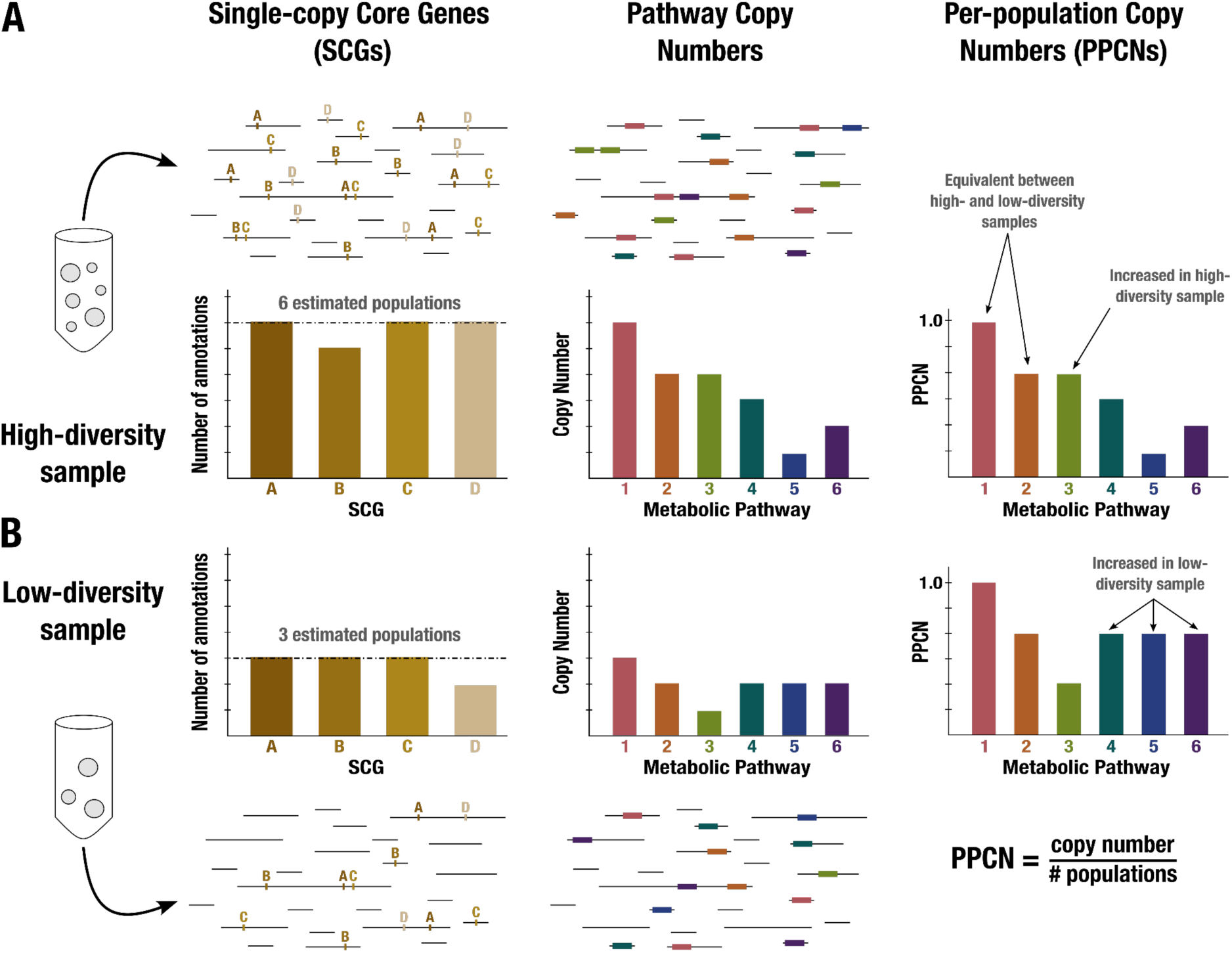
Conceptual diagram of per-population copy number (PPCN) calculation. Each step of the calculation is demonstrated in (**A**) for a sample with high diversity (6 microbial populations) and in (**B**) for a sample with low diversity (3 populations). Metagenome sequences are shown as black lines. The left panel shows the single-copy core genes annotated in the metagenome (indicated by letters), with a barplot showing the counts for different SCGs. The dashed black line indicates the mode of the counts, which is taken as the estimate of the number of populations. The middle panel shows the annotations of metabolic pathways (indicated by boxes and numerically labeled), with a barplot showing the copy number of each pathway (for more details on how this copy number is computed, see Supplementary File 1 and Supplementary Figure 2). The right panel shows the equation for per-population copy number (PPCN), with the barplots indicating the PPCN values for each metabolic pathway in each sample and arrows differentiating between different types of modules based on the comparison of their normalized copy numbers between samples.

### Key biosynthetic pathways are enriched in microbial populations from IBD samples

To gain insight into potential metabolic determinants of microbial survival in the IBD gut environment, we assessed the distribution of metabolic modules within samples from each group (IBD and healthy) with and without using PPCN normalization. A set of 33 metabolic modules were significantly enriched in metagenomes obtained from individuals diagnosed with IBD when PPCN normalization was applied (Figure 2d, 2e). Each metabolic module had an FDR-adjusted p < 2e-10 and an effect size > 0.12 from a Wilcoxon Rank Sum Test comparing IBD and healthy samples. The set included 17 modules that were previously associated with high metabolic independence (Watson et al. 2023) (Figure 2f). However, without PPCN normalization, the signal was masked by the overall higher copy numbers in healthy samples, and the same analysis did not detect higher metabolic potential in microbial populations associated with individuals diagnosed with IBD (Figure 2a), showing weaker differential occurrence between cohorts (Figure 2b, 2c, Supplementary Figure 3). This result suggests that the PPCN normalization is an important step in comparative analyses of metabolisms between samples with disparate levels of diversity.

**Figure 2.**
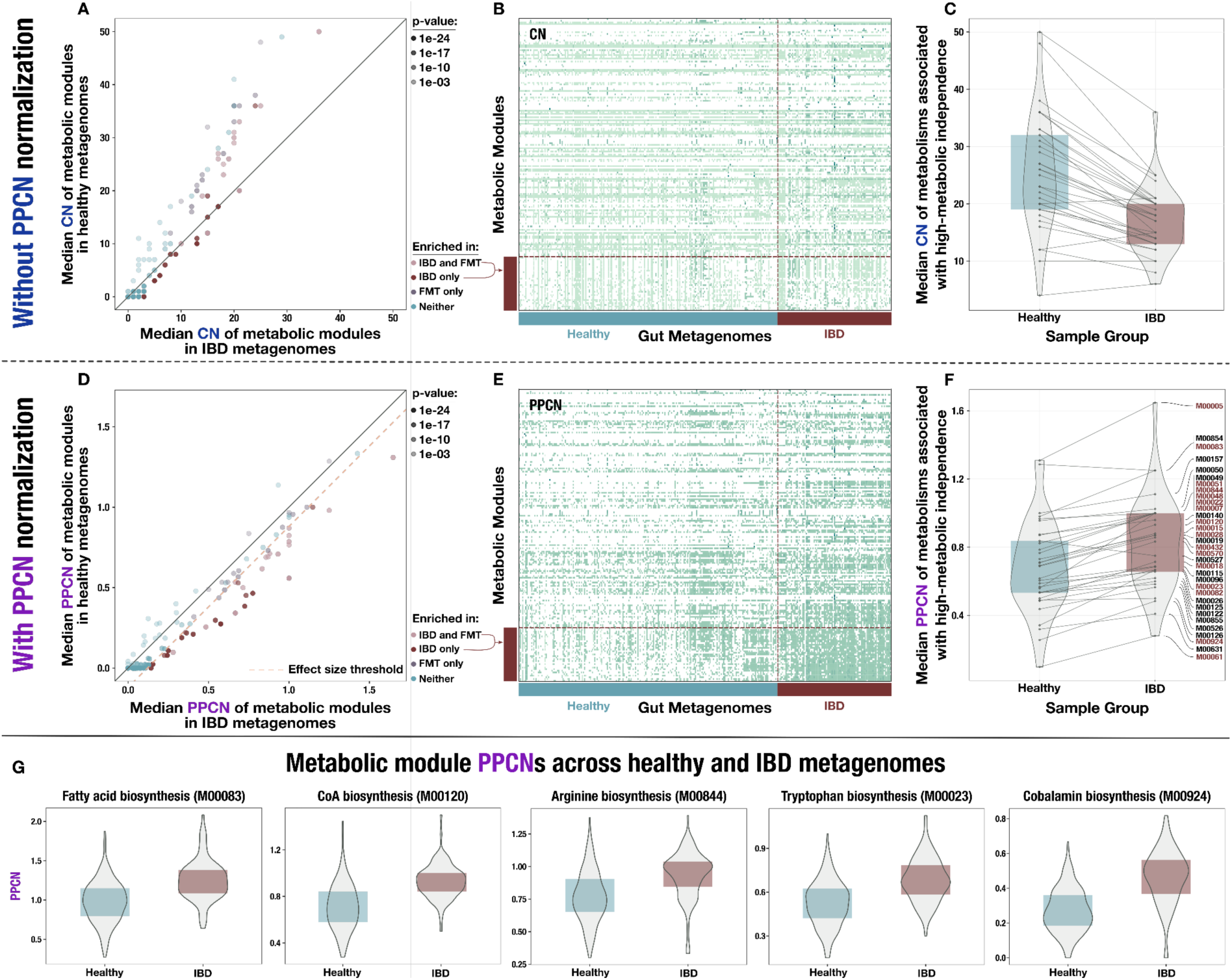
Comparison of metabolic potential across healthy and IBD cohorts. Panels **A – C** show unnormalized copy number data and the remaining panels show normalized per-population copy number (PPCN) data. **A)** Scatterplot of module copy number in IBD samples (x-axis) and healthy samples (y-axis). Transparency of points indicates the p-value of the module in a Wilcoxon Rank Sum test for enrichment (based on PPCN data), and color indicates whether the module is enriched in the IBD samples (in this study), enriched in the good colonizers from the fecal microbiota transplant (FMT) study (Watson et al. 2023), or enriched in both. **B)** Heatmap of unnormalized copy numbers for all modules. IBD-enriched modules are highlighted by the red bar on the left. Sample group is indicated by the blue (healthy) and red (IBD) bars on the bottom. **C)** Boxplots of median copy number for each module enriched in the FMT colonizers from (Watson et al. 2023) in the healthy samples (blue) and the IBD samples (red). Solid lines connect the same module in each plot. **D)** Scatterplot of module PPCN values in IBD samples (x-axis) and healthy samples (y-axis). Transparency and color of points are defined as in panel (A). The pink dashed line indicates the effect size threshold applied to modules when determining their enrichment in IBD. **E)** Heatmap of PPCN values for all modules. Side bars defined as in (B). **F)** Boxplots of median PPCN values for modules enriched in the FMT colonizers from (Watson et al. 2023) in the healthy samples (blue) and the IBD samples (red). Lines defined as in (D). Modules that were also enriched in the IBD samples (in this study) are highlighted in red. **G)** Boxplots of PPCN values for individual modules in the healthy samples (blue) and the IBD samples (red). All example modules were enriched in both this study and in (Watson et al. 2023).

The majority of the metabolic modules that were enriched in the microbiomes of IBD patients encoded biosynthetic capabilities (23 out of 33) that resolved to amino acid metabolism (33%), carbohydrate metabolism (21%), cofactor and vitamin biosynthesis (15%), nucleotide biosynthesis (12%), lipid biosynthesis (6%) and energy metabolism (6%) (Supplementary Table 2a). In contrast to previous reports based on reference genomes (Gevers et al. 2014; Morgan et al. 2012), amino acid synthesis and carbohydrate metabolism were not reduced in the IBD gut microbiome in our dataset. Rather, our results were in accordance with a more recent finding that predicted amino acid secretion potential is increased in the microbiomes of individuals with IBD (Heinken, Hertel, and Thiele 2021).

The metagenome-level enrichment of several key biosynthesis pathways supports the hypothesis that high metabolic independence (HMI) is a determinant of survival for microbial populations in the IBD gut environment. We investigated whether biosynthetic capacity in general was enriched in IBD samples, and 62 out of 88 (70%) biosynthesis pathways described in the KEGG database had a significant enrichment in the IBD sample group at an FDR-adjusted 5% significance level (Supplementary Figure 5d). However, a similar proportion of non-biosynthetic pathways, 63 out of 91 (69%), were also significantly increased in the IBD samples. While biosynthetic capacity is not over-represented in the IBD sample group compared to other types of metabolism, the high proportion of enriched pathways associated with biosynthesis suggests that biosynthetic capacity is important for microbial resilience.

Within our set of 33 pathways that were enriched in IBD, it is notable that all the biosynthesis and central carbohydrate pathways are directly or indirectly linked via shared enzymes and metabolites. Each enriched module shared on average 25.6% of its enzymes and 40.2% of metabolites with the other enriched modules, and overall 18.2% of enzymes and 20.4% of compounds across these pathways were shared (Supplementary Table 2a). Thus, modules may be enriched not just due to the importance of their immediate end products, but also because of their role in the larger metabolic network. The few standalone modules that were enriched included the efflux pump MepA and the beta-Lactam resistance system, which are associated with drug resistance. These capacities may provide an advantage since antibiotics are a common treatment for IBDs (Nitzan et al. 2016), but are not related to the systematic enrichment of biosynthesis pathways that likely provide resilience to general environmental stress rather than to a specific stressor such as antibiotics.

While so far we divided samples into two groups, our dataset also includes individuals who do not suffer from IBD, yet are not healthy either. A recent study using flux balance analysis to model metabolite secretion potential in the dysbiotic, non-dysbiotic, and control gut communities of Crohn’s Disease patients has shown that several predicted microbial metabolic activities align with gradients of host health (Heinken, Hertel, and Thiele 2021). To test whether the HMI signal captures gradients in host health, we included the ‘non-IBD’ group of patients that suffer from gastrointestinal conditions other than IBD in our analysis. The set of 78 samples classified as ‘non-IBD’ indeed represent an intermediate group between healthy individuals and those diagnosed with IBD (Supplementary Figure 5b). While the HMI signal was reduced in ‘non-IBD’ patients, 75% of the pathways enriched in IBD patients were also enriched in the ‘non-IBD’ group compared to healthy individuals. Similarly, when sorting each individual cohort along a health gradient based on cohort descriptions in their respective studies (Supplementary File 1), the relative proportion of metabolic pathways indicative of HMI increased as a function of increasing disease severity (Supplementary Figure 6a). These findings suggest that the HMI signal is sufficiently sensitive to resolve gradients in host health and could serve as a diagnostic tool to monitor changing stress levels in a single individual over time.

Microbiome data generated by different groups can result in systematic biases that may outweigh biological differences between otherwise similar samples (Lozupone et al. 2013; Sinha et al. 2017; Clausen and Willis 2022). The potential impact of such biases constitutes an important consideration for meta-analyses such as ours that analyze publicly available metagenomes from multiple sources. To account for cohort biases, we conducted an analysis of our data on a per-cohort basis, which showed robust differences between the sample groups across multiple cohorts (Supplementary Figure 6b, 6c). Another source of potential bias in our results is due to the representation of microbial functions in genomes in publicly available databases. For instance, we noticed that, independent of the annotation strategy, a smaller proportion of genes resolved to known functions in metagenomic assemblies of the healthy samples compared to the assemblies we generated from the IBD group (Supplementary Figure 4). This highlights the possibility that healthy samples merely appear to harbor less metabolic capabilities due to missing annotations. Indeed, we found that the normalized copy numbers of most metabolic modules were reduced in the healthy group, where 84% of KEGG modules (98 out of 118) have significantly lower median copy numbers (Supplementary Figure 5c, Supplementary File 1). While the presence of a bias between the two cohorts is clear, the source of this bias and its implications are not as clear. One hypothesis that could explain this phenomenon is that the increased proportion of unknown functions in environments where populations with low metabolic independence (LMI) thrive is due to our inability to identify distant homologs of even well-studied functions in poorly studied novel genomes through public databases. If true, this would indeed impair our ability to annotate genes using state-of-the-art functional databases, and bias metabolic module completion estimates. Such a limitation would indeed warrant a careful reconsideration of common workflows and studies that rely on public resources to characterize gene function in complex environments. Another hypothesis that could explain our observation is that the general absence in culture of microbes with smaller genomes (that likely fare better in diverse gut ecosystems) had a historical impact on the characterization of novel functions that represent a relatively larger fraction of their gene repertoire. If true, this would suggest that the unknown functions are unlikely essential for well-studied metabolic capabilities. Furthermore, HMI and LMI genomes may be indistinguishable with respect to the distribution of such novel genes, but the increased number of genes in HMI genomes that resolve to well-studied metabolisms would reduce the proportion of known functions in LMI genomes, and thus in metagenomes where they thrive. While testing these hypotheses falls outside the scope of our work, we find the latter hypothesis more likely due to examples in existing literature that have successfully identified genes that belong to known metabolisms in some of the most obscure organisms via annotation strategies similar to those we have used in our work (Jaffe et al. 2020; Farag et al. 2020).

Taken together, these results (1) demonstrate that the PPCN normalization is an important consideration for investigations of metabolic enrichment in complex microbial communities as a function of microbial diversity, and (2) reveal that the enrichment of HMI populations in an environment offers a high resolution marker to resolve different levels of environmental stress.

### Reference genomes with higher metabolic independence are over-represented in the gut metagenomes of individuals with IBD

So far, our findings demonstrate an overall, metagenome-level trend of increasing HMI within gut microbial communities as a function of IBD status without considering the individual genomes that contribute to this signal. Since the extent of metabolic independence of a microbial genome is a quantifiable trait, we considered a genome-based approach to validating our findings. Given the metagenome-level trends, we expected that the microbial genomes that encode a high number of metabolic modules associated with HMI should be more commonly detected in metagenomes from individuals diagnosed with IBD.

While publicly available reference genomes for microbial taxa will unlikely capture the diversity of individual gut metagenomes, we cast a broad net by surveying the ecology of 19,226 genomes in the Genome Taxonomy Database (GTDB) (Parks et al. 2022) that belonged to three major phyla associated with the human gut environment: Bacteroidetes, Firmicutes, and Proteobacteria (Woting and Blaut 2016; Turnbaugh et al. 2009). We used Human Microbiome Project data (Human Microbiome Project Consortium 2012) to characterize the distribution of these genomes across healthy human gut metagenomes. We used their single-copy core genes to identify genomes that were representative of microbial clades that are systematically detected in the healthy human gut (Figure 3a) and kept those that also occurred in at least 2% of samples from our set of 330 healthy and IBD metagenomes (see Methods). Selection of genomes that are relatively well-detected in the HMP dataset effectively removed taxa that primarily occur outside of the human gut. Of the final set of 338 reference genomes that passed our filters, 258 (76.3%) resolved to Firmicutes, 60 (17.8%) to Bacteroidetes, and 20 (5.9%) to Proteobacteria. Most of these genomes resolved to families common to the colonic microbiota, such as Lachnospiraceae (30.0%), Ruminococcaceae / Oscillospiraceae (23.1%), and Bacteroidaceae (10.1%) (Arumugam et al. 2011), while 5.9% belonged to poorly-studied families with temporary code names (Supplementary Table 3a). Finally, we performed a more comprehensive read recruitment analysis on this smaller set of genomes using all deeply-sequenced metagenomes from cohorts that included healthy, non-IBD, and IBD samples (Figure 3). This provided us with a quantitative summary of the detection patterns of GTDB genome representatives common to the human gut across our dataset.

**Figure 3.**
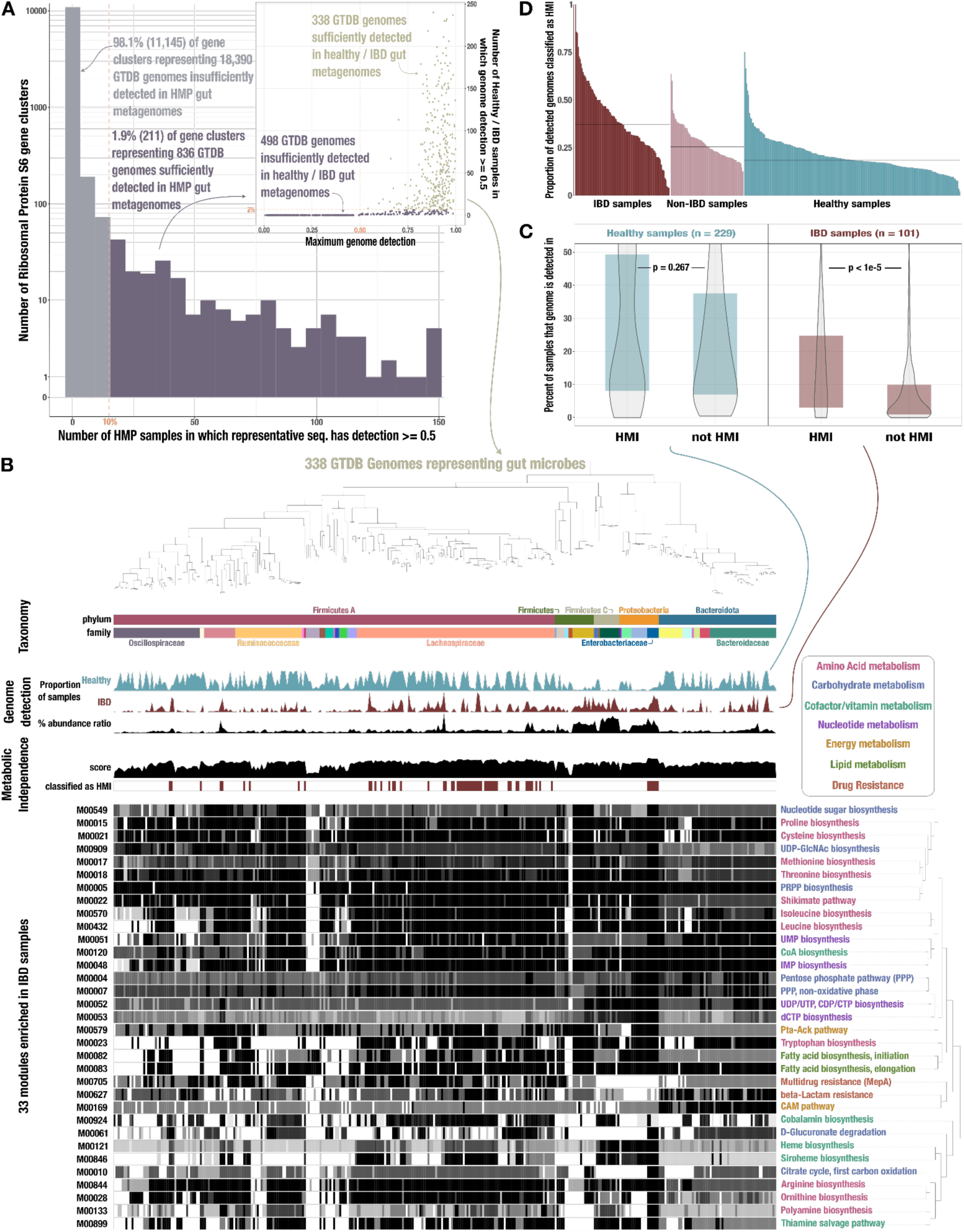
Identification of HMI genomes and their distribution across gut samples. **A)** Histogram of Ribosomal Protein S6 gene clusters (94% ANI) for which at least 50% of the representative gene sequence is covered by at least 1 read (>= 50% ‘detection’) in fecal metagenomes from the Human Microbiome Project (HMP) (Human Microbiome Project Consortium 2012). The dashed line indicates our threshold for reaching at least 50% detection in at least 10% of the HMP samples; gray bars indicate the 11,145 gene clusters that do not meet this threshold while purple bars indicate the 836 clusters that do. The subplot shows data for the 836 genomes whose Ribosomal Protein S6 sequences belonged to one of the passing (purple) gene clusters. The y-axis indicates the number of healthy/IBD gut metagenomes from our set of 330 in which the full genome sequence has at least 50% detection, and the x-axis indicates the genome’s maximum detection across all 330 samples. The dashed line indicates our threshold for reaching at least 50% genome detection in at least 2% of samples; the 338 genomes that pass this threshold are tan and those that do not are purple. The phylogeny of these 338 genomes is shown in **B)** along with the following data, from top to bottom: taxonomic classification as assigned by GTDB; proportion of healthy samples with at least 50% detection of the genome sequence; proportion of IBD samples with at least 50% detection of the genome sequence; square-root normalized ratio of percent abundance in IBD samples to percent abundance in healthy samples; metabolic independence score (sum of completeness scores of 33 HMI-associated metabolic pathways); whether (red) or not (white) the genome is classified as having HMI with a threshold score of 26.4; heatmap of completeness scores for each of the 33 HMI-associated metabolic pathways (0% completeness is white and 100% completeness is black). Pathway name is shown on the right and colored according to its category of metabolism. **C)** Boxplot showing the proportion of healthy (blue) or IBD (red) samples in which genomes of each class are detected >= 50%, with p-values from a Wilcoxon Rank-Sum test on the underlying data. **D)** Barplot showing the proportion of detected genomes (with >= 50% genome sequence covered by at least 1 read) in each sample that are classified as HMI, for each group of samples. The black lines show the median for each group: 37.0% for IBD samples, 25.5% for non-IBD samples, and 18.4% for healthy samples.

We classified each genome as HMI if its average completeness of the 33 HMI-associated metabolic pathways was at least 80%, equivalent to a summed metabolic independence score of 26.4 (Methods). Across all genomes, the mean metabolic independence score was 24.0 (Q1: 19.9, Q3: 25.7). We identified 17.5% (59) of the reference genomes as HMI. HMI genomes were on average substantially larger (3.8 Mbp) than non-HMI genomes (2.9 Mbp) and encoded more genes (3,634 vs. 2,683 genes, respectively), which is in accordance with the reduced metabolic potential of non-HMI populations (Supplementary Table 3a). Our read recruitment analysis showed that HMI reference genomes were present in a significantly higher proportion of IBD samples compared to non-HMI genomes (Figure 3c, p < 1e-5, Wilcoxon Rank Sum test). Similarly, the fraction of HMI populations was significantly higher within a given IBD sample compared to samples classified as ‘non-IBD’ and those from healthy individuals (Figure 3d, p < 1e-24, Kruskal-Wallis Rank Sum test). In contrast, the detection of HMI populations and non-HMI populations was similar in healthy individuals (Figure 3c, p = 0.267, Wilcoxon Rank Sum test). The intestinal environment of healthy individuals likely supports both HMI and non-HMI populations, wherein ‘metabolic diversity’ is maintained by metabolic interactions such as cross feeding. Indeed, loss of cross-feeding interactions in the gut microbiome appears to be associated with a number of human diseases, including IBD (Marcelino et al. 2023). This interpretation is further supported by the fact that the top two HMI-associated pathways are required for the synthesis of cobalamin from glutamate. Auxotrophy for cobalamin biosynthesis is common among gut bacteria that rely on cross-feeding for this essential cofactor (Degnan et al. 2014; Magnúsdóttir et al. 2015; Kelly et al. 2019) (Supplementary File 1).

Overall, the classification of reference gut genomes as HMI and their enrichment in individuals diagnosed with IBD strongly supports the contribution of HMI to stress resilience of individual microbial populations. We note that survival in a disturbed gut environment will likely require a wide variety of additional functions that are not covered in the list of metabolic modules we consider to determine HMI status – for examples, see (Degnan et al. 2014; Martens et al. 2014; Zong et al. 2020; L. Feng et al. 2020; Goodman et al. 2009; Powell et al. 2016). Indeed, there may be many ways for a microbe to be metabolically independent, and our strategy likely failed to identify some HMI populations. Nonetheless, these data suggest that HMI serves as a reliable proxy for the identification of microbial populations that are particularly resilient.

### HMI-associated metabolic potential predicts general stress on gut microbes

Our analysis identified HMI as an emergent property of gut microbial communities associated with individuals diagnosed with IBD. This community-level signal translates to individual microbial populations and provides insights into the microbial ecology of stressed gut environments. HMI-associated metabolic pathways were enriched at the community level, and microbial populations encoding these modules were more prevalent in individuals with IBD than in healthy individuals. Furthermore, the copy number of these pathways and the proportion of HMI populations reflect the severity of environmental stress and translate to host health states (Supplementary Figure 5b, Figure 3d). The ecological implications of these observations suggest that HMI may serve as a predictor of general stress in the human gut environment.

So far, efforts to identify IBD using microbial markers have presented classifiers based on (1) taxonomy in pediatric IBD patients (Papa et al. 2012; Gevers et al. 2014), (2) community composition in combination with clinical data (Halfvarson et al. 2017), (3) untargeted metabolomics and/or species-level relative abundance from metagenomes (Franzosa et al. 2019) and (4) k-mer-based sequence variants in metagenomes that can be linked to microbial genomes associated with IBD (Reiter et al. 2022). Performance varied both between and within studies according to the target classes and data types used for training and validation of each classifier (Supplementary Table 4a). For those studies reporting accuracy, a maximum accuracy of 77% was achieved based on either metabolite profiles (for prediction of IBD-subtype) (Franzosa et al. 2019) or k-mer-based sequence variants (for differentiating between IBD and non-IBD samples) (Reiter et al. 2022). Some studies reported performance as area under the receiver operating characteristic curve (AUROCC), a typical measure of classifier utility describing both sensitivity (ability to correctly identify the disease) and specificity (ability to correctly identify absence of disease). For this metric the highest value was 0.92, achieved by (Franzosa et al. 2019) when using metabolite profiles, with or without species abundance data, for classifying IBD vs non-IBD. However, the majority of these classifiers were trained and tested on a relatively small group of individuals that all come from the same region, i.e. clinical studies confined to a specific hospital. Though some had high performance, they either relied on data that are inaccessible to most laboratories and clinics considering that untargeted metabolomics analyses are difficult to reproduce (Koek et al. 2011; Lin et al. 2020), or they required complex k-mer-based models without the resolution to differentiate gradients in host health (Reiter et al. 2022). These classifiers thus have limited translational potential across global clinical settings and do not provide an ecological framework to explain the observed shifts in community composition and activity. For practical use as a diagnostic tool, a microbiome-based classifier for IBD should rely on an ecologically meaningful, easy to measure, and high-level signal that is robust to host variables like lifestyle, geographical location, and ethnicity. High metabolic independence could potentially fill this gap as a metric related to the ecological filtering that defines microbial community changes in the IBD gut microbiome.

We trained a logistic regression classifier to explore the applicability of HMI as a non-invasive diagnostic tool for IBD. The classifier’s predictors were the per-population copy numbers of IBD-enriched metabolic pathways in a given metagenome. Across the 330 deeply-sequenced IBD and healthy samples included in this analysis, the classifier had high sensitivity and specificity (Figure 4). It correctly identified (on average) 76.8% of samples from individuals diagnosed with IBD and 89.5% of samples representing healthy individuals, for an overall accuracy of 85.6% and an average AUROCC of 0.832 (Supplementary Table 4c). Our model outperforms (Gevers et al. 2014; Halfvarson et al. 2017; Reiter et al. 2022) or has comparable performance to (Franzosa et al. 2019; Papa et al. 2012) the previous attempts to classify IBD from fecal samples in more restrictively-defined cohorts. It also has the advantage of being a simple model, utilizing a relatively low number of features compared to the other classifiers. Thus, HMI shows promise as an accessible diagnostic marker of IBD. Due to the lack of time-series studies that include individuals in the pre-diagnosis phase of IBD development, we cannot test the applicability of HMI to predict IBD onset (Lloyd-Price et al. 2019).

**Figure 4.**
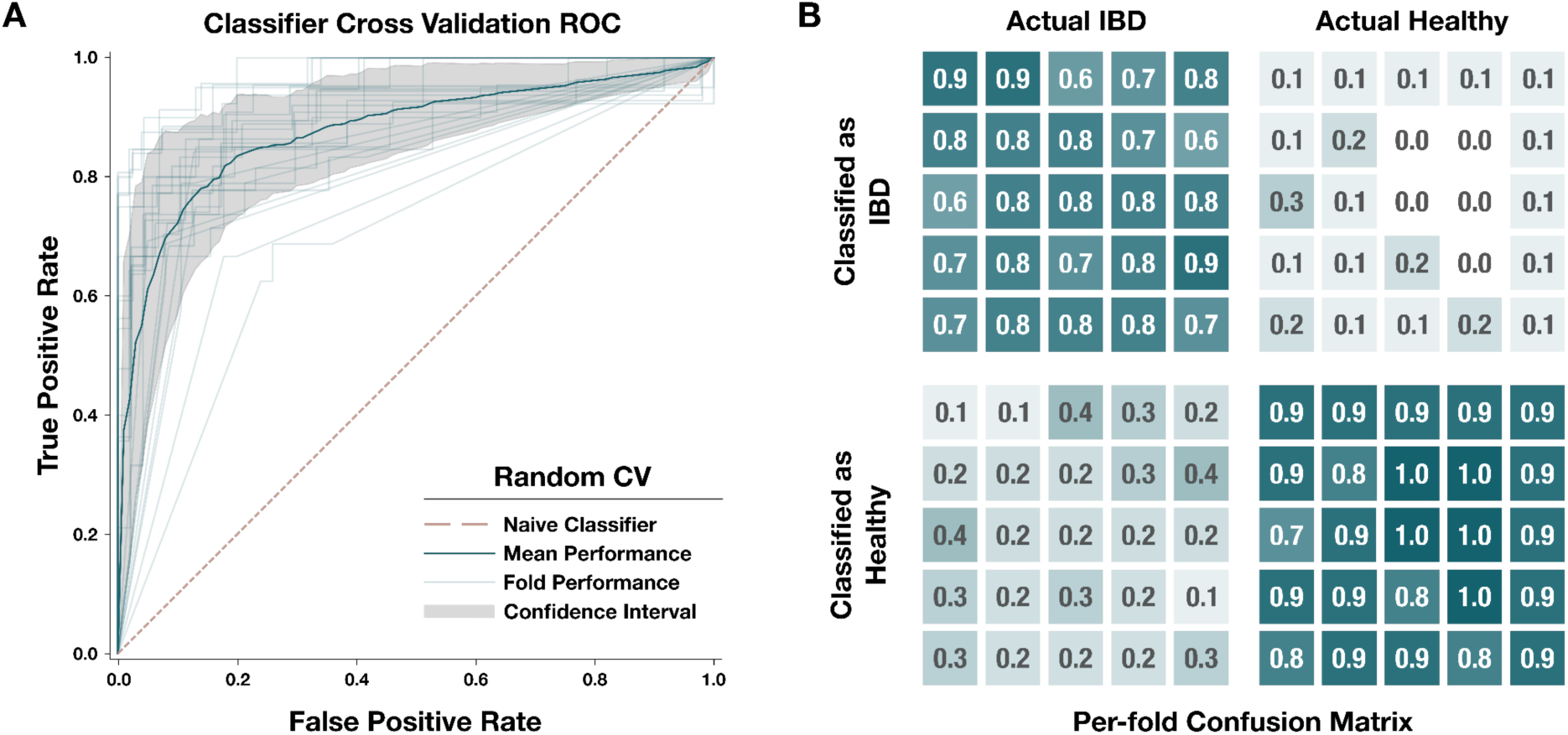
Performance of our metagenome classifier trained on per-population copy numbers of IBD-enriched modules. **A)** Receiver operating characteristic (ROC) curves for 25-fold cross-validation. Each fold used a random subset of 80% of the data for training and the other 20% for testing. In each fold, we calculated a set of IBD-enriched modules from the training dataset and used the PPCN of these modules to train a logistic regression model whose performance was evaluated using the test dataset. Light gray lines show the ROC curve for each fold, the dark blue line shows the mean ROC curve, the gray area delineates the confidence interval for the mean ROC, and the pink dashed line indicates the benchmark performance of a naive (random guess) classifier. **B)** Confusion matrix for each fold of the random cross-validation. Categories of classification, from top left to bottom right, are: true positives (correctly classified IBD samples), false positives (incorrectly classified Healthy samples), false negatives (incorrectly classified IBD samples), and true negatives (correctly classified Healthy samples). Each fold is represented by a box within each category. Opacity of the box indicates the proportion of samples in that category, and the actual proportion is written within the box with one significant digit. Underlying data for this matrix can be accessed in Supplementary Table 4d.

Yet, the gradient of metabolic independence reflected by per-population pathway copy number and the relative increase in the number of HMI populations detected in non-IBD samples (Supplementary Figure 5b, Figure 3d) suggests that the degree of HMI in the gut microbiome may be indicative of general gut stress, such as the stress induced by antibiotic use. Antibiotics can cause long-lasting perturbations of the gut microbiome – including reduced diversity, emergence of opportunistic pathogens, increased microbial load, and development of highly-resistant strains – with potential implications for host health (Ramirez et al. 2020). We applied our metabolism classifier to a metagenomic dataset that reflects the changes in the microbiome of healthy people before, during and up to 6 months following a 4-day antibiotic treatment (Palleja et al. 2018). The resulting pattern of sample classification corresponds to the post-treatment decline and subsequent recovery of species richness documented in the study by (Palleja et al. 2018).

All pre-treatment samples were classified as ‘healthy’ followed by a decline in the proportion of ‘healthy’ samples to a minimum 8 days post-treatment, and a gradual increase until 180 days post treatment, when over 90% of samples were classified as ‘healthy’ (Figure 5, Supplementary Table 4b). These observations support the role of HMI as an ecological driver of microbial resilience during gut stress caused by a variety of environmental perturbations and demonstrate its diagnostic power in reflecting gut microbiome state.

**Figure 5.**
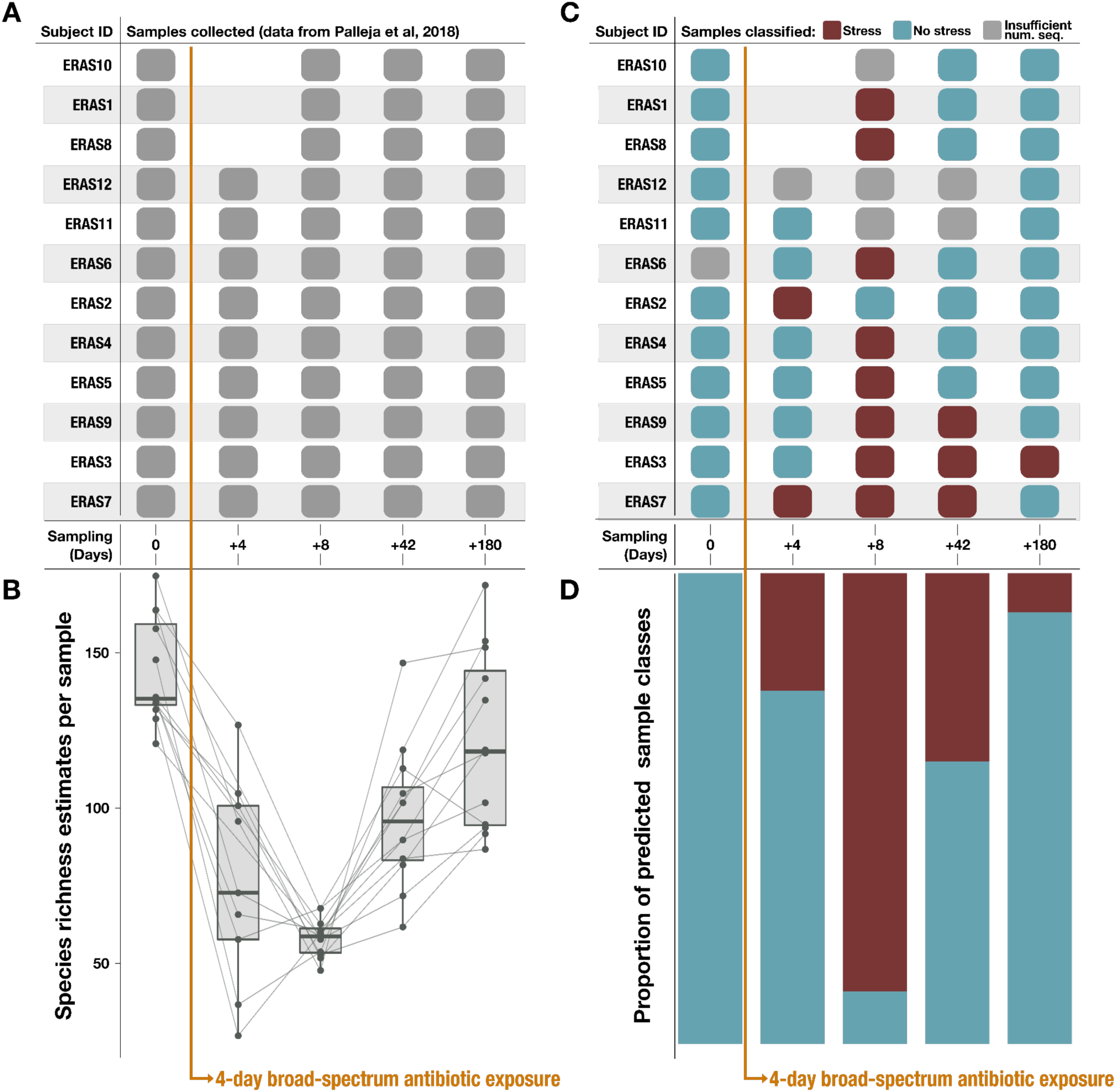
Classification results on an antibiotic time-series dataset from. (Palleja et al. 2018). Note that antibiotic treatment was taken on days 1-4. **A)** Samples collected per subject during the time series. **B)** Species richness data (figure reproduced from (Palleja et al. 2018)). **C)** Classification of each sample by the metabolism classifier profiled in Figure 4. Samples with insufficient sequencing depth were not classified. **D)** Proportion of classes assigned to samples per day in the time series.

## Conclusions

Overall, our observations that stem from the analysis of hundreds of reference genomes, deeply-sequenced gut metagenomes, and multiple categories of human disease states suggest that environmental stress in the human gut – whether it is associated with inflammation, cancer, or antibiotic use – promotes the survival and relative expansion of microbial populations with high metabolic independence. These results establish HMI as a high-level metric to classify gradients of human health states through the gut microbiota that is robust to ethnic, geographical or lifestyle factors. Taken together with recent evidence that models altered ecological relationships within gut microbiomes under stress due to disrupted metabolic cross-feeding (Heinken, Hertel, and Thiele 2021; Marcelino et al. 2023), our data support the hypothesis that the reduction in microbial diversity, or more generally ‘dysbiosis’, is an emergent property of microbial communities responding to disease pathogenesis or other external factors such as antibiotic use that disrupt the gut microbial ecosystem. This paradigm depicts microbes as bystanders by default, rather than perpetrators or drivers of noncommunicable human diseases, and provides an ecological framework to explain the frequently observed reduction in microbial diversity associated with IBD and other noncommunicable human diseases and disorders.

## Methods

A bioinformatics workflow that further details all analyses described below and gives access to reproducible data products is available at the URL https://merenlab.org/data/ibd-gut-metabolism/.

### A new framework for metabolism estimation

We developed a new program ‘anvi-estimate-metabolism‘ (https://anvio.org/m/anvi-estimate-metabolism), which uses gene annotations to estimate ‘completeness’ and ‘copy number’ of metabolic pathways that are defined in terms of enzyme accession numbers. By default, this tool works on metabolic modules from the KEGG MODULE database (Kanehisa et al. 2012, 2023) which are defined by KEGG KOfams (Aramaki et al. 2019), but user-defined modules based on a variety of functional annotation sources are also accepted as input. Completeness estimates describe the percentage of steps (typically, enzymatic reactions) in a given metabolic pathway that are encoded in a genome or a metagenome. Likewise, copy number summarizes the number of distinct sets of enzyme annotations that collectively encode the complete pathway. This program offers two strategies for estimating metabolic potential: a ‘stepwise’ strategy with equivalent treatment for alternative enzymes – i.e, enzymes that can catalyze the same reaction in a given metabolic pathway – and a ‘pathwise’ strategy that accounts for all possible variations of the pathway. The Supplementary File 1 file includes more information on these two strategies and the completeness/copy number calculations. For the analysis of metagenomes, we used stepwise copy number of KEGG modules. Briefly, the calculation of stepwise copy number is done as follows: the copy number of each step in a pathway (typically, one chemical reaction or conversion) is individually evaluated by translating the step definition into an arithmetic expression that summarizes the number of annotations for each required enzyme. In cases where multiple enzymes or an enzyme complex are needed to catalyze the reaction, we take the minimum number of annotations across these components. In cases where there are alternative enzymes that can each catalyze the reaction individually, we sum the number of annotations for each alternative. Once the copy number of each step is computed, we then calculate the copy number of the entire pathway by taking the minimum copy number across all the individual steps. The use of minimums results in a conservative estimate of pathway copy number such that only copies of the pathway with all enzymes present are counted. For the analysis of genomes, we calculated the stepwise completeness of KEGG modules. This calculation is similar to the one described above for copy number, except that the step definition is translated into a boolean expression that, once evaluated, indicates the presence or absence of each step in the pathway. Then, the completeness of the modules is computed as the proportion of present steps in the pathway.

### Metagenomic Datasets and Sample Groups

We acquired publicly-available gut metagenomes from 13 different studies (Le Chatelier et al. 2013; Q. Feng et al. 2015; Franzosa et al. 2019; Lloyd-Price et al. 2019; Qin et al. 2012; Quince et al. 2015; Rampelli et al. 2015; Raymond et al. 2016; Schirmer et al. 2018; Vineis et al. 2016; “BioProject,” n.d.; Wen et al. 2017; Xie et al. 2016). The studies were chosen based on the following criteria: (1) they included shotgun metagenomes of fecal matter (primarily stool, but some ileal pouch luminal aspirate samples (Vineis et al. 2016) are also included); (2) they sampled from people living in industrialized countries (in the case where a study (Rampelli et al. 2015) included samples from hunter-gatherer populations, only the samples from industrialized areas were included in our analysis); (3) they included samples from people with IBD and/or they included samples from people without gastrointestinal (GI) disease or inflammation; and (4) clear metadata differentiating between case and control samples was available. A full description of the studies and samples can be found in Supplementary Table 1a-c. We grouped samples according to the health status of the sample donor. Briefly, the ‘IBD’ group of samples includes those from people diagnosed with Crohn’s disease (CD), ulcerative colitis (UC), or pouchitis. The ‘non-IBD’ group contains non-IBD controls, which includes both healthy people presenting for routine cancer screenings as well as people with benign or non-specific symptoms that are not clinically diagnosed with IBD. Colorectal cancer patients from (Q. Feng et al. 2015) were also put into the ‘non-IBD’ group on the basis that tumors in the GI tract may arise from local inflammation (Kraus and Arber 2009) and represent a source of gut stress without an accompanying diagnosis of IBD. Finally, the ‘HEALTHY’ group contains samples from people without GI-related diseases or inflammation. Note that only control or pre-treatment samples were taken from the studies covering type 2 diabetes (Qin et al. 2012), ankylosing spondylitis (Wen et al. 2017), antibiotic treatment (Raymond et al. 2016), and dietary intervention (“BioProject,” n.d.); these controls were all assigned to the ‘HEALTHY’ group. At least one study (Le Chatelier et al. 2013) included samples from obese people, and these were also included in the ‘HEALTHY’ group.

### Processing of metagenomes

We made single assemblies of most gut metagenomes using the anvi’o metagenomics workflow implemented in the program ‘anvi-run-workflow‘ (Shaiber et al. 2020). This workflow uses Snakemake (Köster and Rahmann 2012), and a tutorial is available at the URL https://merenlab.org/2018/07/09/anvio-snakemake-workflows/. Briefly, the workflow includes quality filtering using ‘iu-filter-quality-minochè (Eren et al. 2013); assembly with IDBA-UD (Peng et al. 2012) (using a minimum contig length of 1000); gene calling with Prodigal v2.6.3 (Hyatt et al. 2010); tRNA identification with tRNAscan-SE v2.0.7 (Chan and Lowe 2019); and gene annotation of ribosomal proteins (Seemann, n.d.), single-copy core gene sets (M. D. Lee 2019), KEGG KOfams (Aramaki et al. 2019), NCBI COGs (Galperin et al. 2021), and Pfam (release 33.1, (Mistry et al. 2021)). The aforementioned annotation was done with programs that relied on HMMER v3.3.2 (Eddy 2011) as well as Diamond v0.9.14.115 (Buchfink, Xie, and Huson 2015). As part of this workflow, all single assemblies were converted into anvi’o contigs databases. Samples from (Vineis et al. 2016) were processed differently because they contained merged reads rather than individual paired-end reads: no further quality filtering was run on these samples, we assembled them individually using MEGAHIT (Li et al. 2015), and we used the anvi’o contigs workflow to perform all subsequent steps described for the metagenomics workflow above. Note that we used a version of KEGG downloaded in December 2020 (for reproducibility, the hash of the KEGG snapshot available via ‘anvi-setup-kegg-kofams‘ is 45b7cc2e4fdc). Additionally, the annotation program ‘anvi-run-kegg-kofams‘ includes a heuristic for annotating hits with bitscores that are just below the KEGG-defined threshold, which is described at https://anvio.org/m/anvi-run-kegg-kofams/.

### Genomic Dataset

We also analyzed microbial genomes from the Genome Taxonomy Database (GTDB), release 95.0 (Parks et al. 2018, 2020). We downloaded all reference genome sequences for the species cluster representatives.

### Processing of GTDB genomes

We converted all GTDB genomes into anvi’o contigs databases and annotated them using the anvi’o contigs workflow, which is similar to the metagenomics workflow described above and uses the same programs for gene identification and annotation.

### Estimation of the number of microbial populations per metagenome

We used single-copy core gene (SCG) sets belonging to each domain of microbial life (Bacteria, Archaea, Protista) (M. D. Lee 2019) to estimate the number of populations from each domain present in a given metagenomic sample. For each domain, we calculated the number of populations by taking the mode of the number of copies of each SCG in the set. We then summed the number of populations from each domain to get a total number of microbial populations within each sample. We accomplished this using SCG annotations provided by ‘anvi-run-hmms‘ (which was run during metagenome processing) and a custom script relying on the anvi’o class ‘NumGenomesEstimator‘ (see reproducible workflow).

### Removal of samples with low sequencing depth

We observed that, at lower sequencing depths, our estimates for the number of populations in a metagenomic sample were moderately correlated with sequencing depth (Supplementary Figure 1, R > 0.5). These estimates rely on having accurate counts of single-copy core genes (SCGs), so we hypothesized that lower-depth samples were systematically missing SCGs, especially from populations with lower abundance. Since accurate population number estimates are critical for proper normalization of pathway copy numbers, keeping these lower-depth samples would have introduced a bias into our metabolism analyses. To address this, we removed samples with low sequencing depth from downstream analyses using a sequencing depth threshold of 25 million reads, such that the remaining samples exhibited a weaker correlation (R < 0.5) between sequencing depth and number of estimated populations. We kept samples for which both the R1 file and the R2 file contained at least 25 million reads (and for the (Vineis et al. 2016) dataset, we kept samples containing at least 25 million merged reads). This produced our final sample set of 408 metagenomes.

### Estimation of normalized pathway copy numbers in metagenomes

We ran ‘anvi-estimate-metabolism‘, in genome mode and with the ‘--add-copy-number‘ flag, on each individual metagenome assembly to compute stepwise copy numbers for KEGG modules from the combined gene annotations of all populations present in the sample. We then divided these copy numbers by the number of estimated populations within each sample to obtain a per-population copy number (PPCN) for each pathway.

### Selection of IBD-enriched pathways

We used a one-sided Mann-Whitney-Wilcoxon test with a FDR-adjusted p-value threshold of p <= 2e-10 on the per-sample PPCN values for each module individually to identify the pathways that were most significantly enriched in the IBD sample group compared to the healthy group. We calculated the median per-population copy number of each metabolic pathway in the IBD samples, and again in the healthy samples. After filtering for p-values <= 2e-10, we also applied a minimum effect size threshold based on the median per-population copy number in each group (M_IBD_ - M_Healthy_ >= 0.12) – this threshold was calculated by taking the mean effect size over all pathways that passed the p-value threshold. The set originally contained 34 pathways that passed both thresholds, but we removed one redundant module (M00006) which represents the first half of another module in the set (M00004).

### Test for enrichment of biosynthesis pathways

We used a one-sided Fisher’s exact test (also known as hypergeometric test, see e.g., (Boyle et al. 2004)) for testing the independence between the metabolic pathways identified to be IBD-enriched (i.e., using the methods described in “Selection of IBD-enriched pathways) and functionality (i.e., pathways annotated to be involved in biosynthesis).

### Pathway comparisons

Because the 33 IBD-enriched pathways were selected using PPCNs of healthy and IBD samples, statistical tests comparing PPCN distributions for these modules need to be interpreted with care, because the hypotheses were selected and tested on the same dataset (Fithian, Sun, and Taylor 2014). Therefore, to assess the statistical validity of the identified IBD-enriched modules, we performed the following repeated sample-split analysis: we first randomly split the IBD and healthy samples into the equal-sized training and validation sets. We select IBD-enriched modules in the training set using the Mann-Whitney-Wilcoxon test, and then compute the p-values on the validation set. We repeat this sample split analysis 1,000 times with an FDR-adjusted p-value threshold of 1e-10 on the first split; most identified modules (89.4%; 95% CI: [87.5%, 91.3%]) on the training sets remain significant at a slightly less stringent threshold (1e-8) on the validation sets. This indicates that the approach we used to identify IBD-enriched modules yields stable and statistically significant results on this dataset.

### Metagenome classification

We trained logistic regression models to classify samples as ‘IBD’ or ‘healthy’ using per-population copy numbers of IBD-enriched modules as features. We ran a 25-fold cross-validation pipeline on the set of 330 healthy and IBD metagenomes in our analysis, using an 80% train – 20% test random split of the data in each fold. The pipeline included selection of IBD-enriched pathways within the training samples using the same strategy as described above, followed by training and testing of a logistic regression model as implemented in the ‘sklearn‘ Python package. We set the ‘penalty‘ parameter of the model to “None” and the ‘max_iter‘ parameter to 20,000 iterations, and we used the same random state in each fold to ensure changes in performance only come from differences in the training data rather than differences in model initialization. To summarize the overall performance of the classifier, we took the mean (over all folds) of each performance metric.

We trained a final classifier using the 33 IBD-enriched pathways selected earlier from the entire set of 330 healthy and IBD metagenomes. We then applied this classifier to the metagenomic samples from (Palleja et al. 2018), which we processed in the same way as the other samples in our analysis (including removal of samples with low sequencing depth and calculation of PPCNs of KEGG modules for use as input features to the classifier model).

### Identification of gut microbial genomes from the GTDB

We took 19,226 representative genomes from the GTDB species clusters belonging to the phyla Firmicutes, Bacteroidetes, and Proteobacteria, which are most common in the human gut microbiome (Woting and Blaut 2016). To evaluate which of these genomes might represent gut microbes in a computationally-tractable manner, we ran the anvi’o ‘EcoPhylo’ workflow (https://anvio.org/m/ecophylo) to contextualize these populations within 150 healthy gut metagenomes from the Human Microbiome Project (HMP) (Human Microbiome Project Consortium 2012). Briefly, the EcoPhylo workflow (1) recovers sequences of a gene family of interest from each genome and metagenomic sample in the analysis, (2) clusters resulting sequences and picks representative sequences using mmseqs2 (Steinegger and Söding 2017), and (3) uses the representative sequences to rapidly summarize the distribution of each population cluster across the metagenomic samples through metagenomic read recruitment analyses. Here, we used the Ribosomal Protein S6 as our gene of interest, since it was the most frequently-assembled single-copy-core gene in our set of GTDB genomes. We clustered the Ribosomal Protein S6 sequences from GTDB genomes at 94% nucleotide identity.

To identify genomes that were likely to represent gut microbes, we selected genomes whose ribosomal protein S6 belonged to a gene cluster where at least 50% of the representative sequence was covered (i.e. detection >= 0.5x) in more than 10% of samples (i.e. n > 15). There are 100 distinct individuals represented in the 150 HMP gut metagenomes – 56 of which were sampled just once and 46 of which were sampled at 2 or 3 time points – so this threshold is equivalent to detecting the genome in 5% - 15% of individuals. From this selection we obtained a set of 836 genomes; however, these were not exclusively gut microbes, as some non-gut populations have similar ribosomal protein S6 sequences to gut microbes and can therefore pass this selection step. To eliminate these, we mapped our set of 330 healthy and IBD metagenomes to the 836 genomes using the anvi’o metagenomics workflow, and extracted genomes whose entire sequence was at least 50% covered (i.e. detection >= 0.5x) in over 2% (n > 6) of these samples. Our final set of 338 genomes was used in downstream analysis.

### Genome phylogeny

To create the phylogeny, we identified the following ribosomal proteins that were annotated in at least 90% (n = 304) of the genomes: Ribosomal_S6, Ribosomal_S16, Ribosomal_L19, Ribosomal_L27, Ribosomal_S15, Ribosomal_S20p, Ribosomal_L13, Ribosomal_L21p, Ribosomal_L20, and Ribosomal_L9_C. We used ‘anvi-get-sequences-for-hmm-hits‘ to extract the amino acid sequences for these genes, align the sequences using MUSCLE v3.8.1551 (Edgar 2004), and concatenate the alignments. We used trimAl v1.4.rev15 (Capella-Gutiérrez, Silla-Martínez, and Gabaldón 2009) to remove any positions containing more than 50% of gap characters from the final alignment. Finally, we built the tree with IQtree v2.2.0.3 (Minh et al. 2020), using the WAG model and running 1,000 bootstraps.

### Determination of HMI status for genomes

We estimated metabolic potential for each genome with ‘anvi-estimate-metabolism‘ (in genome mode) to get stepwise completeness scores for each KEGG module, and then we used the script ‘anvi-script-estimate-metabolic-independencè to give each genome a metabolic independence score based on completeness of the 33 IBD-enriched pathways. Briefly, the latter script calculates the score by summing the completeness scores of each pathway of interest. Genomes were classified as having high metabolic independence (HMI) if their score was greater than or equal to 26.4. We calculated this threshold by requiring these 33 pathways to be, on average, at least 80% complete in a given genome.

### Genome distribution across sample groups

We mapped the gut metagenomes from the healthy, non-IBD, and IBD groups to each genome using the anvi’o metagenomics workflow in reference mode. We used ‘anvi-summarizè to obtain a matrix of genome detection across all samples. We summarized this data as follows: for each genome, we computed the proportion of samples in each group in which at least 50% of the genome sequence was covered by at least 1 read (>= 50% detection). For each sample, we calculated the proportion of detected genomes that were classified as HMI. We also computed the percent abundance of each genome in each sample by dividing the number of reads mapping to that genome by the total number of reads in the sample.

### Visualizations

We used ggplot2 (Wickham 2016) to generate most of the initial data visualizations. The phylogeny and heatmap in Figure 3 were generated by the anvi’o interactive interface and the ROC curves in Figure 4 were generated using the pyplot package of matplotlib (Hunter 2007). These visualizations were refined for publication using Inkscape, an open-source graphical editing software that is available at https://inkscape.org/.

## Data Availability

Accession numbers for publicly available data are listed in our Supplementary Tables at doi:10.6084/m9.figshare.22679080. Our Supplementary Files are also available at doi:10.6084/m9.figshare.22679080. Contigs databases of our assemblies for the 408 deeply-sequenced metagenomes can be accessed at doi:10.5281/zenodo.7872967, and databases for our assemblies of the (Palleja et al. 2018) metagenomes can be accessed at doi:10.5281/zenodo.7897987. Contigs databases of the 338 GTDB gut reference genomes are available at doi:10.5281/zenodo.7883421.

## Acknowledgements

We thank Chris Quince for advice on statistical significance testing. IV acknowledges support from the National Science Foundation Graduate Research Fellowship under Grant No. 1746045; ADW acknowledges support from the National Institutes of General Medical Sciences under R35 GM133420. YTC acknowledges support from the Stanford Data Science Postdoctoral Fellowship. RB acknowledges support from the National Institutes of Health under R35 GM128716. ECF acknowledges support from the University of Chicago International Student Fellowship. Additional support for ECF and AME came from an NIH NIDDK grant (RC2 DK122394) to AME.

## Author Contributions

IV, JF and AME conceived the study. IV developed methodology. IV, MSS and AME developed computational analysis tools. IV, JF, and AME performed formal analyses. YTC supported statistical analyses. RB, BJ, ADW and MKY conducted investigations. ECF, ARW, CV and AFG provided resources. ECF, ARW and IV curated data. IV and AME prepared figures. IV, JF and AME wrote the paper with critical input from all authors. JF and AME supervised the project.

## Supplemental Figures

**Supplementary Figure 1.**
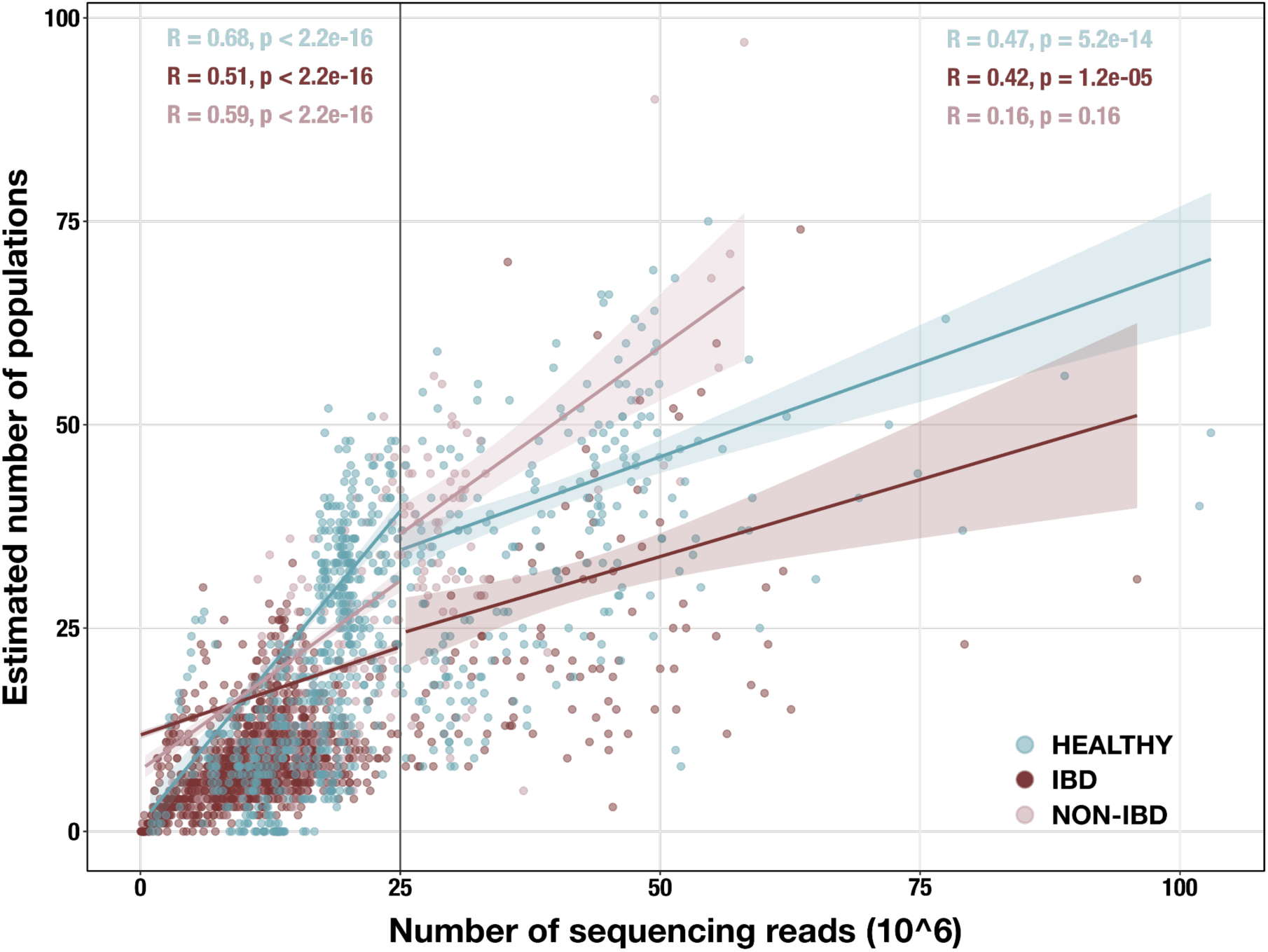
Scatterplot of sequencing depth vs estimated number of microbial populations in each of 2,893 stool metagenomes. Sequencing depth is represented by the number of R1 reads, except for (Vineis et al. 2016) samples, in which case it is the number of merged paired-end reads. The vertical line indicates our sequencing depth threshold of 25 million reads. Per-group Spearman’s correlation coefficients and p-values are shown for the subset of samples with depth < 25 million reads (top left) and for the subset with depth >= 25 million reads (top right). Regression lines are shown for each group in each subset, with standard error indicated by the colored background.

**Supplementary Figure 2.**
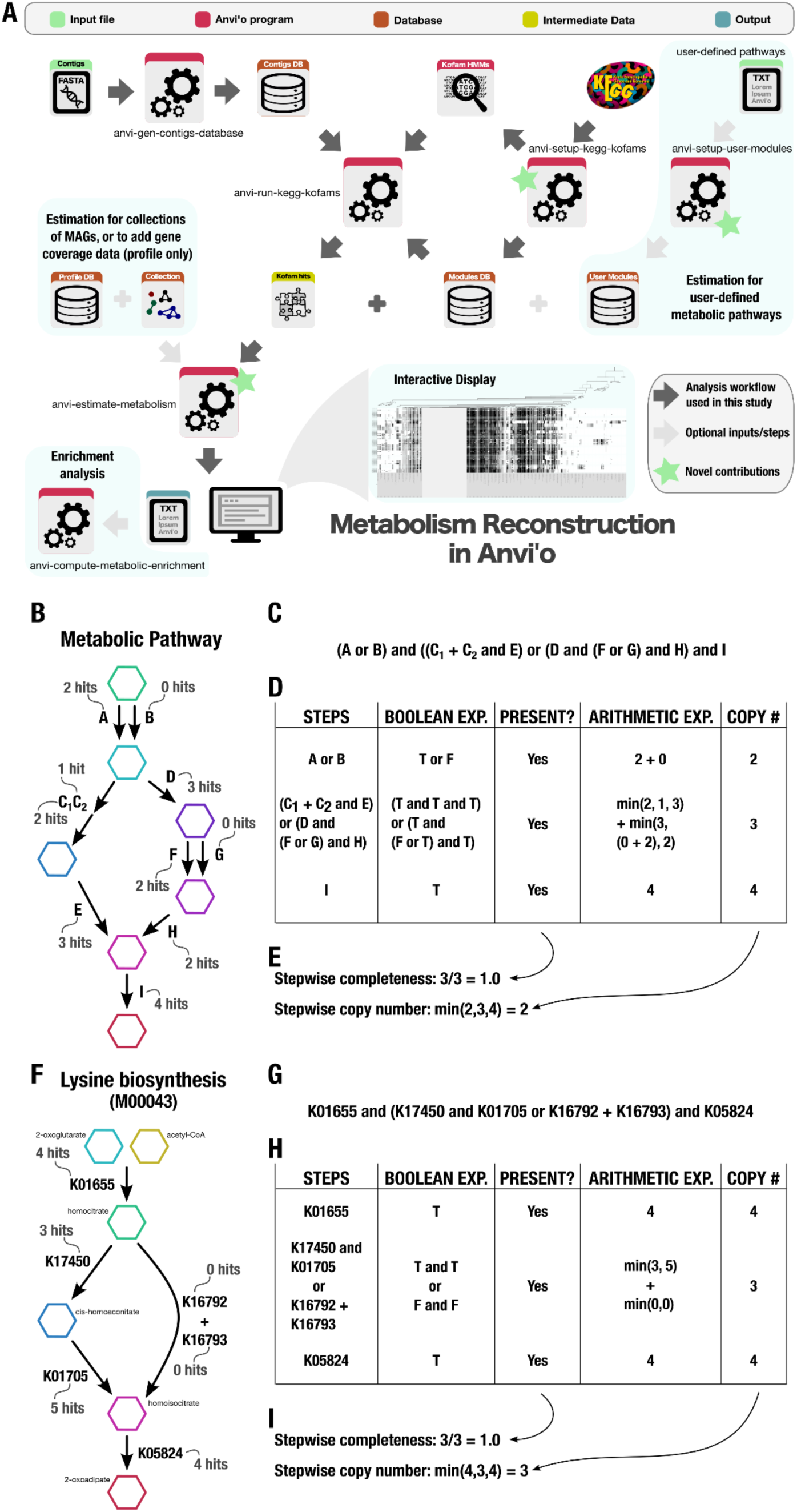
Technical details of the metabolism reconstruction software framework in anvi’o. A) Workflow of metabolism reconstruction programs and their inputs/outputs. Dark arrows indicate the primary analysis path utilized in this study. Blue background indicates optional features in the framework. A demonstration of completeness score and copy number calculations for metabolic pathways (performed by the program ‘anvi-estimate-metabolism‘ is shown using example enzyme annotation data in panels B – E (for a theoretical pathway) and F – I (for a real pathway). **B)** Theoretical metabolic pathway, where hexagons represent metabolites, arrows represent chemical reactions, letters represent enzymes (subscripts indicate enzyme components), and the example number of gene annotation hits for each enzyme is written in gray. **C)** The definition of the theoretical pathway from panel B, written in terms of the required enzymes. **D)** Table showing the major steps in the pathway and example calculations for step presence and copy number. Step presence is calculated by evaluating a boolean expression created from the step definition in which enzymes with > 0 hits are replaced with True (T) and the others with False (F). Step copy number is calculated by evaluating the corresponding arithmetic expression in which the enzymes are replaced with their annotation counts. **E)** Final calculations of completeness score (fraction of present steps) and copy number for the theoretical metabolic pathway. **F – I)** Same as panels B – E, but for KEGG module M00043.

**Supplementary Figure 3.**
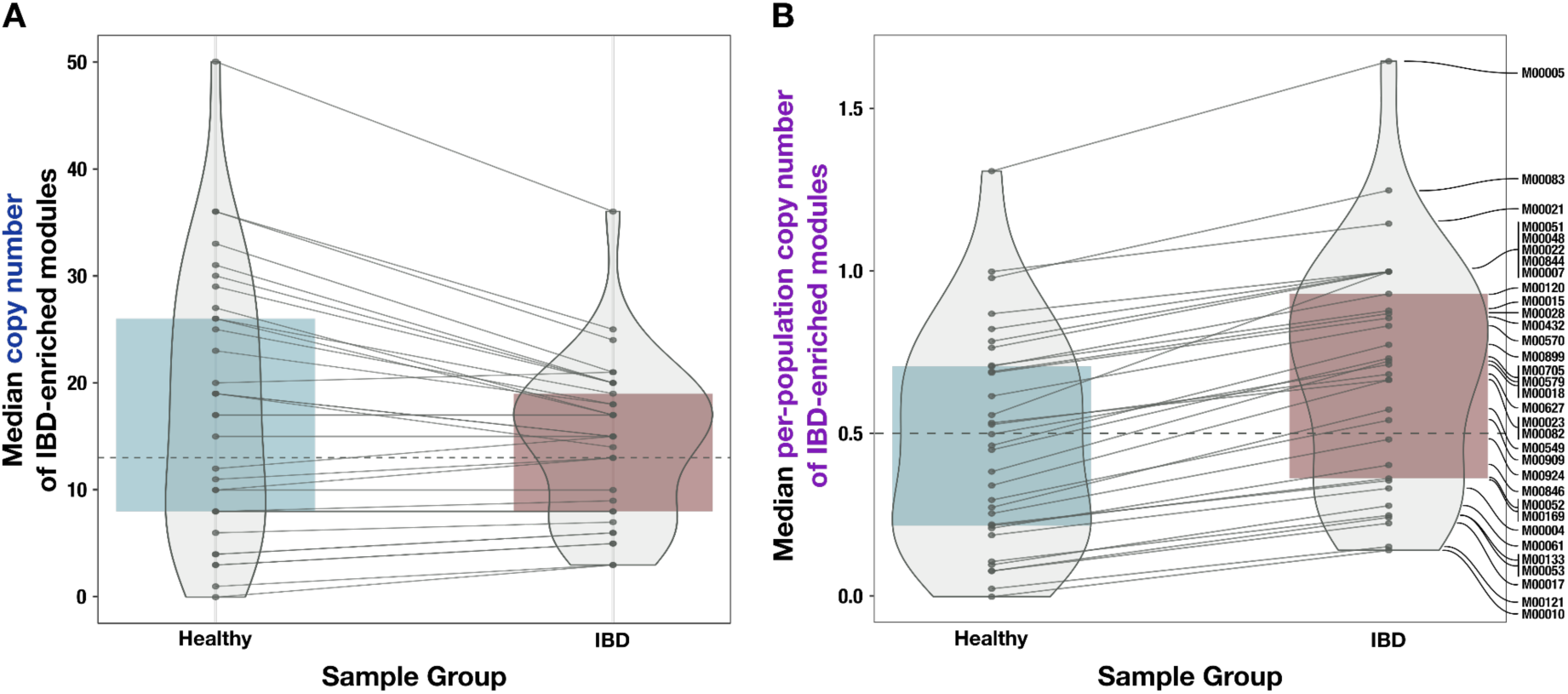
Comparison of unnormalized copy number data and normalized (per-population copy number, or PPCN) data for the IBD-enriched modules. **A)** Boxplot of median copy numbers for each module in the healthy samples (blue) and IBD samples (red). **B)** Boxplots of median PPCN for each module in the healthy samples (blue) and IBD samples (red). Lines connect data points for the same module in each plot. The gray dashed line in each plot indicates the overall median value.

**Supplementary Figure 4.**
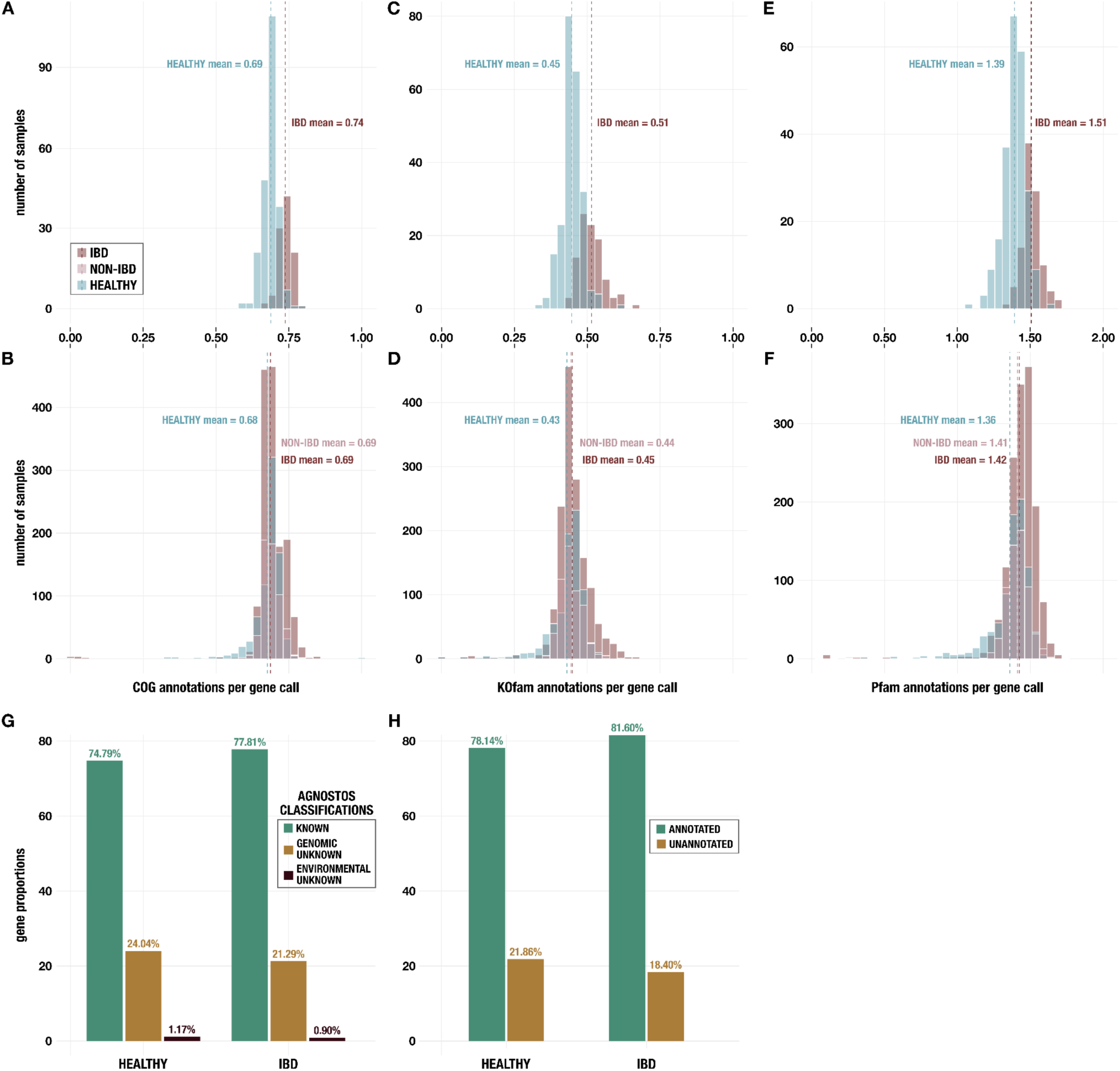
Histograms of annotations per gene call from **A,B)** NCBI COGs; **C,D)** KEGG KOfams; and **E,F)** Pfams. Panels A, C, and E show data for metagenomes in the subset of 330 deeply-sequenced samples from healthy people and people with IBD, and panels B, D, and F show data for all 2,893 samples including those from non-IBD controls. **G)** Proportion of genes with each classification from AGNOSTOS (Vanni et al. 2022) in the subset of 330 deeply-sequenced samples. **H)** Proportion of genes with at least one annotation from KEGG KOfams (Aramaki et al. 2020), NCBI COGs (Galperin et al. 2015), or Pfams (Mistry et al. 2021) (green) and proportion without any annotation (brown) in the subset of 330 deeply-sequenced samples.

**Supplementary Figure 5.**
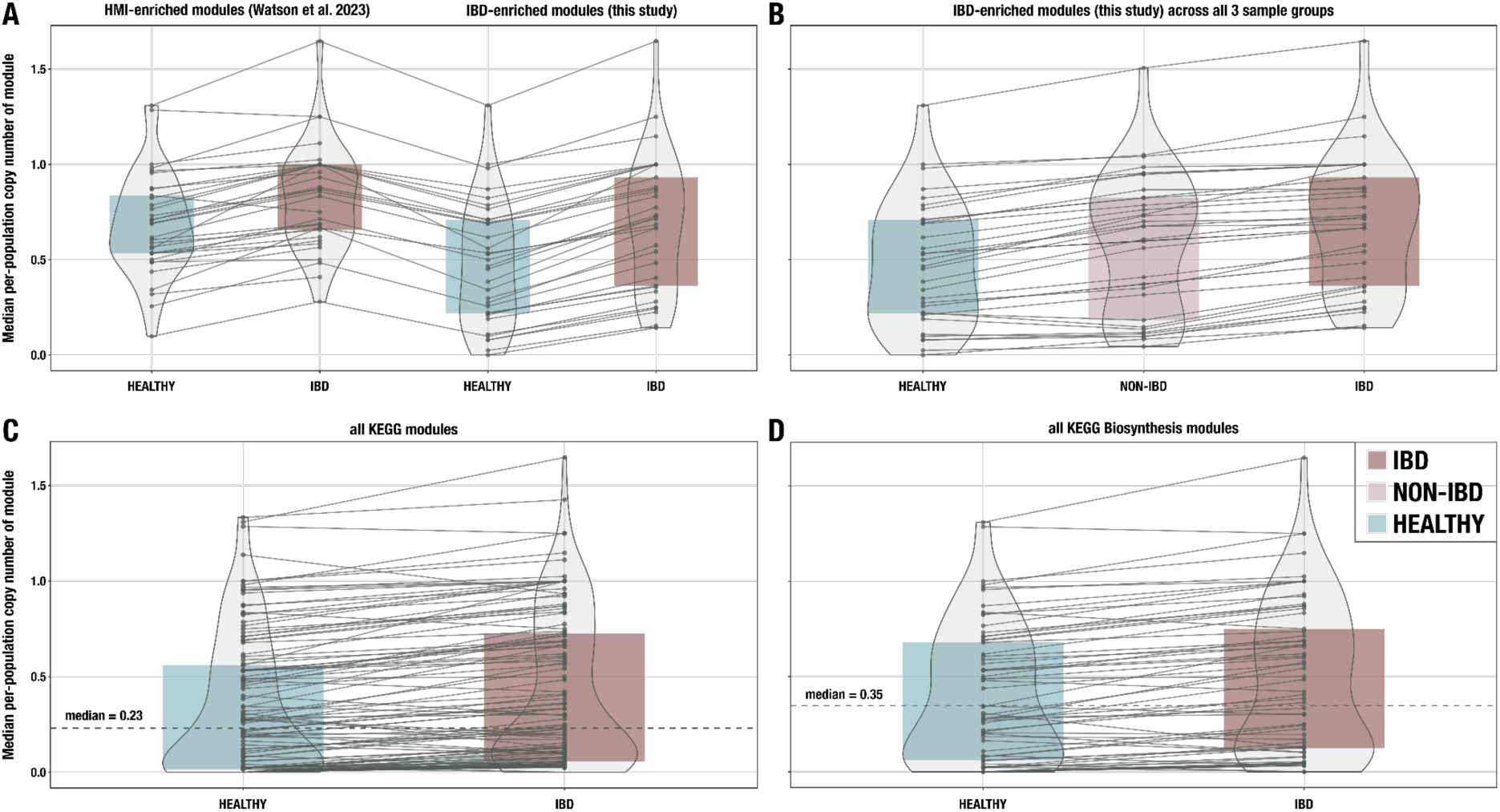
Additional boxplots of median per-population copy number for various subsets of metabolic pathways and metagenome samples. A) 33 modules enriched in HMI populations from (Watson et al. 2023) compared to the 33 IBD-enriched modules from this study, with medians computed in the set of deeply-sequenced healthy (n = 229) and IBD (n = 101) samples. **B)** The 33 IBD-enriched modules from this study, with medians computed in the set of deeply-sequenced healthy (n = 229), non-IBD (n = 78), and IBD (n = 101) samples. **C)** All KEGG modules (n = 117) with non-zero copy number in at least one sample, with medians computed in the set of deeply-sequenced healthy (n = 229) and IBD (n = 101) samples. **D)** All biosynthesis modules (n = 88) from the KEGG MODULE database, with medians computed in the set of deeply-sequenced healthy (n = 229) and IBD (n = 101) samples. Where applicable, dashed lines indicate the overall median for all modules, and solid lines connect the points for the same module in each sample group. The IBD sample group is highlighted in red, the NONIBD group in pink, and the HEALTHY group in blue.

**Supplementary Figure 6.**
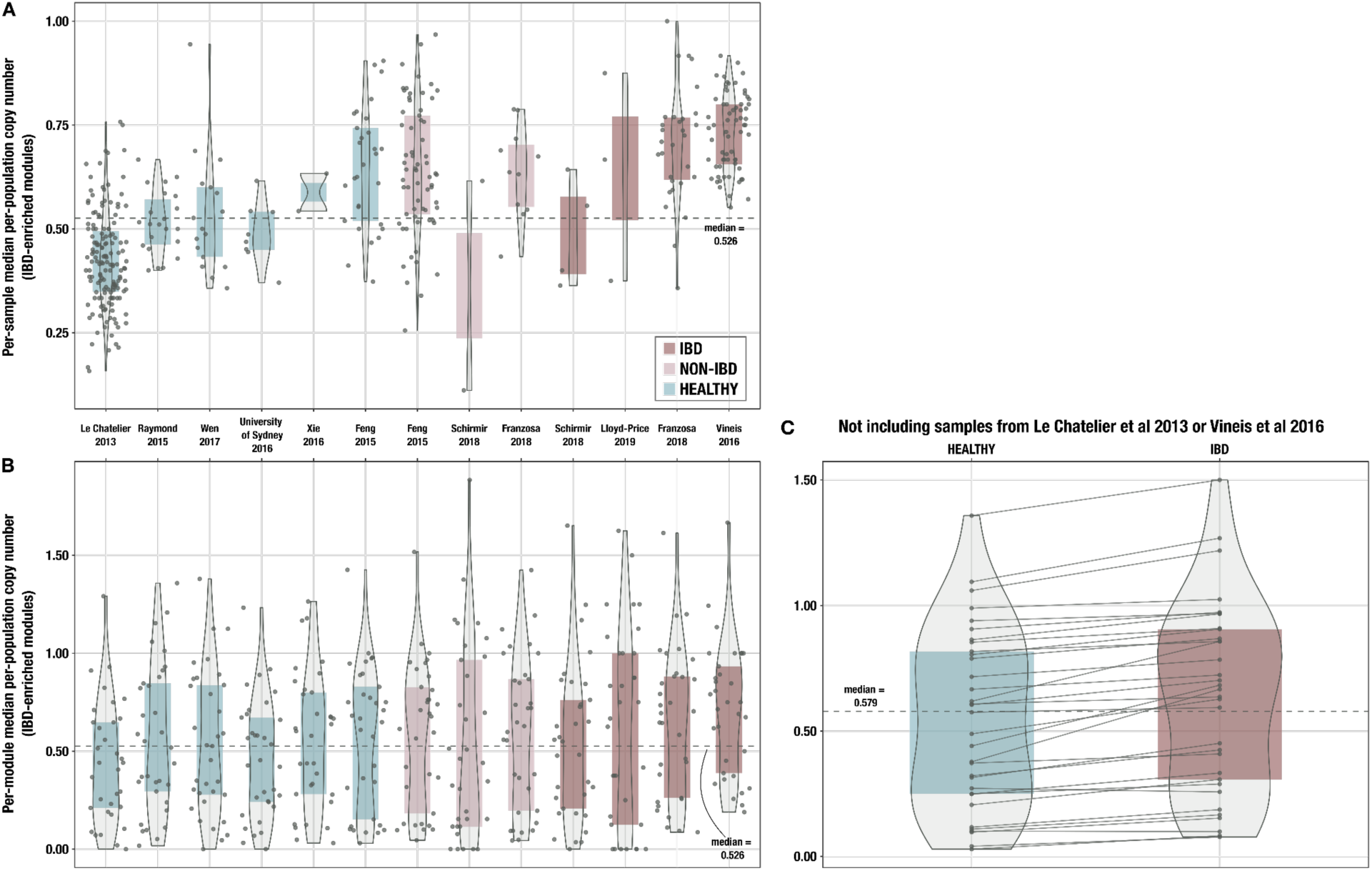
Boxplots of median per-population copy number of 33 IBD-enriched modules for samples from each individual cohort, **A)** with medians computed within each sample (ie, one point per sample) and **B)** with medians computed for each IBD-enriched module (ie, one point per module). The x-axis indicates study of origin. **C)** Boxplots of median per-population copy number of 33 IBD-enriched modules for the 115 samples in the deeply-sequenced set that are not from (Le Chatelier et al. 2013) or (Vineis et al. 2016). The dashed line indicates the overall median for all 33 modules, and solid lines connect the points for the same module in each sample group.

**Supplementary Figure 7.**
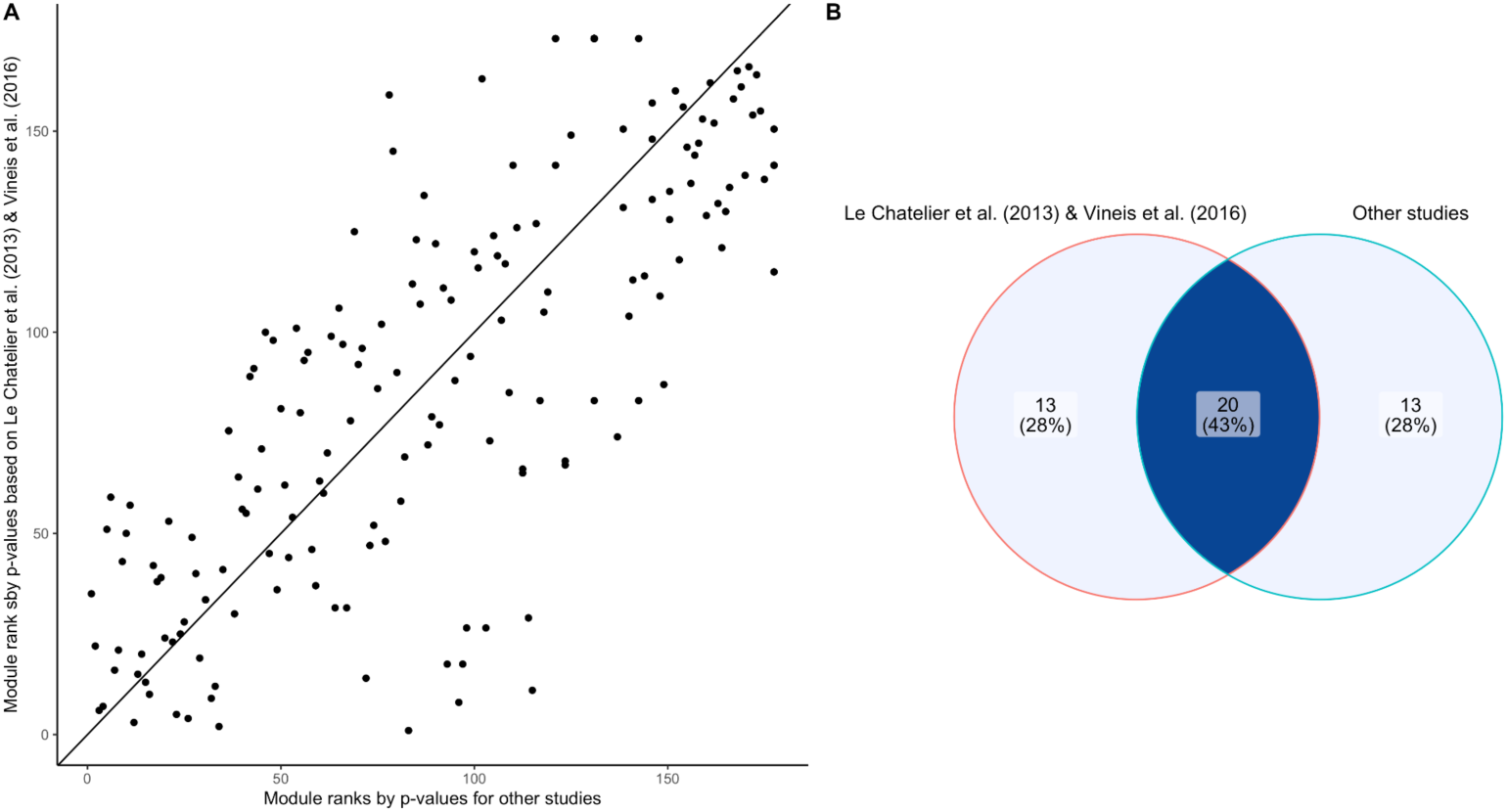
Assessing batch effect of the IBD-enrichment study. **A)** Scatter plot comparing the module ranks of Wilcoxon-Mann-Whitney p-values comparing IBD and healthy subjects on (Le Chatelier et al. 2013) and (Vineis et al. 2016) (y axis) and the rest of our dataset (x axis). **B)** Venn diagram displaying the overlap of IBD-enriched modules identified by the 33 smallest p-values in (Le Chatelier et al. 2013) and (Vineis et al. 2016) and the rest of our dataset. There is good agreement (20 out of 33) between the two sets of modules, indicating generalizability of the signals across studies used in our sample set.

## Supplemental Tables

Supplementary Tables and our Supplementary Files can be accessed at doi:10.6084/m9.figshare.22679080.

**Table 1: Samples and cohorts used in this study.** a) Description of studies/cohorts providing publicly-available gut metagenomes from healthy people, non-IBD controls, and people with IBD. For each study, we note the sample groups it contributes metagenomes to; whether or not those samples were sufficiently deeply sequenced to be included in the main analyses; the country of origin of the samples; the sample type (fecal metagenome or ileal pouch luminal aspirate); the number of samples it contributes to each group before and after applying the sequencing depth threshold; and cohort details/exclusions as described within the study. b) Description of 408 samples included in the primary analyses of this manuscript (ie, those with sufficient sequencing depth of >= 25 million reads), including their associated diagnosis (ulcerative colitis (UC), Crohn’s disease (CD), non-IBD, healthy, colorectal cancer with adenoma (CRC_ADENOMA), or colorectal cancer with carcinoma (CRC_CARCINOMA)); study of origin; sample group; sequencing depth; and number of microbial populations estimated to be represented within the metagenome. c) Description of all samples initially considered and their SRA accession numbers. d) The number of gene calls and the number/proportion of annotations per gene call for KOfams, COGs, and Pfams in each sample. e) The number of genes with at least one functional annotation and the number of tRNAs in each sample from the subset of deeply-sequenced samples. f) Description of the 57 antibiotic time-series gut metagenomes from (Palleja et al. 2018) used for classifier testing, including SRA accession number; sampling day in the time series; sequencing depth; and estimated numbers of microbial populations represented in the sample.

**Table 2: Metabolism data in metagenomes.** a) Description of the 33 KEGG modules enriched in IBD samples, including: module name, KEGG categorization, and definition; their median per-population copy numbers (PPCN) in the healthy sample group and IBD sample group; the p-value, FDR-adjusted p-value, and W statistic from the per-module Wilcoxon Rank Sum test used to determine enrichment in IBD; the difference between its median PPCN in IBD samples and median PPCN in healthy samples (‘effect size’); the fraction of samples in which the module occurs with non-zero copy number; whether the module is also enriched in the HMI populations analyzed in (Watson et al. 2023); the number of total enzymes in the module; the number of total compounds in the modules; and the numbers and proportions of shared enzymes or compounds between this module and the other IBD-enriched modules. b) Description of all 179 KEGG modules with non-zero copy number in at least one metagenome. Most of the columns match the corresponding column in sheet (b) with the exception of the ‘enrichment status’ column, which indicates whether the module was found to be enriched in the IBD samples in this study (‘IBD_ENRICHED’), in the high-metabolic independence genomes in (Watson et al. 2023) (‘HMI_ENRICHED’), in both (‘HMI_AND_IBD’), or in neither (‘OTHER’). c) Matrix of stepwise copy number of each module in each deeply-sequenced gut metagenome. d) Per-population copy number of each module in each deeply-sequenced gut metagenome in the IBD, non-IBD and healthy sample groups. e) Per-population copy number of each module in each antibiotic time-series sample from (Palleja et al. 2018).

**Table 3: GTDB genome data.** a) List of 338 GTDB representative genomes identified as gut microbes, their taxonomy, metabolic independence score, classification as high metabolic independence (‘HMI’) or not (‘non-HMI’), genome length in base pairs, and number of gene calls. b) Matrix of stepwise completeness of each module in each genome. c) Matrix of genome detection in each deeply-sequenced gut metagenome in the IBD, non-IBD, and healthy sample groups. d) Percent abundance of each genome in each deeply-sequenced gut metagenome. e) Per-genome proportion of samples from each sample group that the genome is detected in using a threshold of 50% (ie, at least half of the genome sequence is covered by at least one sequencing read in a given sample). f) Per-sample proportion of detected genomes that are classified as HMI. g) Average completion of each IBD-enriched module within the HMI genome group and the non-HMI genome group, as well as the difference between these values.

**Table 4: Metagenome classifier information.** a) Details and performance of previously-published classifiers for IBD and IBD subtypes. For each classifier, we summarize the cohort details as described by the study; the size of training datasets and validation datasets (if any); the type(s) of samples, data, and extracted features used for classification; the target classes (that is, what the samples were being classified as); the classifier type and training/validation strategy; and the performance metrics as reported by the study. b) Classification of each (Palleja et al. 2018) metagenome by our logistic regression model trained for distinguishing IBD vs healthy samples on the basis of PPCN data for IBD-enriched modules. This table describes whether the sample was classified as healthy (‘HEALTHY’) or stressed (‘IBD’, which we consider to be equivalent to an identification of gut stress), and also whether the sample had low sequencing depth (< 25 million reads) or not. c) Summary of the performance of our metagenome classifier across different training/validation strategies using the IBD and healthy metagenome samples. It also includes the details of our final classifier trained on all 330 samples, though performance data is not available for this model since there were no IBD/healthy samples left for validation – however, see manuscript for its performance on the (Palleja et al. 2018) antibiotic time-series dataset. The subsequent sheets include per-fold data and performance information for each train-test strategy: d) random split cross-validation (25-fold) on PPCN data; e) leave-two-studies-out cross-validation (24-fold); and f) (10-fold) cross-validation leaving out samples from the two dominating studies in our dataset, (Le Chatelier et al 2013) and (Vineis et al 2016).

**Table 5: Details of available software for metabolism estimation.** For each tool (including the one published in this study), we summarize: the software category (based upon the tool’s architecture and mode of use); its metabolism reconstruction strategy (whether it is a pathway prediction tool or a modeling tool or both); the data source(s) it uses for enzyme and metabolic pathway information; how it calculates pathway completeness or generates models (depending on reconstruction strategy); what input and output types it accepts/generates; any additional capabilities as advertised by the tool’s publication; whether or not the tool is open-source; the program type; and what language(s) it is developed in (if known). The reference publication and code repository or webpage for each tool is also included.

**Table 6: Data from validation of the PPCN approach with simulated metagenomic data.** a) Distribution (mean and standard deviation) of PPCN and PPCN error (computed relative to either average genomic completeness or average genomic copy number) across each validation test case, as well as the proportion of correct, off-by-one, and off-by-two community size estimates in each test case. b) Spearman’s correlation test results between sample parameters (genome size, community size, and diversity level) and important values computed in our approach (PPCN, PPCN accuracy metrics, and accuracy of community size estimates). Negative values are shown in red and nonsignificant p-values are highlighted in blue. c) Normalized and rank-ordered relative abundance data from the top 20 most abundant microbial populations in healthy gut metagenomes that we used to recreate a ‘typical’ relative abundance curve for our simulated metagenomes, based upon data from <u>(Beghini et al. 2021)</u>. Each initial column provides the data from a single sample, and the final four columns describe: the average relative abundance at each rank-order; the averages when scaled such that the minimum abundance is 1; the corresponding (integer) coverage values for each scaled average relative abundance value; and the coverage values when increased by 20x for sufficient sequencing depth for assembly.

