## Supplementary File 1 for "Microbes with higher metabolic independence are enriched in human gut microbiomes under stress"

[1]Biophysical Sciences Program, The University of Chicago, Chicago, IL 60637, USA; [2]Department of Medicine, The University of Chicago, Chicago, IL 60637, USA; [3]Data Science Institute and Department of Biomedical Data Science, Stanford University, Stanford, CA, 94305, USA; [4]Committee on Microbiology, The University of Chicago, Chicago, IL 60637, USA; [5]MARUM Center for Marine Environmental Sciences, University of Bremen, Bremen, Germany; [6]Department of Biostatistics, University of Washington, Seattle, WA, 98195, USA; [7]Toyota Technological Institute at Chicago, Chicago, IL 60605, USA; [8]Lundbeck Foundation GeoGenetics Centre, GLOBE Institute, University of Copenhagen, Copenhagen, Denmark; [9]Institute for Chemistry and Biology of the Marine Environment, University of Oldenburg, Oldenburg, Germany; [10]Marine 'Omics Bridging Group, Max Planck Institute for Marine Microbiology, 28359 Bremen, Germany; [11]Alfred Wegener Institute for Polar and Marine Research, Bremerhaven, Germany; [12]Helmholtz Institute for Functional Marine Biodiversity, 26129, Oldenburg, Germany

#### Low sequencing depth results in poor characterization of community richness in assembled metagenomes

Within our dataset, we observed a correlation between the estimated number of distinct populations in assembled sequences and sequencing depth, i.e. the number of short reads generated from a given sample. In shallow sequencing of metagenomes, short reads may not cover the entirety of population genomes, thus decreasing the rate of recovery of single-copy core genes (SCGs) in the assembly and resulting in an underestimation of the number of populations present. Indeed, the linear relationship between sequencing depth and the number of observed microbial populations we observe in lower depths of sequencing (Supplementary Figure 1) starts to plateau once the sequencing depth exceeds approximately 25 million reads, suggesting that our strategy to estimate the number of distinct microbial populations within these samples serves as a good approximation of the true number of genomes only at relatively higher depths of sequencing. Since an incomplete recovery of population genomes in metagenomic samples also interferes with a meaningful quantification of metabolic potential in a given sample, we set a minimum sequencing depth threshold of 25 million sequencing reads (Supplementary Figure 1). A set of 408 samples (101 IBD, 229 healthy, and 78 non-IBD) from 10 different studies passed our quality threshold to be utilized for further analysis.

Low sequencing depth disproportionately affects IBD metagenomes, thereby reducing our ability to effectively study this disease model in comparison with healthy controls. It also disproportionately affects some studies over others, which could allow cohort or study-specific effects to influence the differential signal between the groups. However, we concluded that the benefits of stringent thresholding outweigh the potential complications arising from imbalanced cohort sizes in our sample subset.

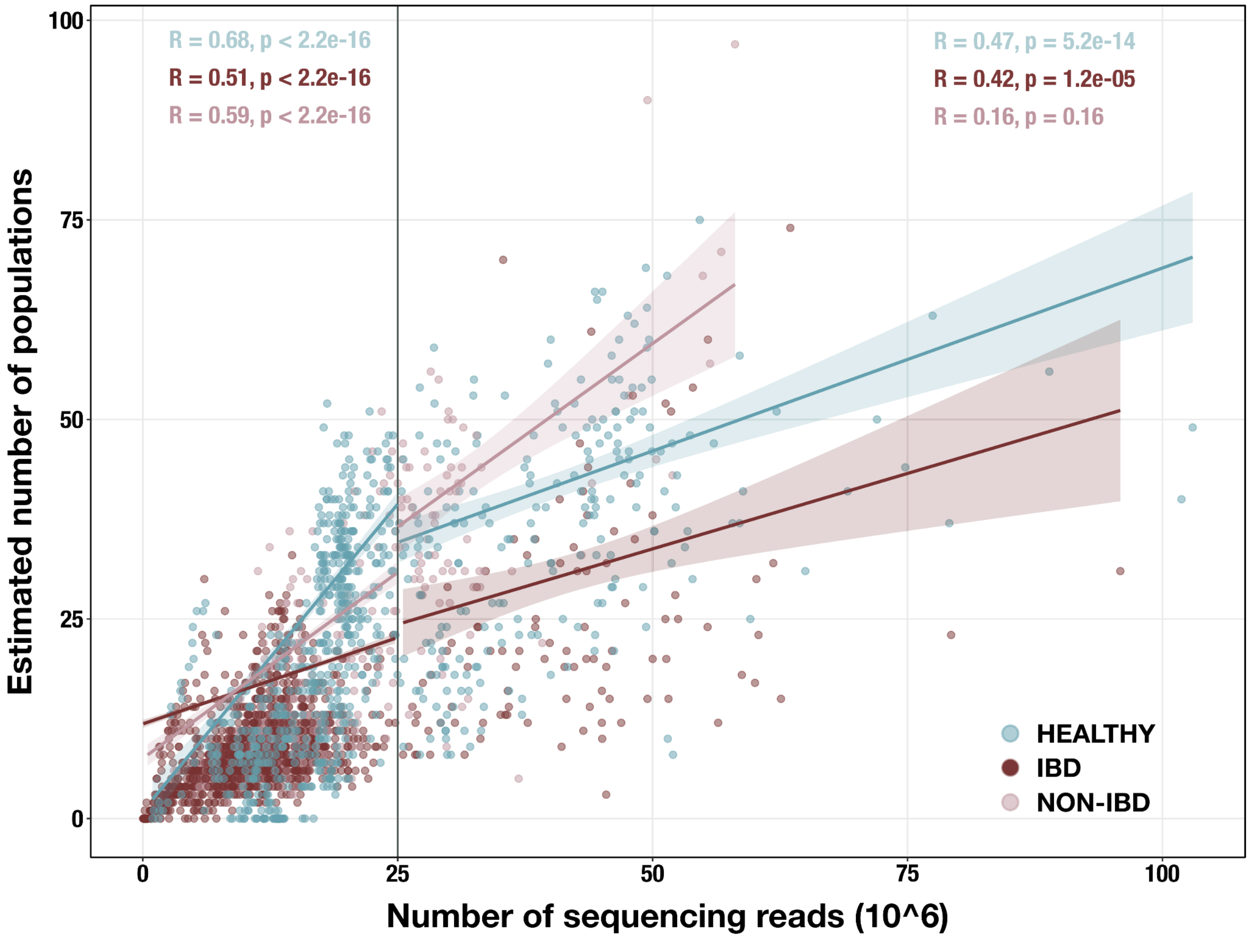


**Supplementary Figure 1. Scatterplot of sequencing depth vs estimated number of microbial populations in each of 2,893 stool metagenomes.** Sequencing depth is represented by the number of R1 reads, except for (Vineis et al. 2016) samples, in which case it is the number of merged paired-end reads. The vertical line indicates our sequencing depth threshold of 25 million reads. Per-group Spearman's correlation coefficients and p-values are shown for the subset of samples with depth < 25 million reads (top left) and for the subset with depth >= 25 million reads (top right). Regression lines are shown for each group in each subset, with standard error indicated by the colored background.

### Technical details of metabolism estimation in anvi'o

This section describes technical details of the program `anvi-estimate-metabolism`, which is the main program in the metabolism reconstruction framework in anvi'o (Supplementary Figure 2a). Documentation for this program, including an extended and more up-to-date version of these technical details, can be found at https://anvio.org/m/anvi-estimate-metabolism.

A

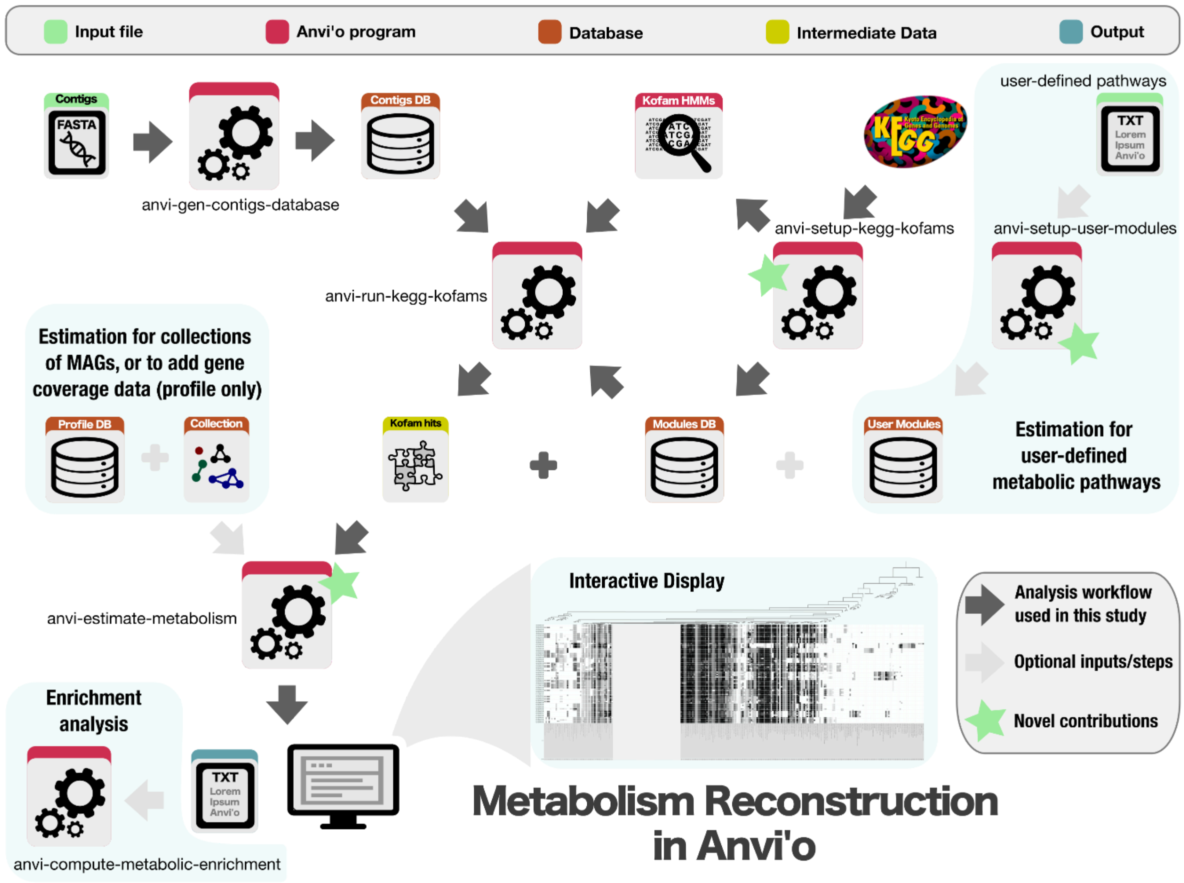


B

#### Metabolic Pathway

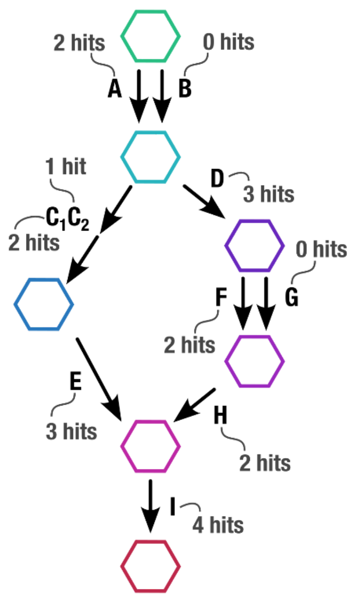


C

(A or B) and (($C_1$ + $C_2$ and E) or (D and (F or G) and H) and I

D

| STEPS | BOOLEAN EXP. | PRESENT? | ARITHMETIC EXP. | COPY # |
|---|---|---|---|---|
| A or B | T or F | Yes | 2 + 0 | 2 |
| ($C_1$ + $C_2$ and E) or (D and (F or G) and H) | (T and T and T) or (T and (F or T) and T) | Yes | min(2, 1, 3) + min(3, (0 + 2), 2) | 3 |
| I | T | Yes | 4 | 4 |

E

Stepwise completeness: 3/3 = 1.0

Stepwise copy number: min(2,3,4) = 2

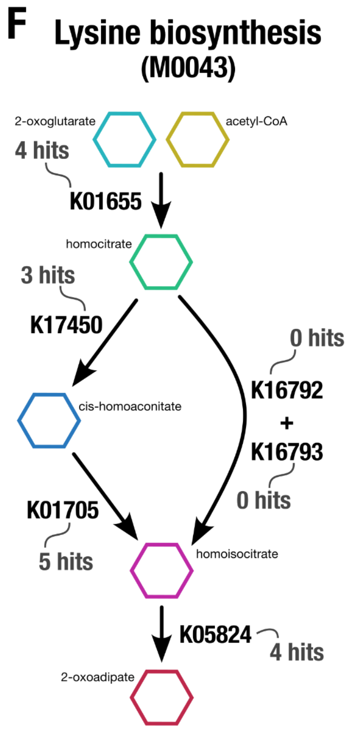


**G**

**K01655 and (K17450 and K01705 or K16792 + K16793) and K05824**

**H**

| STEPS | BOOLEAN EXP. | PRESENT? | ARITHMETIC EXP. | COPY # |
|---|---|---|---|---|
| K01655 | T | Yes | 4 | 4 |
| K17450 and K01705 or K16792 + K16793 | T and T or F and F | Yes | min(3, 5) + min(0,0) | 3 |
| K05824 | T | Yes | 4 | 4 |

**I**

**Stepwise completeness: 3/3 = 1.0**

**Stepwise copy number: min(4,3,4) = 3**

**Supplementary Figure 2. Technical details of the metabolism reconstruction software framework in anvi'o. A)** Workflow of metabolism reconstruction programs and their inputs/outputs. Dark arrows indicate the primary analysis path utilized in this study. Blue background indicates optional features in the framework. A demonstration of completeness score and copy number calculations for metabolic pathways (performed by the program `anvi-estimate-metabolism` is shown using example enzyme annotation data in panels B – E (for a theoretical pathway) and F – I (for a real pathway). **B)** Theoretical metabolic pathway, where hexagons represent metabolites, arrows represent chemical reactions, letters represent enzymes (subscripts indicate enzyme components), and the example number of gene annotation hits for each enzyme is written in gray. **C)** The definition of the theoretical pathway from panel B, written in terms of the required enzymes. **D)** Table showing the major steps in the pathway and example calculations for step presence and copy number. Step presence is calculated by evaluating a boolean expression created from the step definition in which enzymes with > 0 hits are replaced with True (T) and the others with False (F). Step copy number is calculated by evaluating the corresponding arithmetic expression in which the enzymes are replaced with their annotation counts. **E)** Final calculations of completeness score (fraction of present steps) and copy number for the theoretical metabolic pathway. **F – I)** Same as panels B – E, but for KEGG module M00043. A high-resolution version of this figure is available at https://doi.org/10.6084/m9.figshare.22851173.

#### Summary of program usage

The program `anvi-estimate-metabolism` predicts the metabolic capabilities of organisms based on their genetic content. It relies upon enzyme annotations and metabolism information from KEGG, specifically using metabolic modules from the KEGG MODULE (Kanehisa et al. 2023) database, which are defined in terms of KEGG Orthologs (KOs) that can be annotated via the KOfam database of hidden Markov model (HMM) profiles (Aramaki et al. 2020). It can also work with user-defined metabolic pathways, as described in the documentation page https://anvio.org/m/user-modules-data. In our analysis, we used the program `anvi-run-kegg-kofams` to annotate the KOs used for downstream metabolism estimation; this software implements a heuristic to obtain more accurate and comprehensive annotation results than other contemporary tools for KOfam annotation, as described in (Kananen et al. 2024).

The program `anvi-estimate-metabolism` determines which enzymes are annotated in an input sample and uses these functions to compute the completeness and copy number of each metabolic module within the sample. Input samples can be individual genomes, binned or unbinned metagenomes, or ad-hoc lists of enzyme accessions. The output of `anvi-estimate-metabolism` is one or more tabular text files detailing the completeness and copy number scores per module as well as (customizable) information such as pathway metadata; shared/unique enzymes; gene coverage data; and pathway substrates, intermediates, and products. A detailed output description and examples can be found at https://anvio.org/m/kegg-metabolism/.

#### Module definitions and interpretation strategies

Metabolic pathways are defined by the enzymes responsible for each reaction in the pathway, using the convention established by the KEGG MODULE database. In these definitions, commas separate alternative enzymes that can catalyze the same reaction, spaces separate subsequent reactions, plus signs indicate essential components of enzyme complexes, minus signs indicate non-essential components of complexes, and parentheses indicate the order of operations. These definitions can also be written in terms of the logical relationships between reactions, such that spaces and plus signs are converted into ‘AND’ relationships and commas are converted into ‘OR’ relationships (Supplementary Figure 2b-c and f-g).

`anvi-estimate-metabolism` has two strategies for interpreting module definition strings that treat alternative enzymes and pathway branches differently. One is the ‘pathwise’ strategy, which considers all possible combinations of enzymes. In this method, each alternative set of enzymes that could be used together to catalyze every reaction in the metabolic pathway is called a ‘path’ through the module. The program computes completeness and copy number metrics for each path separately, and then identifies the most complete path(s) as the most biologically-relevant representative of the module as a whole. Alternatively, with the ‘stepwise’ strategy the module definition is parsed into high-level ‘steps’ that each encompasses a set of alternative enzymes for a particular reaction or branch point. The presence and copy numbers of each step are respectively combined into a completeness score and copy number for the entire module.

#### Calculation of stepwise completeness and copy number

The analyses in this paper rely on the ‘stepwise’ metrics of module completeness and copy number, which are calculated as demonstrated in Supplementary Figure 2d-e and h-i. We divide each module into steps by splitting the definition string on the outermost ‘AND’ relationships (spaces not within parentheses). To determine whether each step is present, we convert the step definition into a Boolean expression in which ‘True’ represents annotated enzymes and ‘False’ represents enzymes without annotations. If the Boolean expression evaluates to ‘True’, then the step is considered present. The module completeness score is the number of present steps divided by the total number of steps. To determine the step copy number, we convert the step definition into an arithmetic expression wherein ‘AND’ relationships become minimum operations and ‘OR’ relationships become addition operations. We take the minimum of all per-step copy numbers obtained by evaluating these arithmetic operations to get the overall module copy number.

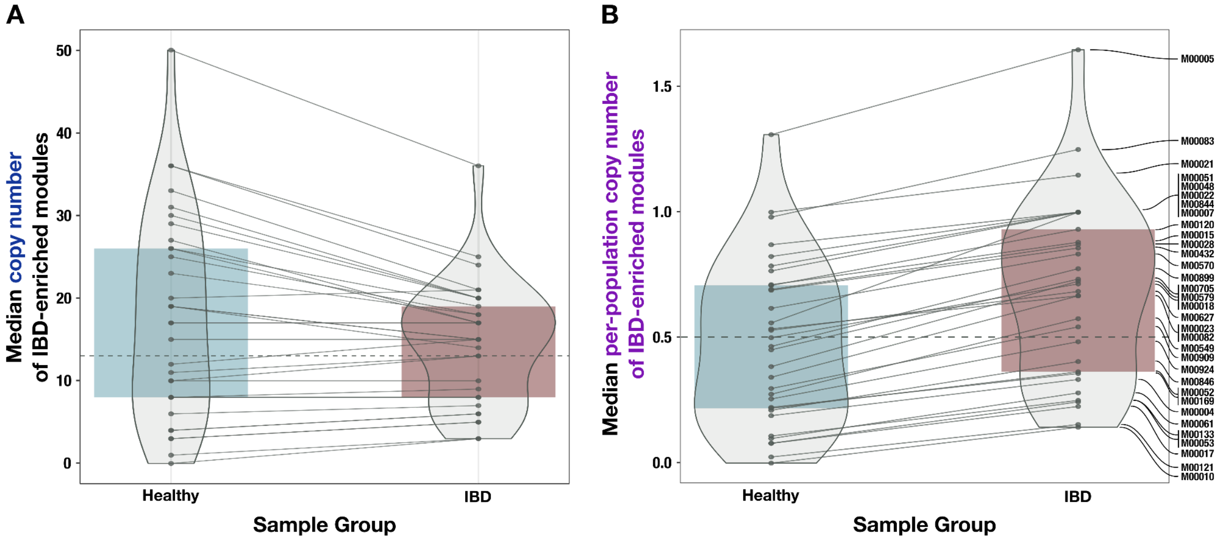


**Supplementary Figure 3. Comparison of unnormalized copy number data and normalized (per-population copy number, or PPCN) data for the IBD-enriched modules. A)** Boxplot of median copy numbers for each module in the healthy samples (blue) and IBD samples (red). **B)** Boxplots of median PPCN for each module in the healthy samples (blue) and IBD samples (red). Lines connect data points for the same module in each plot. The gray dashed line in each plot indicates the overall median value.

#### Differential annotation efficiency between IBD and Healthy samples

We observed that the proportion of predicted genes with functional annotations was markedly less in healthy metagenomes than in IBD samples, for both sequence homology-based annotation methods (NCBI Clusters of Orthologous Groups, or COGs) and annotation with probabilistic models (KEGG KOfams and Pfams) (Supplementary Figure 4, Supplementary Table 1d). One possible interpretation of this that aligns with our metabolic competency hypothesis is that the populations with low metabolic independence (LMI) that thrive in the healthy gut environment are relatively less well-characterized than the HMI populations that are more likely to survive in the stressful conditions of IBD, resulting in an annotation bias against healthy samples. This interpretation is congruent with our observation that most uncharacterized gut microbial genomes from the GTDB, which have temporary code names in place of taxonomic assignments, were identified as non-HMI (Figure 3b).

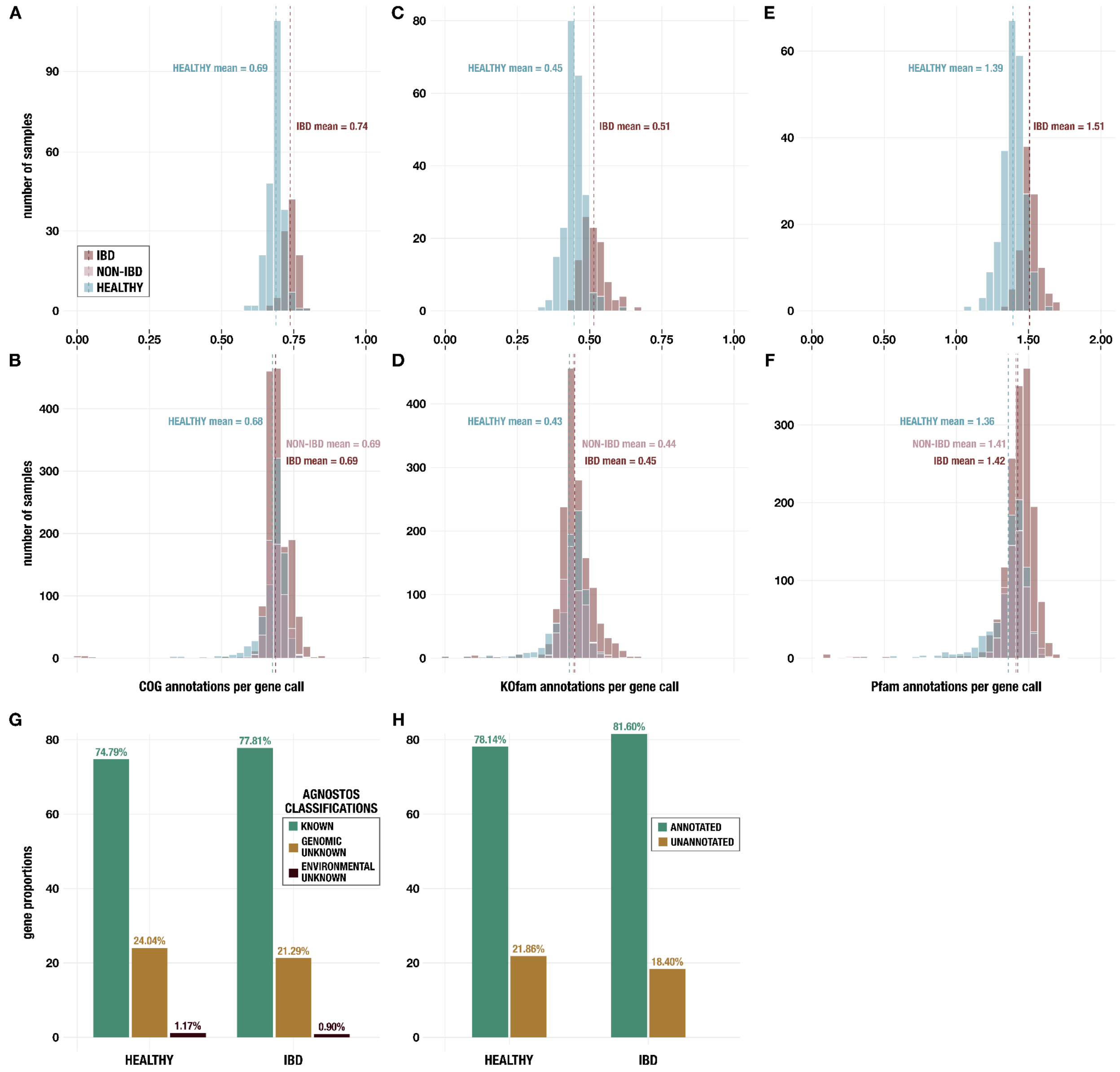


**Supplementary Figure 4. Histograms of annotations per gene call** from **A,B)** NCBI COGs; **C,D)** KEGG KOfams; and **E,F)** Pfams. Panels A, C, and E show data for metagenomes in the subset of 330 deeply-sequenced samples from healthy people and people with IBD, and panels B, D, and F show data for all 2,893 samples including those from non-IBD controls. **G)** Proportion of genes with each classification from AGNOSTOS (Vanni et al. 2022) in the subset of 330 deeply-sequenced samples. **H)** Proportion of genes with at least one annotation from KEGG KOfams (Aramaki et al. 2020), NCBI COGs (Galperin et al. 2015), or Pfams (Mistry et al. 2021) (green) and proportion without any annotation (brown) in the subset of 330 deeply-sequenced samples.

The reduced metabolic capacity of LMI microbes and their resulting reliance on robust community interactions (i.e., cross-feeding) may increase their resistance to cultivation that typically aims to isolate individual populations rather than communities. Microbes auxotrophic for key metabolites rely on metabolic interactions with their surrounding community and current cultivation practices may not sufficiently account for the lack of such interactions. As a result, the available genomes of such populations would be limited to sequences from metagenomic surveys, which are often incomplete and/or composite and are therefore typically not included in efforts to generate models and non-redundant sequence databases for gene annotation (Aramaki et al. 2020; Galperin et al. 2015; Sonnhammer, Eddy, and Durbin 1997). Thus, the reduced proportion of annotated genes in healthy metagenomes may reflect missing annotations due to lack of sufficiently-homologous sequences in state-of-the-art databases. The true reduction in metabolic potential in the healthy sample group may not be as extensive as we have observed in this study.

The discrepancy in annotation efficiency between the healthy and IBD groups disappeared when analyzing all 2,893 samples (Supplementary Figure 4). This suggests that the observed annotation bias does not strongly affect microbial populations that are readily assembled via shallow sequencing – likely, these are populations of high relative abundance in both healthy and IBD samples. Populations of lower abundance, which are less likely to be assembled from shallow metagenomes due to lack of sufficient coverage, are probably also less well-characterized as a result. For this to contribute to fewer annotations per gene in healthy samples would necessitate that healthy samples contain relatively more low-abundance populations than IBD samples. Indeed, this is the case: healthy samples contain an average of 86 detected genomes from our set of GTDB gut microbes, and those genomes have a low average percent abundance of 0.61% across these samples. Non-IBD samples are similar, having an average of 77 detected genomes per-sample with an average percent abundance of 0.79%. IBD samples, meanwhile, contain 30 detected genomes on average, with a higher average percent abundance of 2.24%. Therefore, the lack of characterization of low-abundance populations may be another factor that contributes to the relative reduction in gene annotations in the healthy samples.

To understand the potential origins of the reduced annotation rate in healthy metagenomes, we ran AGNOSTOS (Vanni et al. 2022) to classify known and unknown genes within the healthy and IBD sample groups. AGNOSTOS clusters genes to contextualize them within an extensive reference dataset and then categorizes each gene as ‘known’ (has homology to genes annotated with Pfam domains of known function), ‘genomic unknown’ (has homology to genes in genomic reference databases that do not have known functional domains), or ‘environmental unknown’ (has homology to genes from metagenomes or MAGs that do not have known functional domains). The resulting classifications confirm that healthy metagenomes contain fewer ‘known’ genes than metagenomes in the IBD sample group – the proportion of ‘known’ genes classified by AGNOSTOS is about 3.0% less in the healthy metagenomes than in the IBD sample group, which is similar to the ~3.5% decrease in the proportion of ‘unannotated’ genes observed by simply counting the number of genes with at least one functional annotation (Supplementary Figure 4g-h, Supplementary Table 1e). Furthermore, the majority of the unannotated genes in either sample group were categorized by AGNOSTOS as ‘genomic unknown’ (Supplementary Figure 4g), suggesting that the unannotated sequences are genes without biochemically-characterized functions currently associated with them and are thus legitimately lacking a functional annotation in our analysis, rather than representing distant homologs of known protein families that we failed to annotate. Based upon the classifications, a systematic technical bias is unlikely driving the annotation discrepancy between the sample groups.

Our observations into the annotation efficiency of microbial genes in different contexts, and our speculations regarding the sources of such bias and its implications warrant further investigations.

#### Pathway enrichment without consideration of effect size leads to nonspecific results

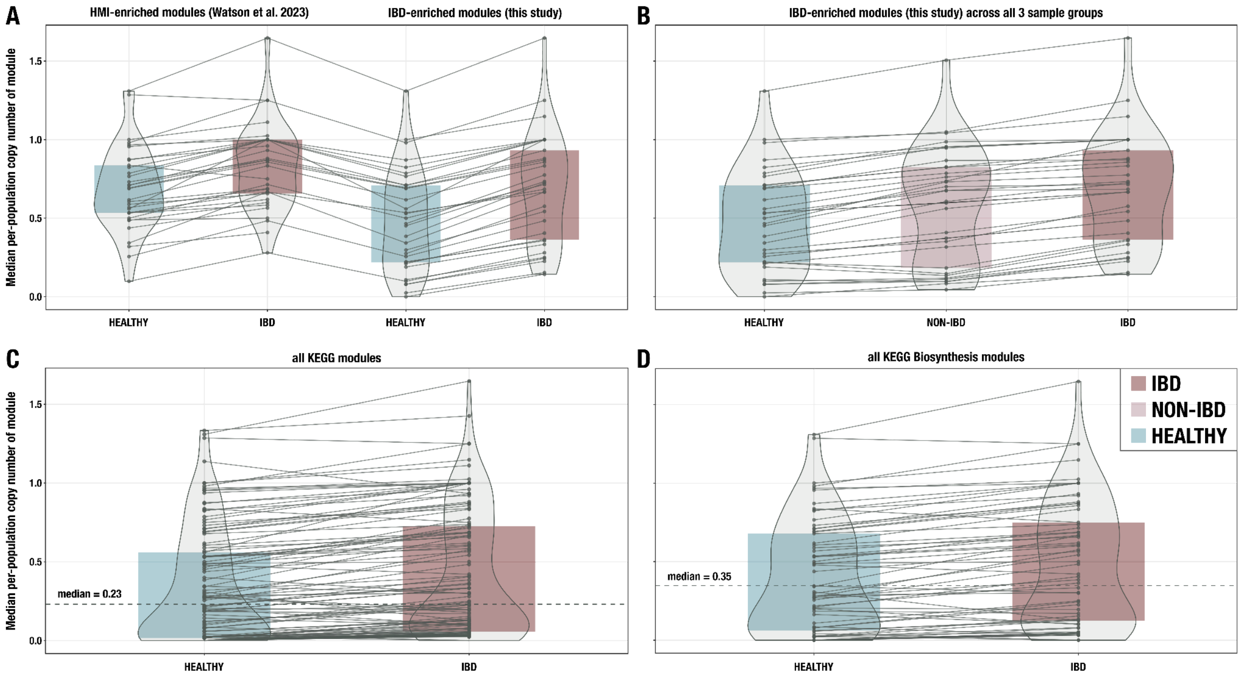


**Supplementary Figure 5. Additional boxplots of median per-population copy number for various subsets of metabolic pathways and metagenome samples. A)** 33 modules enriched in HMI populations from (Watson et al. 2023) compared to the 33 IBD-enriched modules from this study, with medians computed in the set of deeply-sequenced healthy (n = 229) and IBD (n = 101) samples. **B)** The 33 IBD-enriched modules from this study, with medians computed in the set of deeply-sequenced healthy (n = 229), non-IBD (n = 78), and IBD (n = 101) samples. **C)** All KEGG modules (n = 117) with non-zero copy number in at least one sample, with medians computed in the set of deeply-sequenced healthy (n = 229) and IBD (n = 101) samples. **D)** All biosynthesis modules (n = 88) from the KEGG MODULE database, with medians computed in the set of deeply-sequenced healthy (n = 229) and IBD (n = 101) samples. Where applicable, dashed lines indicate the overall median for all modules, and solid lines connect the points for the same module in each sample group. The IBD sample group is highlighted in red, the NON-IBD group in pink, and the HEALTHY group in blue.

Our analyses indicate that the majority of KEGG modules had higher per-population copy number in IBD metagenomes (Figure 2e, Supplementary Table 2b). Indeed, when we examine all modules with non-zero median per-population copy number in at least one group of samples (n = 117), their median normalized copy number is systematically higher in the IBD group than in the healthy group (Supplementary Figure 5c; 98 out of 117 modules have a higher normalized copy number in the IBD group than in the healthy group at 5% FDR-adjusted significance level using a one-sided Wilcoxon test). This result is likely a natural outcome of the differential distribution of HMI and non-HMI genomes in the two sample groups, as seen in our analysis of reference genomes (Figure 3c, 3d), where the overrepresentation of HMI populations with larger genomes that encode many more complete pathways in IBD samples leads to higher per-population copy numbers computed at the metagenome level. The consistent elevation in PPCN of metabolic modules in IBD could also be attributed, at least in part, to the aforementioned functional annotation biases that seem to disproportionally affect the characterization of healthy metagenomes. The lower annotation efficiency in healthy metagenomes could result in partial copies of pathways, which are ignored by our stringent copy number calculation that only counts complete copies. Therefore, to narrow down our results and identify which pathways are *particularly* important for microbial resilience in the IBD gut environment, we considered only those pathways with the largest difference in normalized copy number ('effect size') between the two groups to identify metabolic modules that are truly elevated in IBD metagenomes (see Methods).

With similar considerations we also investigated whether biosynthetic capacity in general was enriched in IBD samples. For this, we expanded our analysis to also consider biosynthesis pathways that did not meet the enrichment criteria we have used for inclusion in the final set of 33 IBD-enriched modules. As expected, we found that the majority of all biosynthesis pathways in the KEGG Module database (n = 88) have significantly higher normalized copy numbers in IBD samples (Supplementary Figure 5d; at a 5% FDR-adjusted significance level, 62 out of 88 (70%) biosynthesis pathways have a higher normalized copy number using a one-sided Wilcoxon test). This analysis also showed a similar increase for non-biosynthetic pathways: 63 out of 91 (69%) non-biosynthetic pathways showed significant increase in IBD samples (two-sample test for equality of proportion: 0.88). Overall, these data indicate that without the consideration of effect size, both biosynthetic and non-biosynthetic capacity appear to be increased in the IBD gut microbiome. In contrast, maintenance of a higher metabolic capacity for the biosynthesis of essential nutrients emerges as an important factor for microbial resilience in IBD through a strict enrichment criteria in addition to statistical significance scores calculated for differential abundance.

### A review of HMI-associated modules in the context of gut microbiome literature

The 33 pathways that are enriched in microbial communities associated with individuals with IBD likely provide competencies that are critical for survival in the stressed gut environment. In this section, we offer a review of IBD-enriched modules with existing gut microbiome scientific literature.

#### Amino acid pathways

Of the eight proteinogenic amino acids that can be synthesized with IBD-enriched pathways, **leucine, tryptophan, threonine, isoleucine,** and **methionine** are essential amino acids for humans (Lopez and Mohiuddin 2023), while cysteine and **arginine** are semi-essential (Rehman et al. 2020; Tong and Barbul 2004). Furthermore, leucine, tryptophan, isoleucine, and cysteine can reduce oxidative stress for intestinal epithelial cells (Katayama and Mine 2007), which may be beneficial given the increased gastrointestinal oxygen levels associated with IBD (Rigottier-Gois 2013). Several of these amino acids have been analyzed for their potential therapeutic effects in IBD (Yulan Liu, Wang, and Hu 2017). Regardless, it is unknown if the depleted gut microbiome in IBD would produce these amino acids in sufficient quantity to promote health benefits to the host, especially considering that the microbes themselves require these molecules for protein production and as energy sources – for instance, in proteolytic fermentation (L. Wu et al. 2021; Lin et al. 2017).

The **Shikimate pathway,** which converts phosphoenolpyruvate (PEP) and erythrose 4-phosphate (E4P) to chorismate, is a prerequisite for **tryptophan biosynthesis**. This pathway is only present in microorganisms and plants, and it also produces intermediates for other metabolic pathways such as quinate degradation and antibiotic synthesis (Herrmann and Weaver 1999). A recent analysis of paired fecal metagenomes and metatranscriptomes from the Human Microbiome Project using reference genome-based functional inference demonstrated that the Shikimate pathway is typically incomplete in gut microbes and only transcriptionally active in a few, suggesting that most gut microbes are auxotrophic for aromatic amino acids and therefore rely on dietary sources and potentially cross-feeding to obtain these molecules or their precursors (Mesnage and Antoniou 2020). This offers a potential explanation for the enrichment of the Shikimate and tryptophan biosynthesis pathways in the IBD gut microbiome, where a depleted community may restrict the availability of cross-fed metabolites. Indeed, a tryptophan-deficient diet alters the composition of the gut microbiota in aged mice (Yusufu et al. 2021), providing auxiliary evidence that the loss of this amino acid impacts microbial survival. Furthermore, the serum levels of tryptophan are reduced in individuals with IBD due to high host metabolism rates (Nikolaus et al. 2017), which may exacerbate the lack of bioavailable tryptophan for gut microbes. On the host side, tryptophan and its derivatives influence a number of physiological processes, though it is unclear how much the microbial production of tryptophan contributes to these effects (Agus, Planchais, and Sokol 2018). Nevertheless, the lack of tryptophan appears to worsen intestinal inflammation while supplementation can attenuate it (Kim et al. 2010; Hashimoto et al. 2012).

**Cysteine biosynthesis** was previously found to be enriched in the IBD gut microbiome based on reference genome analysis of 16S ribosomal RNA gene amplicons (Morgan et al. 2012), in which the authors propose that cysteine metabolism could be important to microbial management of oxidative stress via the production of glutathione, which is protective against reactive oxygen species (Sherrill and Fahey 1998; Tepe, Shimko, and Duran 2006), from cysteine and glutamate. Cysteine can also be converted into hydrogen sulfide ($H_2S$) by host colonocytes and some intestinal microbes. Though $H_2S$ produced by colonocytes can help support their energy production, excess microbially-derived $H_2S$ in the lumen is a risk factor for gut mucosal inflammation and $H_2S$ may play a role in colorectal carcinogenesis (Blachier, Beaumont, and Kim 2019). Interestingly, cysteine biosynthesis is also enriched in the gut microbiomes of postmenopausal women, where it is thought to contribute to elevated homocysteine levels and therefore to increased risk of cardiovascular disease (Zhao et al. 2019).

**Leucine** and **isoleucine**, as branched-chain amino acids (BCAAs), are important nutrients and signaling molecules in humans (Gojda and Cahova 2021). Gut microbial synthesis of these compounds does contribute to human BCAA pools, as evidenced by experiments with heavy isotope labeling and correlations between serum and fecal BCAA levels (Metges et al. 1999; Dhakan et al. 2019). The extent of this exchange has not been characterized in individuals with IBD. However, a study of individuals receiving anti-integrin therapy for Crohn's disease demonstrated that pathways for biosynthesis of L-isoleucine and arginine were enriched at baseline in the gut microbiomes of responders to the therapy (Ananthakrishnan et al. 2017). In a longitudinal study of mice, biosynthesis pathways for leucine and proline were more abundant in animals modeling IBD (Sharpton et al. 2017).

**Threonine** and **proline** are both important components of intestinal mucins (Johansson and Hansson 2016; Faure et al. 2005) and thus contribute to mucosal barrier integrity, which is typically impaired in IBD (Johansson et al. 2010). For instance, threonine, proline, and cysteine supplementation has been shown to reduce symptoms and restore lactobacilli and bifidobacteria counts in rats with DSS-induced inflammation (Sprong, Schonewille, and van der Meer 2010; Faure et al. 2006). The latter observation suggests the importance of an external source of these three amino acids to the fitness of the lactobacilli and bifidobacterial populations and thereby supports the idea that they are community metabolites.

We also found **methionine** biosynthesis to be enriched in the IBD gut microbiome. In individuals with quiescent inflammatory bowel disease, reduced serum levels of methionine, proline, and tryptophan are correlated with changes in the gut microbiome that are associated with increased symptoms of fatigue (Borren et al. 2021), demonstrating a putative link between methionine bioavailability, microbial abundances, and host wellbeing. Indeed, L-methionine supplementation in piglets results in improved mucosal integrity and villus architecture (Chen et al. 2014), and the activated form of methionine, S-adenosylmethionine, can reverse colon lesions and cytoskeletal damage in intestinal cells in DSS-treated mice (Oz et al. 2005). Yet reducing methionine in high-fat diets given to mice was shown to improve intestinal barrier function, reduce inflammation, and increase the abundance of short-chain fatty acid-producing microbes (Y. Yang et al. 2019), so the net impact of methionine on host health and microbial fitness remains unclear.

**Arginine** has been well-studied in the context of inflammatory bowel disease. It has been shown to reduce cytokine production, promote intestinal healing and improve intestinal barrier function in DSS-treated mice, perhaps by enhancing production of nitric oxide (NO) (Coburn et al. 2012; Gobert et al. 2004; Singh et al. 2019). NO is a free radical that has been implicated in regulating mucosal barrier integrity, gastrointestinal motility, and protection against oxidative stress, though overproduction of this compound can have detrimental effects (Kolios, Valatas, and Ward 2004; Walker et al. 2018). Biosynthesis of ornithine, which is both a precursor and a derivative of arginine, was enriched in the IBD gut microbiome as well, in agreement with another study that reported an increase in ornithine biosynthesis in the gut microbiome of individuals with active ulcerative colitis (Hellmann et al. 2023). Finally, polyamines – which are derived from arginine and were also represented in the enriched pathways – promote intestinal barrier function (L. Liu et al. 2009); for instance, by regulating the growth of intestinal epithelial cells (McCormack and Johnson 1991).

#### Carbohydrate pathways

Three KEGG modules describing the **pentose phosphate pathway** were enriched in IBD samples - the entire pentose phosphate cycle (M00004), the oxidative phase (M00006), and the non-oxidative phase (M00007). We removed the oxidative phase (M00006) from our set of IBD-enriched modules because it was an exact copy of the initial steps in M00004; however, we kept the non-oxidative phase (M00007) in our set because it is defined using slightly different enzymes than the non-oxidative portion of M00004. M00007 is defined in four steps and utilizes a ribulose-phosphate 3-epimerase and a ribose 5-phosphate isomerase in the last two steps, while the non-oxidative phase in M00004 is defined in three steps and utilizes a glucose-6-phosphate isomerase in the last step. The pentose phosphate pathway (PPP) is a ubiquitous pathway in most bacteria and eukaryotes, as it plays a central role in cellular metabolism. It produces the important cellular intermediates ribose 5-phosphate and erythrose 4-phosphate, which are used for synthesis of nucleotides and aromatic amino acids, respectively (Soderberg 2005). In fact, erythrose 4-phosphate is one of the inputs to the Shikimate pathway, another IBD-enriched module discussed above. The PPP also produces NADPH, a reducing equivalent important for reductive reactions and prevention of oxidative stress (Kruger and von Schaewen 2003; Christodoulou et al. 2018). Beyond its link to other enriched amino acid biosynthesis pathways, it is unusual that such a central pathway would have an increased copy number in the IBD gut microbiome rather than being equally distributed across all samples. Some gut microbes are known to lack the transaldolase gene in this pathway and may instead encode an alternative pathway for pentose degradation called the sedoheptulose 1,7-bisphosphate pathway (SBPP) (Garschagen, Franke, and Deppenmeier 2021); it is therefore possible that the enrichment of the more common PPP in IBD is related to an increased ratio of microbial populations that use the PPP rather than the SBPP in the less-diverse microbiome of IBD patients, though this requires further investigation to verify.

The first carbon oxidation of the **citric acid cycle** (TCA cycle), which is a three-step conversion from oxaloacetate to 2-oxoglutarate (alpha-ketoglutarate), is enriched in the IBD samples. Similar to the PPP, the citric acid cycle is a central metabolic pathway, especially with regards to generation of energy and key metabolites for other pathways (Akram 2014). It is unclear why only this particular portion of the cycle would be enriched, though this could perhaps be attributed to the role of alpha-ketoglutarate in the production of glutamate, the precursor to proline, ornithine and arginine (three amino acids with enriched biosynthesis pathways in the IBD sample group, as discussed above). It has been said that 2-oxoglutarate is the most fundamental compound of this cycle, serving as the link between carbon and nitrogen metabolism and also as a critical element in the recovery of amine groups for amino acid and protein production (Pierzynowski and Pierzynowska 2022; Huergo Luciano F. and Dixon

Ray 2015). Thus, the enrichment of 2-oxoglutarate production capacity in the IBD gut environment could be related to the enrichment of amino acid biosynthesis pathways.

Two nucleotide sugar biosynthesis pathways are enriched in the IBD gut microbiome. One of these is **synthesis of UDP-glucose**, which is an important molecule implicated in a variety of key cellular metabolisms. It is an intermediate in polysaccharide biosynthesis and pyrimidine metabolism, a precursor of lipopolysaccharides in the outer cell membrane of Gram-negative bacteria, and an extracellular signaling molecule (Ralevic 2015)**.** Additionally, as an agonist for P2Y-14 receptors, it could play a role in modulating host gastrointestinal functions like muscular contraction (Bassil et al. 2009), and in modulating host inflammatory responses by activating this receptor specifically in T-lymphocytes (Scrivens and Dickenson 2005) and in immature monocyte-derived dendritic cells (MDDC) (Skelton et al. 2003). The other enriched nucleotide sugar pathway is **UDP-GlcNAc biosynthesis**. Flux through this pathway is linked to a multitude of other central metabolisms, including amino acid and fatty acid metabolism (Hardivillé and Hart 2014). Furthermore, UDP-GlcNAc is an important substrate in protein glycosylation pathways (Hardivillé and Hart 2014; Ryczko et al. 2016), and a precursor to critical cell wall components in bacteria (Yao Liu and Breukink 2016; Mikkola 2020; van Dam, Olrichs, and Breukink 2009). In the gut, this molecule has been implicated in regulation of nutrient uptake by the host (Ryczko et al. 2016).

**D-Glucuronate (glucuronic acid) degradation** into pyruvate and D-glyceraldehyde 3-phosphate is also enriched in the IBD gut microbiome. Some gut microbes are capable of growth on host-derived uronic acids (Lopez-Siles et al. 2012), so this pathway may serve as a source of energy to microbes living in the IBD gut environment. In mice, there is evidence that derivatives of glycosaminoglycan degradation such as D-glucuronate can worsen colitis (Lee et al. 2009).

Finally, the **phosphoribosyl diphosphate (PRPP) biosynthesis pathway** is important because PRPP is used in the formation of glycosidic bonds as well as in the biosynthesis of a number of cofactors, amino acids, and nucleotides (Hove-Jensen et al. 2017). It is discussed further below in the context of nucleotide metabolism.

#### Cofactor and vitamin pathways

Biosynthesis or salvage pathways for the following five cofactors and vitamins are enriched in IBD: **heme, siroheme, thiamine (vitamin B1), cobalamin (vitamin B12)**, and **coenzyme A (CoA)**. **Heme** is required for aerobic respiration (Gruss, Borezée-Durant, and Lechardeur 2012) and the increase in this pathway may be related to elevated oxygen levels in the gut as a result of inflammation, which promotes the growth of aerotolerant microbes (Shah 2016; Cevallos et al. 2019). Dietary heme has also been associated with gut dysbiosis, aggravated colitis, and increased cytotoxicity in the colon (Constante et al. 2017; Ijssennagger et al. 2015); and genes related to heme and siroheme biosynthesis have also been found with high abundance in infants with neonatal necrotizing enterocolitis (Claud et al. 2013).

Both **thiamine** and **cobalamin** are important cofactors that are commonly shared between gut microbes (Magnúsdóttir et al. 2015), suggesting that microbes incapable of synthesizing these cofactors are unlikely to thrive in the low-diversity microbial communities of the IBD gut environment. Neither of these vitamins is produced by host cells but they are typically acquired from dietary sources (cobalamin, in particular, is absorbed in the small intestine) (Seetharam and Alpers 1982; Degnan, Taga, and Goodman 2014; Hossain, Amarasena, and Mayengbam 2022), so the enrichment of these pathways is unlikely to have a large impact on host health.

**Coenzyme A** can be produced from pantothenate (vitamin B5) by most gut microbes (Magnúsdóttir et al. 2015) and its biosynthesis has been described as ‘essential’ considering that CoA is required for a large number of enzymatic reactions (Spry, Kirk, and Saliba 2008; Leonardi et al. 2005). It is therefore interesting that this pathway appears to be enriched in the IBD gut microbiome, which implies a relative deficiency of CoA biosynthesis in the healthy gut microbiome. It is possible that the module is spuriously enriched, despite its low p-value of 5.7e-21, given the short length of this pathway – it has 3 major steps when the KEGG module definition is interpreted in a 'stepwise' fashion by anvi-estimate-metabolism, though there are in fact 5 chemical conversions (Supplementary Table 2a). An alternative possibility is that the KEGG Ortholog hidden Markov models (HMMs) for the required enzymes do not sufficiently represent the diversity of these proteins across the gut microbiota, which could cause this pathway to be undercounted due to lack of proper annotations.

#### Nucleotide pathways

The IBD-enriched modules include pathways for synthesis of the first complete **purine**, **inosine monophosphate (IMP)** as well as a series of **pyrimidine biosynthesis pathways** encoding the conversion from uridine monophosphate (UMP) to ribonucleotides (UDP/UTP, CDP/CTP) and finally to the cytosine deoxyribonucleotide (dCTP). The **phosphoribosyl diphosphate (PRPP) biosynthesis pathway** is also included in this list; though it is classified as central carbohydrate metabolism in KEGG due to its role in glycosidic bond formation, this molecule is an important precursor for nucleotide biosynthesis (both purines and pyrimidines) and synthesis of the amino acids tryptophan and histidine (Hove-Jensen et al. 2017). Though many microbes are capable of producing their own nucleotides, some – especially lactic acid bacteria – are not and rely on uptake of exogenous nucleosides and bases, which are converted to nucleotides via salvage pathways (Nygaard 2014; Kilstrup et al. 2005). Notably, these salvage pathways are not enriched in the IBD gut microbiome, suggesting that self-sufficiency in nucleotide biosynthesis (especially in the early stages in this process) is selected for in these communities. This also implies the importance of pyrimidine and purine cross-feeding in the healthy gut environment, which is supported by evidence that some gut microbes (e.g. *Bacteroides vulgatus*) actively secrete nucleosides in the colon (Wong, Fong, and Yu 2023; Teng et al. 2023).

#### Lipid pathways

Two lipid biosynthesis pathways – **initiation and elongation of fatty acids** – are enriched in IBD. Fatty acids are essential components of cell membranes and also serve as signaling molecules (Brown, Clardy, and Xavier 2023); thus, the ability to synthesize them is an important fitness determinant. For example, gut *Bacteroides* species that are deficient in sphingolipid production capabilities are much less resilient to oxidative stress than wild-type species (An et al. 2011). Since oxidative stress is a hallmark of IBD, it is possible that this environment selects for microbes capable of fatty acid biosynthesis.

#### Energy pathways

The **Pta-Ack pathway** is important for microbial energy production and adaptation to different growth conditions via the 'acetate switch', which enables either production or consumption of acetate depending on available nutrients (Wolfe 2005). Short-chain fatty acids (SCFAs) such as acetate serve as important energy sources to intestinal epithelial cells. They also play a role in regulating gut barrier function and host immune responses (Martin-Gallausiaux et al. 2021; Zhang et al. 2022), and impaired absorption and oxidation of SCFAs can contribute to the development of IBD (Zhang et al. 2022). Acetate promotes host intestinal IgA production and thereby has a protective effect against gut inflammation (W. Wu et al. 2017), but acetate levels are reduced in children with IBD (Treem et al. 1994). Further study is required to determine the flux direction of the Pta-Ack pathway and whether it contributes to the reduction of acetate in the IBD gut environment.

**CAM metabolism** is categorized as a carbon fixation pathway in the KEGG MODULE database yet is a short (2-step) pathway utilizing enzymes required in other common metabolisms. Its first step is catalyzed by phosphoenolpyruvate carboxylase (PEPCK), an enzyme that is involved in gluconeogenesis, serine biosynthesis, and carbon skeleton conversions in the citric acid cycle (J. Yang, Kalhan, and Hanson 2009). Its second step is catalyzed by malate dehydrogenases, a ubiquitous class of enzymes that convert 2-hydroxy acids to 2-keto acids and are involved in gluconeogenesis, the TCA cycle, glyoxylate bypass, and amino acid synthesis (Minarik et al. 2002; Musrati et al. 1998). The increase in this pathway in IBD gut microbiomes could be attributed in part to the increase in aerobic respiration due to elevated oxygen levels (Shah 2016;

Cevallos et al. 2019) and in part to the increase in amino acid biosynthesis capacity as evidenced by the multiple amino acid pathways that are also enriched.

#### Drug resistance pathways

The use of antibiotics to treat IBD and its complications is known to increase antibiotic resistance in the gut microbiome (Nitzan et al. 2016; Ledder 2019) and several studies have noted that individuals exposed to antibiotics are more likely to develop IBD (Kronman et al. 2012; Ungaro et al. 2014; Ledder 2019; Shaw, Blanchard, and Bernstein 2011). This potentially explains the enrichment of two drug resistance pathways in the IBD microbiome: **efflux pump MepA** (conferring multidrug resistance) and the ***bla* system** (conferring beta-lactam resistance), as higher rates of antibiotic exposure in this sample group naturally leads to selection for resistance phenotypes (Levy 2000; Alekshun and Levy 2007). Beta-lactamases in particular have been found with higher frequency in people with IBD (Vich Vila et al. 2018; Leung et al. 2012; Vaisman et al. 2013). Increased microbial drug resistance can heighten the risk of a severe infection such as *Clostridium difficile* infection (CDI) (Llor and Bjerrum 2014). CDI already occurs with higher frequency in individuals with IBD (Jodorkovsky, Young, and Abreu 2010), though the higher incidence of CDI is not necessarily linked to chronic antibiotic use in these individuals (at least in one retrospective study of Crohn's disease) (Roy and Lichtiger 2016). Regardless, antibiotic resistance is a global health problem that affects everyone, not just those with IBD.

#### Most enriched pathways in HMI reference genomes

The three HMI-associated pathways with the largest difference in average completion (>40%) between HMI and non-HMI reference genomes were **siroheme biosynthesis**, **cobalamin biosynthesis**, and **tryptophan biosynthesis** (Supplementary Table 3g). Siroheme and cobalamin biosynthesis represent complex pathways that require 6-8 and 11-13 enzymatic steps, respectively, and both compounds belong to the tetrapyrroles that are involved in various essential biological functions (Bryant, Neil Hunter, and Warren 2020). Siroheme is a cofactor required for nitrite and sulfite reduction and its biosynthetic pathway provides the precursors required for cobalamin biosynthesis.

Genes belonging to biosynthetic pathways of siroheme and cobalamin had higher average relative abundance in infants diagnosed with neonatal necrotizing enterocolitis (Claud et al. 2013), an inflammatory bowel condition affecting premature newborns. The siroheme biosynthesis pathway is upregulated in some human pathogens in response to high nitric oxide (NO) levels likely in relation to the NO detoxification function of nitrite reductase (Porrini et al. 2021). Increased NO levels are commonly associated with active inflammation in IBD (Soufli et al. 2016).

While **siroheme** is central to sulfite and nitrite reduction in prokaryotes, **cobalamin** (vitamin B12) is essential not only for the majority of gut microbes (~80%) (Kelly et al. 2019; Hossain, Amarasena, and Mayengbam 2022; Degnan et al. 2014) but also for the human host, and functions as a coenzyme in key metabolic pathways in humans and bacteria. However, only relatively few gut microbes (~20-40%) encode the metabolic pathway for its synthesis (Degnan et al. 2014; Magnúsdóttir et al. 2015; Kelly et al. 2019) and humans largely rely on cobalamin supplied via their diet. B12 deficiency in humans leads to reduced villi length (Berg et al. 1972) and may affect intestinal barrier functioning (Bressenot et al. 2013). However, microbially-produced cobalamin alone is insufficient to sustain the host's requirements (Magnúsdóttir et al. 2015). The high average completion of this complex pathway in reference genomes classified as HMI (86%) in contrast to non-HMI reference genomes (40%) demonstrates the importance of metabolic independence for the survival of microorganisms in stressed gut environments, whereas in a healthy gut environment cross-feeding of B-vitamins likely supports those microbes that do not have metabolic means to synthesize them (Magnúsdóttir et al. 2015).

**Tryptophan** is an essential amino acid that serves as a precursor for a variety of microbial (Alkhalaf and Ryan 2015) and human metabolites that play a potential role in IBD pathogenesis (Agus, Planchais, and Sokol 2018). Tryptophan metabolites mediate a variety of host microbe interactions in the human gut (Agus, Planchais, and Sokol 2018), contribute to gut barrier integrity, and exert anti-inflammatory functions (Bansal et al. 2010; Roager and Licht 2018). While fecal tryptophan concentrations can be elevated in IBD patients (Jansson et al. 2009), tryptophan host metabolism via the

Kynurenine pathway also appears to be elevated in disease, resulting in decreased serum levels of the amino acid (Nikolaus et al. 2017). At the same time, a tryptophan-deficient diet in mice is linked to intestinal inflammation and alterations of the microbial community composition (Hashimoto et al. 2012; Yusufu et al. 2021).

Overall, our data contains no evidence or indication for any direct links between the increased representation of microbial metabolic modules in IBD and the role of the products these metabolic activities yield in human disease states.

#### Characterizing cohort-specific metabolic capacity across the gradient of health and disease

We sought to evaluate the cohort-specific trends in metabolic capacity by computing the median per-population copy number of the 33 IBD-enriched modules within each sample from each study. Considering the heterogeneity within each sample group, we ordered the studies from most healthy to least healthy, using the cohort description from each publication to approximate relative healthiness based on the number and types of exclusions listed for healthy or non-IBD controls, or on the diagnostic criteria for people with IBD (Supplementary Table 1a).

The amount of detail provided as well as the gastrointestinal conditions considered varied between studies. To overcome this challenge, we placed more emphasis on exclusionary conditions to sort samples based on host health status. For instance, (Le Chatelier et al. 2013) and (Raymond et al. 2016) used the most stringent criteria in the selection of healthy individuals. Both studies excluded patients with gastrointestinal-related conditions like disease, surgery, and medication; medications affecting the immune system; or antibiotics. Additionally, (Le Chatelier et al. 2013) also excluded individuals diagnosed with type-2 diabetes while (Raymond et al. 2016) did not, and we assigned a higher ‘health score’ to samples classified as healthy by (Le Chatelier et al. 2013). Similarly, (Schirmer et al. 2018) applied more stringent exclusion criteria for samples classified as ‘non-IBD’ controls than (Franzosa et al. 2019) and was therefore considered a healthier cohort within that group.

To arrange samples classified as ‘IBD’ along a gradient of host health status, we considered similar cohorts to be more unhealthy if their diagnosis was supported by several lines of evidence. For example, (Schirmer et al. 2018) diagnosed IBD based on a screening colonoscopy and included existing patients with consistent diagnosis over the past 5+ years, (Lloyd-Price et al. 2019) required a combination of endoscopic and histopathologic evidence for diagnosis, and (Franzosa et al. 2019) only considered patients diagnosed via endoscopic, histopathologic, and radiographic approaches. Regardless, these cohorts are likely extremely similar in healthiness. Patients with the lowest health score were described by (Vineis et al. 2016) – with a cohort composed of total proctocolectomy patients with ileal pouches, some of which developed pouchitis.

Ordering the per-sample median PPCN values along this gradient of cohort health indicates that the HMI metric for gut microbial metabolic capacity increases as host health decreases (Supplementary Figure 6a). Therefore, HMI adequately captures the variability in gut environment conditions that challenge microbial survival.

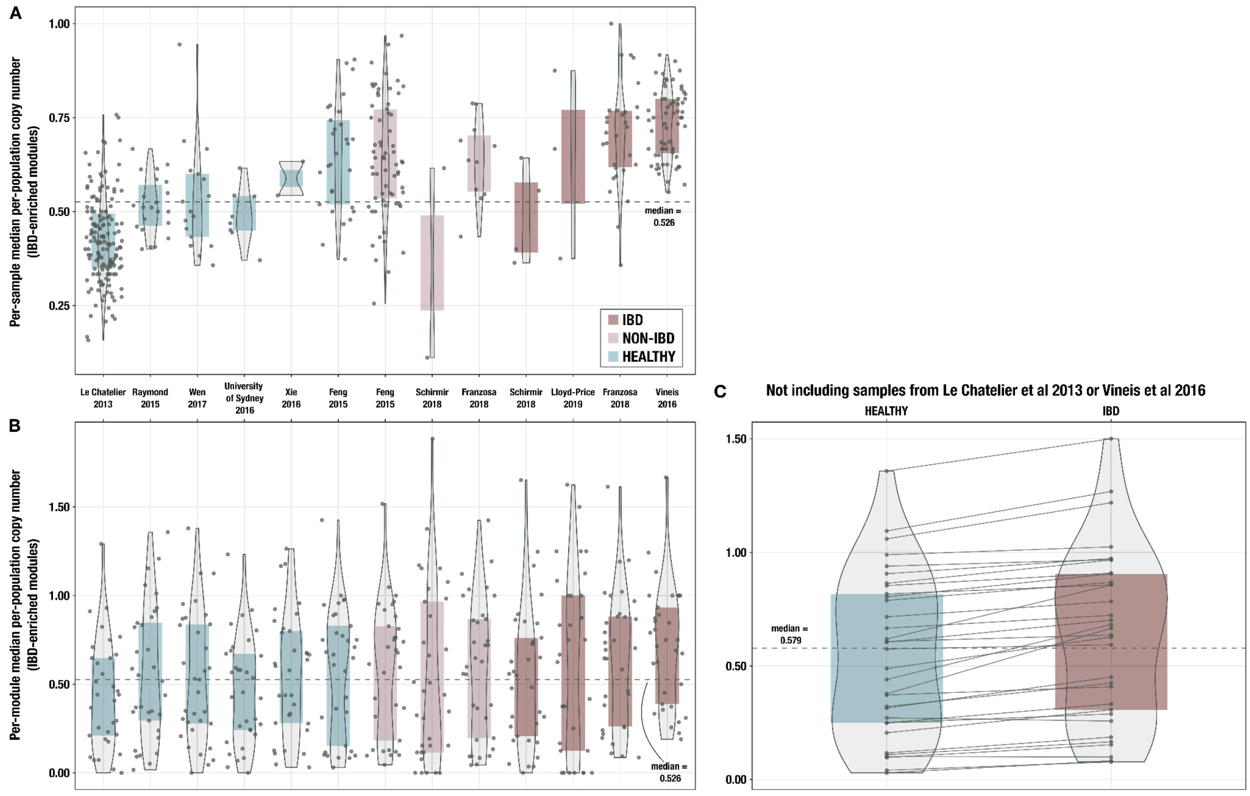


**Supplementary Figure 6. Boxplots of median per-population copy number of 33 IBD-enriched modules for samples from each individual cohort**, **A)** with medians computed within each sample (ie, one point per sample) and **B)** with medians computed for each IBD-enriched module (ie, one point per module). The x-axis indicates study of origin. **C)** Boxplots of median per-population copy number of 33 IBD-enriched modules for the 115 samples in the deeply-sequenced set that are not from (Le Chatelier et al. 2013) or (Vineis et al. 2016). The dashed line indicates the overall median for all 33 modules, and solid lines connect the points for the same module in each sample group.

### Considerations of batch effect

One concern in comparing samples from multiple studies is that differential sample processing strategies could contribute to the signal between groups; in other words, batch effects could partially explain the observed trends between different cohorts. However, by including samples from a variety of studies in each group (healthy, non-IBD and IBD) for our meta-analysis, we can mitigate the impact of batch effects on our observations. The similar distribution of the median normalized copy number for each of the 33 IBD-enriched metabolic modules (summarized across all samples within a given study), across all studies within a given sample group (Supplementary Figure 6b), confirms that the sample group explains more of the trend than the study of origin.

Two studies dominate our sample set: (Le Chatelier et al. 2013) contributes 151 (52.8%) of the healthy samples, and (Vineis et al. 2016) contributes 64 (63.4%) of the IBD samples (Supplementary Table 1b). To exclude that a cohort effect between these studies influences our observations, we repeated the IBD-enrichment analysis on (i) (Le Chatelier et al. 2013) and (Vineis et al. 2016) only; as well as on (ii) the remaining samples. While the results obtained from the two larger studies tend to have smaller p-values, the top IBD-enriched modules are broadly similar (Kendall correlation of Wilcoxon test p-values computed on two subsets: 0.59; see Supplementary Figure 7), demonstrating that we are capturing generic signals across studies in our sample set.

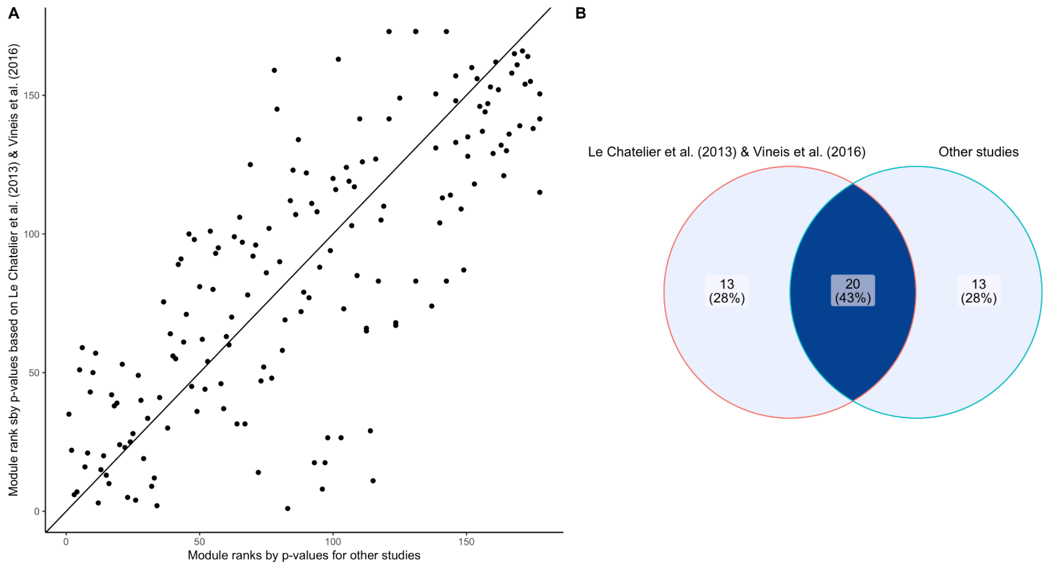


**Supplementary Figure 7. Assessing batch effect of the IBD-enrichment study. A)** Scatter plot comparing the module ranks of Wilcoxon-Mann-Whitney p-values comparing IBD and healthy subjects on (Le Chatelier et al. 2013) and (Vineis et al. 2016) (y axis) and the rest of our dataset (x axis). **B)** Venn diagram displaying the overlap of IBD-enriched modules identified by the 33 smallest p-values in (Le Chatelier et al. 2013) and (Vineis et al. 2016) and the rest of our dataset. There is good agreement (20 out of 33) between the two sets of modules, indicating generalizability of the signals across studies used in our sample set.

### Testing the generalizability of the metagenome classifier

To check whether performance of our logistic regression classifier was similar across the different studies in our sample set, we tested the model's performance using a leave-two-studies-out cross-validation strategy, whereby we trained the classifier on all samples except for those from one IBD study and one healthy study, and then tested it using samples from the two studies that were left out, for a total of 24 folds. Performance was quite variable across the different folds, as expected considering the large range of sample sizes from each study and the variability in health status of each cohort. The best overall performance occurred when testing on healthy samples from (Le Chatelier et al. 2013), with average accuracy of 89.9% across 3 folds. The worst performance occurred when testing on healthy samples from (Feng et al. 2015), with average accuracy of 43.1% across 4 folds. In the fold leaving out healthy samples from (Le Chatelier et al. 2013) and IBD samples from (Vineis et al. 2016), no IBD-enriched modules had p-values below our FDR-adjusted significance threshold of 2e-10 and therefore no classifier was trained. As these two studies contributed the largest number of samples to our deeply sequenced subset (Le Chatelier et al. 2013: n = 151 out of 330 or 45.8%, all of which were healthy samples. Vineis et al. 2016: n = 64 out of 330 or 19.4%, all of which were IBD samples), we considered that cohort-specific or study-specific effects could be driving the differential signal between healthy and IBD samples. To test this, we removed the samples from (Le Chatelier et al. 2013) and (Vineis et al. 2016) and ran 10-fold cross-validation using an 80-20 train-test split of the remaining 115 samples (37 IBD, 78 healthy), using the 33 IBD-enriched modules (computed from the full sample set) as features. We found that the model performed better than a naive classifier, with an average fold accuracy of 66.5%, average true Healthy rate of 69.4%, and an average true IBD rate of 61%. Therefore, while a portion of the signal in our initial analysis is indeed attributable to the differences between samples from (Le Chatelier et al. 2013) and (Vineis et al. 2016), the classifier still captures an IBD-specific signal across the other studies using this set of IBD-enriched pathways.

Furthermore, we note that the two dominating studies represent individuals at the extremes of the health gradient across our sample set, as described previously. The (Le Chatelier et al. 2013) cohort, with its numerous exclusionary conditions, contains the healthiest individuals, while the (Vineis et al. 2016) cohort of proctocolectomy and pouchitis patients contains the unhealthiest. It is therefore unsurprising that there is a large contrast in the metabolic potential of the gut microbiome in these individuals, considering the biological differences in their respective gut environments. This is also supported by the aforementioned ability of HMI to resolve the variability in host health, as demonstrated in Supplementary Figures 5b and 6a.

Constante, Marco, Gabriela Fragoso, Annie Calvé, Macha Samba-Mondonga, and Manuela M. Santos. 2017. “Dietary Heme Induces Gut Dysbiosis, Aggravates Colitis, and Potentiates the Development of Adenomas in Mice.” *Frontiers in Microbiology* 8 (September): 1809.

Dam, Vincent van, Nick Olrichs, and Eefjan Breukink. 2009. “Specific Labeling of Peptidoglycan Precursors as a Tool for Bacterial Cell Wall Studies.” *Chembiochem: A European Journal of Chemical Biology* 10 (4): 617–24.

Degnan, Patrick H., Natasha A. Barry, Kenny C. Mok, Michiko E. Taga, and Andrew L. Goodman. 2014. “Human Gut Microbes Use Multiple Transporters to Distinguish Vitamin $B_{12}$ Analogs and Compete in the Gut.” *Cell Host & Microbe* 15 (1): 47–57.

Degnan, Patrick H., Michiko E. Taga, and Andrew L. Goodman. 2014. “Vitamin B12 as a Modulator of Gut Microbial Ecology.” *Cell Metabolism* 20 (5): 769–78.

Dhakan, D. B., A. Maji, A. K. Sharma, R. Saxena, J. Pulikkan, T. Grace, A. Gomez, J. Scaria, K. R. Amato, and V. K. Sharma. 2019. “The Unique Composition of Indian Gut Microbiome, Gene Catalogue, and Associated Fecal Metabolome Deciphered Using Multi-Omics Approaches.” *GigaScience* 8 (3). https://doi.org/10.1093/gigascience/giz004.

Faure, Magali, Christine Mettraux, Denis Moennoz, Jean-Philippe Godin, Jacques Vuichoud, Florence Rochat, Denis Breuillé, Christiane Obled, and Irène Corthésy-Theulaz. 2006. “Specific Amino Acids Increase Mucin Synthesis and Microbiota in Dextran Sulfate Sodium-Treated Rats.” *The Journal of Nutrition* 136 (6): 1558–64.

Faure, Magali, Denis Moënnoz, Franck Montigon, Christine Mettraux, Denis Breuillé, and Olivier Ballèvre. 2005. “Dietary Threonine Restriction Specifically Reduces Intestinal Mucin Synthesis in Rats.” *The Journal of Nutrition* 135 (3): 486–91.

Feng, Qiang, Suisha Liang, Huijue Jia, Andreas Stadlmayr, Longqing Tang, Zhou Lan, Dongya Zhang, et al. 2015. “Gut Microbiome Development along the Colorectal Adenoma-Carcinoma Sequence.” *Nature Communications* 6 (March): 6528.

Franzosa, Eric A., Alexandra Sirota-Madi, Julian Avila-Pacheco, Nadine Fornelos, Henry J. Haiser, Stefan Reinker, Tommi Vatanen, et al. 2019. “Gut Microbiome Structure and Metabolic Activity in Inflammatory Bowel Disease.” *Nature Microbiology* 4 (2): 293–305.

Galperin, Michael Y., Kira S. Makarova, Yuri I. Wolf, and Eugene V. Koonin. 2015. “Expanded Microbial Genome Coverage and Improved Protein Family Annotation in the COG Database.” *Nucleic Acids Research* 43 (Database issue): D261–69.

Garschagen, Laura S., Thomas Franke, and Uwe Deppenmeier. 2021. “An Alternative Pentose Phosphate Pathway in Human Gut Bacteria for the Degradation of C5 Sugars in Dietary Fibers.” *The FEBS Journal* 288 (6): 1839–58.

Gobert, Alain P., Yulan Cheng, Mahmood Akhtar, Benjamin D. Mersey, Darren R. Blumberg, Raymond K. Cross, Rupesh Chaturvedi, et al. 2004. “Protective Role of Arginase in a Mouse Model of Colitis.” *Journal of Immunology* 173 (3): 2109–17.

Gojda, Jan, and Monika Cahova. 2021. “Gut Microbiota as the Link between Elevated BCAA Serum Levels and Insulin Resistance.” *Biomolecules* 11 (10). https://doi.org/10.3390/biom11101414.

Gruss, Alexandra, Elise Borezée-Durant, and Delphine Lechardeur. 2012. “Chapter Three - Environmental Heme Utilization by Heme-Auxotrophic Bacteria.” In *Advances in Microbial Physiology*, edited by Robert K. Poole, 61:69–124. Academic Press.

Hardivillé, Stéphan, and Gerald W. Hart. 2014. “Nutrient Regulation of Signaling, Transcription, and Cell Physiology by O-GlcNAcylation.” *Cell Metabolism* 20 (2): 208–13.

Hashimoto, Tatsuo, Thomas Perlot, Ateequr Rehman, Jean Trichereau, Hiroaki Ishiguro, Magdalena Paolino, Verena Sigl, et al. 2012. “ACE2 Links Amino Acid Malnutrition to Microbial Ecology and Intestinal Inflammation.” *Nature* 487 (7408): 477–81.

Hellmann, Jennifer, Allison Ta, Nicholas J. Ollberding, Ramona Bezold, Kathleen Lake, Kimberly Jackson, Kelsie Dirksing, Erin Bonkowski, David B. Haslam, and Lee A. Denson. 2023. “Patient-Reported Outcomes Correlate With Microbial Community Composition Independent of Mucosal Inflammation in Pediatric Inflammatory Bowel Disease.” *Inflammatory Bowel Diseases* 29 (2): 286–96.

Herrmann, Klaus M., and Lisa M. Weaver. 1999. “THE SHIKIMATE PATHWAY.” *Annual Review of Plant Physiology and Plant Molecular Biology* 50 (June): 473–503.

Hossain, Khandkar Shaharina, Sathya Amarasena, and Shyamchand Mayengbam. 2022. “B Vitamins and Their Roles in Gut Health.” *Microorganisms* 10 (6). https://doi.org/10.3390/microorganisms10061168.

Hove-Jensen, Bjarne, Kasper R. Andersen, Mogens Kilstrup, Jan Martinussen, Robert L. Switzer, and Martin Willemoës. 2017. “Phosphoribosyl Diphosphate (PRPP): Biosynthesis, Enzymology, Utilization, and Metabolic Significance.” *Microbiology and Molecular Biology Reviews: MMBR* 81 (1). https://doi.org/10.1128/MMBR.00040-16.

Huergo Luciano F., and Dixon Ray. 2015. “The Emergence of 2-Oxoglutarate as a Master Regulator Metabolite.” *Microbiology and Molecular Biology Reviews: MMBR* 79 (4): 419–35.

Ijssennagger, Noortje, Clara Belzer, Guido J. Hooiveld, Jan Dekker, Saskia W. C. van Mil, Michael Müller, Michiel Kleerebezem, and Roelof van der Meer. 2015. “Gut Microbiota Facilitates Dietary Heme-Induced Epithelial Hyperproliferation by Opening the Mucus Barrier in Colon.” *Proceedings of the National Academy of Sciences of the United States of America* 112 (32): 10038–43.

Jansson, Janet, Ben Willing, Marianna Lucio, Ages Fekete, Johan Dicksved, Jonas Halfvarson, Curt Tysk, and Philippe Schmitt-Kopplin. 2009. “Metabolomics Reveals Metabolic Biomarkers of Crohn’s Disease.” *PloS One* 4 (7): e6386.

Jodorkovsky, Daniela, Yuki Young, and Maria T. Abreu. 2010. “Clinical Outcomes of Patients with Ulcerative Colitis and Co-Existing Clostridium Difficile Infection.” *Digestive Diseases and Sciences* 55 (2): 415–20.

Johansson, Malin E. V., Jenny K. Gustafsson, Karolina E. Sjöberg, Joel Petersson, Lena Holm, Henrik Sjövall, and Gunnar C. Hansson. 2010. “Bacteria Penetrate the Inner Mucus Layer before Inflammation in the Dextran Sulfate Colitis Model.” *PloS One* 5 (8): e12238.

Johansson, Malin E. V., and Gunnar C. Hansson. 2016. “Immunological Aspects of Intestinal Mucus and Mucins.” *Nature Reviews. Immunology* 16 (10): 639–49.

Kananen, Kathryn, Iva A. Veseli, Christian J. Quiles Perez, Samuel E. Miller, A. Murat Eren, and Patrick H. Bradley. 2024. “Adaptive Adjustment of Significance Thresholds Produces Large Gains in Microbial Gene Annotations and Metabolic Insights.” *bioRxiv*. https://doi.org/10.1101/2024.07.03.601779.

Kanehisa, Minoru, Miho Furumichi, Yoko Sato, Masayuki Kawashima, and Mari Ishiguro-Watanabe. 2023. “KEGG for Taxonomy-Based Analysis of Pathways and Genomes.” *Nucleic Acids Research* 51 (D1): D587–92.

Katayama, Shigeru, and Yoshinori Mine. 2007. “Antioxidative Activity of Amino Acids on Tissue Oxidative Stress in Human Intestinal Epithelial Cell Model.” *Journal of Agricultural and Food Chemistry* 55 (21): 8458–64.

Kelly, Caleb J., Erica E. Alexeev, Linda Farb, Thad W. Vickery, Leon Zheng, Campbell Eric L,

David A. Kitzenberg, et al. 2019. “Oral Vitamin B12 Supplement Is Delivered to the Distal Gut, Altering the Corrinoid Profile and Selectively Depleting Bacteroides in C57BL/6 Mice.” *Gut Microbes* 10 (6): 654–62.

Kilstrup, Mogens, Karin Hammer, Peter Ruhdal Jensen, and Jan Martinussen. 2005. “Nucleotide Metabolism and Its Control in Lactic Acid Bacteria.” *FEMS Microbiology Reviews* 29 (3): 555–90.

Kim, Connie J., Jennifer A. Kovacs-Nolan, Chengbo Yang, Tania Archbold, Ming Z. Fan, and Yoshinori Mine. 2010. “L-Tryptophan Exhibits Therapeutic Function in a Porcine Model of Dextran Sodium Sulfate (DSS)-Induced Colitis.” *The Journal of Nutritional Biochemistry* 21 (6): 468–75.

Kolios, George, Vassilis Valatas, and Stephen G. Ward. 2004. “Nitric Oxide in Inflammatory Bowel Disease: A Universal Messenger in an Unsolved Puzzle.” *Immunology* 113 (4): 427–37.

Kronman, Matthew P., Theoklis E. Zaoutis, Kevin Haynes, Rui Feng, and Susan E. Coffin. 2012. “Antibiotic Exposure and IBD Development among Children: A Population-Based Cohort Study.” *Pediatrics* 130 (4): e794–803.

Kruger, Nicholas J., and Antje von Schaewen. 2003. “The Oxidative Pentose Phosphate Pathway: Structure and Organisation.” *Current Opinion in Plant Biology* 6 (3): 236–46.

Le Chatelier, Emmanuelle, Trine Nielsen, Junjie Qin, Edi Prifti, Falk Hildebrand, Gwen Falony, Mathieu Almeida, et al. 2013. “Richness of Human Gut Microbiome Correlates with Metabolic Markers.” *Nature* 500 (7464): 541–46.

Ledder, Oren. 2019. “Antibiotics in Inflammatory Bowel Diseases: Do We Know What We’re Doing?” *Translational Pediatrics* 8 (1): 42–55.

Lee, Hye-Sung, Song-Yi Han, Kwon-Yeul Ryu, and Dong-Hyun Kim. 2009. “The Degradation of Glycosaminoglycans by Intestinal Microflora Deteriorates Colitis in Mice.” *Inflammation* 32 (1): 27–36.

Leonardi, Roberta, Yong-Mei Zhang, Charles O. Rock, and Suzanne Jackowski. 2005. “Coenzyme A: Back in Action.” *Progress in Lipid Research* 44 (2-3): 125–53.

Leung, Wesley, Gurtej Malhi, Barbara M. Willey, Allison J. McGeer, Bjug Borgundvaag, Reka Thanabalan, Piraveina Gnanasuntharam, et al. 2012. “Prevalence and Predictors of MRSA, ESBL, and VRE Colonization in the Ambulatory IBD Population.” *Journal of Crohn’s & Colitis* 6 (7): 743–49.

Levy, S. B. 2000. “The Future of Antibiotics: Facing Antibiotic Resistance.” *Clinical Microbiology and Infection: The Official Publication of the European Society of Clinical Microbiology and Infectious Diseases* 6 Suppl 3: 101–6.

Lin, Rui, Wentian Liu, Meiyu Piao, and Hong Zhu. 2017. “A Review of the Relationship between the Gut Microbiota and Amino Acid Metabolism.” *Amino Acids* 49 (12): 2083–90.

Liu, Lan, Xin Guo, Jaladanki N. Rao, Tongtong Zou, Lan Xiao, Tingxi Yu, Jennifer A. Timmons, Douglas J. Turner, and Jian-Ying Wang. 2009. “Polyamines Regulate E-Cadherin Transcription through c-Myc Modulating Intestinal Epithelial Barrier Function.” *American Journal of Physiology. Cell Physiology* 296 (4): C801–10.

Liu, Yao, and Eefjan Breukink. 2016. “The Membrane Steps of Bacterial Cell Wall Synthesis as Antibiotic Targets.” *Antibiotics (Basel, Switzerland)* 5 (3). https://doi.org/10.3390/antibiotics5030028.

Liu, Yulan, Xiuying Wang, and Chien-An Andy Hu. 2017. “Therapeutic Potential of Amino Acids in Inflammatory Bowel Disease.” *Nutrients* 9 (9). https://doi.org/10.3390/nu9090920.

Llor, Carl, and Lars Bjerrum. 2014. “Antimicrobial Resistance: Risk Associated with Antibiotic Overuse and Initiatives to Reduce the Problem.” *Therapeutic Advances in Drug Safety* 5 (6): 229–41.

Lloyd-Price, Jason, Cesar Arze, Ashwin N. Ananthakrishnan, Melanie Schirmer, Julian Avila-Pacheco, Tiffany W. Poon, Elizabeth Andrews, et al. 2019. “Multi-Omics of the Gut

Microbial Ecosystem in Inflammatory Bowel Diseases." *Nature* 569 (7758): 655–62.
Lopez, Michael J., and Shamim S. Mohiuddin. 2023. *Biochemistry, Essential Amino Acids*. StatPearls Publishing.
Lopez-Siles, Mireia, Tanweer M. Khan, Sylvia H. Duncan, Hermie J. M. Harmsen, L. Jesús Garcia-Gil, and Harry J. Flint. 2012. "Cultured Representatives of Two Major Phylogroups of Human Colonic Faecalibacterium Prausnitzii Can Utilize Pectin, Uronic Acids, and Host-Derived Substrates for Growth." *Applied and Environmental Microbiology* 78 (2): 420–28.
Magnúsdóttir, Stefanía, Dmitry Ravcheev, Valérie de Crécy-Lagard, and Ines Thiele. 2015. "Systematic Genome Assessment of B-Vitamin Biosynthesis Suggests Co-Operation among Gut Microbes." *Frontiers in Genetics* 6 (April): 148.
Martin-Gallausiaux, Camille, Ludovica Marinelli, Hervé M. Blottière, Pierre Larraufie, and Nicolas Lapaque. 2021. "SCFA: Mechanisms and Functional Importance in the Gut." *The Proceedings of the Nutrition Society* 80 (1): 37–49.
McCormack, S. A., and L. R. Johnson. 1991. "Role of Polyamines in Gastrointestinal Mucosal Growth." *The American Journal of Physiology* 260 (6 Pt 1): G795–806.
Mesnage, Robin, and Michael N. Antoniou. 2020. "Computational Modelling Provides Insight into the Effects of Glyphosate on the Shikimate Pathway in the Human Gut Microbiome." *Current Research in Toxicology* 1 (June): 25–33.
Metges, C. C., A. E. El-Khoury, L. Henneman, K. J. Petzke, I. Grant, S. Bedri, P. P. Pereira, A. M. Ajami, M. F. Fuller, and V. R. Young. 1999. "Availability of Intestinal Microbial Lysine for Whole Body Lysine Homeostasis in Human Subjects." *The American Journal of Physiology* 277 (4): E597–607.
Mikkola, Satu. 2020. "Nucleotide Sugars in Chemistry and Biology." *Molecules* 25 (23). https://doi.org/10.3390/molecules25235755.
Minarik, P., N. Tomaskova, M. Kollarova, and M. Antalik. 2002. "Malate Dehydrogenases-Structure and Function." *General Physiology and Biophysics* 21 (3): 257–66.
Mistry, Jaina, Sara Chuguransky, Lowri Williams, Matloob Qureshi, Gustavo A. Salazar, Erik L. L. Sonnhammer, Silvio C. E. Tosatto, et al. 2021. "Pfam: The Protein Families Database in 2021." *Nucleic Acids Research* 49 (D1): D412–19.
Morgan, Xochitl C., Timothy L. Tickle, Harry Sokol, Dirk Gevers, Kathryn L. Devaney, Doyle V. Ward, Joshua A. Reyes, et al. 2012. "Dysfunction of the Intestinal Microbiome in Inflammatory Bowel Disease and Treatment." *Genome Biology* 13 (9): R79.
Musrati, R. A., M. Kollárová, N. Mernik, and D. Mikulásová. 1998. "Malate Dehydrogenase: Distribution, Function and Properties." *General Physiology and Biophysics* 17 (3): 193–210.
Nikolaus, Susanna, Berenice Schulte, Natalie Al-Massad, Florian Thieme, Dominik M. Schulte, Johannes Bethge, Ateequr Rehman, et al. 2017. "Increased Tryptophan Metabolism Is Associated With Activity of Inflammatory Bowel Diseases." *Gastroenterology* 153 (6): 1504–16.e2.
Nitzan, Orna, Mazen Elias, Avi Peretz, and Walid Saliba. 2016. "Role of Antibiotics for Treatment of Inflammatory Bowel Disease." *World Journal of Gastroenterology: WJG* 22 (3): 1078–87.
Nygaard, Per. 2014. "Purine and Pyrimidine Salvage Pathways." In *Bacillus Subtilis and Other Gram-Positive Bacteria*, 359–78. *Bacillus Subtilis*and Other Gram-Positive Bacteria. Washington, DC, USA: ASM Press.
Oz, Helieh S., Theresa S. Chen, Craig J. McClain, and Willem J. S. de Villiers. 2005. "Antioxidants as Novel Therapy in a Murine Model of Colitis." *The Journal of Nutritional Biochemistry* 16 (5): 297–304.
Pierzynowski, Stefan, and Kateryna Pierzynowska. 2022. "Alpha-Ketoglutarate, a Key Molecule Involved in Nitrogen Circulation in Both Animals and Plants, in the Context of Human Gut

Microbiota and Protein Metabolism.” *Advances in Medical Sciences* 67 (1): 142–47.
Porrini, Constance, Cyprien Guérin, Seav-Ly Tran, Rozenn Dervyn, Pierre Nicolas, and Nalini Ramarao. 2021. “Implication of a Key Region of Six Bacillus Cereus Genes Involved in Siroheme Synthesis, Nitrite Reductase Production and Iron Cluster Repair in the Bacterial Response to Nitric Oxide Stress.” *International Journal of Molecular Sciences* 22 (10). https://doi.org/10.3390/ijms22105079.
Ralevic, V. 2015. “UDP-Glucose☆.” In *Reference Module in Biomedical Sciences*. Elsevier.
Raymond, Frédéric, Amin A. Ouameur, Maxime Déraspe, Naeem Iqbal, Hélène Gingras, Bédis Dridi, Philippe Leprohon, et al. 2016. “The Initial State of the Human Gut Microbiome Determines Its Reshaping by Antibiotics.” *The ISME Journal* 10 (3): 707–20.
Rehman, Tahniat, Muhammad Asim Shabbir, Muhammad Inam-Ur-Raheem, Muhammad Faisal Manzoor, Nazir Ahmad, Zhi-Wei Liu, Muhammad Haseeb Ahmad, Azhari Siddeeg, Muhammad Abid, and Rana Muhammad Aadil. 2020. “Cysteine and Homocysteine as Biomarker of Various Diseases.” *Food Science & Nutrition* 8 (9): 4696–4707.
Rigottier-Gois, Lionel. 2013. “Dysbiosis in Inflammatory Bowel Diseases: The Oxygen Hypothesis.” *The ISME Journal* 7 (7): 1256–61.
Roager, Henrik M., and Tine R. Licht. 2018. “Microbial Tryptophan Catabolites in Health and Disease.” *Nature Communications* 9 (1): 3294.
Roy, Abhik, and Simon Lichtiger. 2016. “Clostridium Difficile Infection: A Rarity in Patients Receiving Chronic Antibiotic Treatment for Crohn’s Disease.” *Inflammatory Bowel Diseases* 22 (3): 648–53.
Ryczko, Michael C., Judy Pawling, Rui Chen, Anas M. Abdel Rahman, Kevin Yau, Julia K. Copeland, Cunjie Zhang, et al. 2016. “Metabolic Reprogramming by Hexosamine Biosynthetic and Golgi N-Glycan Branching Pathways.” *Scientific Reports* 6 (March): 23043.
Schirmer, Melanie, Eric A. Franzosa, Jason Lloyd-Price, Lauren J. McIver, Randall Schwager, Tiffany W. Poon, Ashwin N. Ananthakrishnan, et al. 2018. “Dynamics of Metatranscription in the Inflammatory Bowel Disease Gut Microbiome.” *Nature Microbiology* 3 (3): 337–46.
Scrivens, Michelle, and John M. Dickenson. 2005. “Functional Expression of the P2Y14 Receptor in Murine T-Lymphocytes.” *British Journal of Pharmacology* 146 (3): 435–44.
Seetharam, B., and D. H. Alpers. 1982. “Absorption and Transport of Cobalamin (vitamin B12).” *Annual Review of Nutrition* 2: 343–69.
Shah, Yatrik M. 2016. “The Role of Hypoxia in Intestinal Inflammation.” *Molecular and Cellular Pediatrics* 3 (1): 1.
Sharpton, Thomas, Svetlana Lyalina, Julie Luong, Joey Pham, Emily M. Deal, Courtney Armour, Christopher Gaulke, Shomyseh Sanjabi, and Katherine S. Pollard. 2017. “Development of Inflammatory Bowel Disease Is Linked to a Longitudinal Restructuring of the Gut Metagenome in Mice.” *mSystems* 2 (5). https://doi.org/10.1128/mSystems.00036-17.
Shaw, Souradet Y., James F. Blanchard, and Charles N. Bernstein. 2011. “Association between the Use of Antibiotics and New Diagnoses of Crohn’s Disease and Ulcerative Colitis.” *The American Journal of Gastroenterology* 106 (12): 2133–42.
Sherrill, C., and R. C. Fahey. 1998. “Import and Metabolism of Glutathione by Streptococcus Mutans.” *Journal of Bacteriology* 180 (6): 1454–59.
Singh, Kshipra, Alain P. Gobert, Lori A. Coburn, Daniel P. Barry, Margaret Allaman, Mohammad Asim, Paula B. Luis, et al. 2019. “Dietary Arginine Regulates Severity of Experimental Colitis and Affects the Colonic Microbiome.” *Frontiers in Cellular and Infection Microbiology* 9 (March): 66.
Skelton, Lisa, Mike Cooper, Marianne Murphy, and Adam Platt. 2003. “Human Immature Monocyte-Derived Dendritic Cells Express the G Protein-Coupled Receptor GPR105 (KIAA0001, P2Y14) and Increase Intracellular Calcium in Response to Its Agonist, Uridine Diphosphoglucose.” *Journal of Immunology* 171 (4): 1941–49.

Soderberg, Tim. 2005. “Biosynthesis of Ribose-5-Phosphate and Erythrose-4-Phosphate in Archaea: A Phylogenetic Analysis of Archaeal Genomes.” *Archaea* 1 (5): 347–52.

Sonnhammer, E. L., S. R. Eddy, and R. Durbin. 1997. “Pfam: A Comprehensive Database of Protein Domain Families Based on Seed Alignments.” *Proteins* 28 (3): 405–20.

Soufli, Imene, Ryma Toumi, Hayet Rafa, and Chafia Touil-Boukoffa. 2016. “Overview of Cytokines and Nitric Oxide Involvement in Immuno-Pathogenesis of Inflammatory Bowel Diseases.” *World Journal of Gastrointestinal Pharmacology and Therapeutics* 7 (3): 353–60.

Sprong, R. C., A. J. Schonewille, and R. van der Meer. 2010. “Dietary Cheese Whey Protein Protects Rats against Mild Dextran Sulfate Sodium-Induced Colitis: Role of Mucin and Microbiota.” *Journal of Dairy Science* 93 (4): 1364–71.

Spry, Christina, Kiaran Kirk, and Kevin J. Saliba. 2008. “Coenzyme A Biosynthesis: An Antimicrobial Drug Target.” *FEMS Microbiology Reviews* 32 (1): 56–106.

Teng, Huajing, Yan Wang, Xin Sui, Jiawen Fan, Shuai Li, Xiao Lei, Chen Shi, et al. 2023. “Gut Microbiota-Mediated Nucleotide Synthesis Attenuates the Response to Neoadjuvant Chemoradiotherapy in Rectal Cancer.” *Cancer Cell* 41 (1): 124–38.e6.

Tepe, Nalan, Leah A. Shimko, and Metin Duran. 2006. “Toxic Effects of Thiol-Reactive Compounds on Anaerobic Biomass.” *Bioresource Technology* 97 (4): 592–98.

Tong, Betty C., and Adrian Barbul. 2004. “Cellular and Physiological Effects of Arginine.” *Mini Reviews in Medicinal Chemistry* 4 (8): 823–32.

Treem, W. R., N. Ahsan, M. Shoup, and J. S. Hyams. 1994. “Fecal Short-Chain Fatty Acids in Children with Inflammatory Bowel Disease.” *Journal of Pediatric Gastroenterology and Nutrition* 18 (2): 159–64.

Ungaro, Ryan, Charles N. Bernstein, Richard Gearry, Anders Hviid, Kaija-Leena Kolho, Matthew P. Kronman, Souradet Shaw, Herbert Van Kruiningen, Jean-Frédéric Colombel, and Ashish Atreja. 2014. “Antibiotics Associated with Increased Risk of New-Onset Crohn’s Disease but Not Ulcerative Colitis: A Meta-Analysis.” *The American Journal of Gastroenterology* 109 (11): 1728–38.

Vaisman, Alon, Kevin Pivovarov, Allison McGeer, Barbara Willey, Bjug Borgundvaag, Vanessa Porter, Piraveina Gnanasuntharam, Yanliang Wei, and Geoffrey C. Nguyen. 2013. “Prevalence and Incidence of Antimicrobial-Resistant Organisms among Hospitalized Inflammatory Bowel Disease Patients.” *The Canadian Journal of Infectious Diseases & Medical Microbiology = Journal Canadien Des Maladies Infectieuses et de La Microbiologie Medicale / AMMI Canada* 24 (4): e117–21.

Vanni, Chiara, Matthew S. Schechter, Silvia G. Acinas, Albert Barberán, Pier Luigi Buttigieg, Emilio O. Casamayor, Tom O. Delmont, et al. 2022. “Unifying the Known and Unknown Microbial Coding Sequence Space.” *eLife* 11 (March). https://doi.org/10.7554/eLife.67667.

Vich Vila, Arnau, Floris Imhann, Valerie Collij, Soesma A. Jankipersadsing, Thomas Gurry, Zlatan Mujagic, Alexander Kurilshikov, et al. 2018. “Gut Microbiota Composition and Functional Changes in Inflammatory Bowel Disease and Irritable Bowel Syndrome.” *Science Translational Medicine* 10 (472). https://doi.org/10.1126/scitranslmed.aap8914.

Vineis, J. H., D. L. Ringus, H. G. Morrison, T. O. Delmont, S. Dalal, L. H. Raffals, D. A. Antonopoulos, et al. 2016. “Patient-Specific Bacteroides Genome Variants in Pouchitis. mBio 7.”

Walker, Miriam Y., Siddharth Pratap, Janet H. Southerland, Cherae M. Farmer-Dixon, Kesavalu Lakshmyya, and Pandu R. Gangula. 2018. “Role of Oral and Gut Microbiome in Nitric Oxide-Mediated Colon Motility.” *Nitric Oxide: Biology and Chemistry / Official Journal of the Nitric Oxide Society* 73 (February): 81–88.

Watson, Andrea R., Jessika Füssel, Iva Veseli, Johanna Zaal DeLongchamp, Marisela Silva, Florian Trigodet, Karen Lolans, et al. 2023. “Metabolic Independence Drives Gut Microbial Colonization and Resilience in Health and Disease.” *Genome Biology* 24 (1): 78.

Wolfe, Alan J. 2005. “The Acetate Switch.” *Microbiology and Molecular Biology Reviews: MMBR* 69 (1): 12–50.
Wong, Chi Chun, Winnie Fong, and Jun Yu. 2023. “Gut Microbes Promote Chemoradiotherapy Resistance via Metabolic Cross-Feeding.” *Cancer Cell* 41 (1): 12–14.
Wu, Liuting, Zhiru Tang, Huiyuan Chen, Zhongxiang Ren, Qi Ding, Kaiyang Liang, and Zhihong Sun. 2021. “Mutual Interaction between Gut Microbiota and Protein/amino Acid Metabolism for Host Mucosal Immunity and Health.” *Animal Nutrition (Zhongguo Xu Mu Shou Yi Xue Hui)* 7 (1): 11–16.
Wu, W., M. Sun, F. Chen, A. T. Cao, H. Liu, Y. Zhao, X. Huang, et al. 2017. “Microbiota Metabolite Short-Chain Fatty Acid Acetate Promotes Intestinal IgA Response to Microbiota Which Is Mediated by GPR43.” *Mucosal Immunology* 10 (4): 946–56.
Yang, Jianqi, Satish C. Kalhan, and Richard W. Hanson. 2009. “What Is the Metabolic Role of Phosphoenolpyruvate Carboxykinase?” *The Journal of Biological Chemistry* 284 (40): 27025–29.
Yang, Yuhui, Yuanhong Zhang, Yuncong Xu, Tingyu Luo, Yueting Ge, Yuge Jiang, Yonghui Shi, Jin Sun, and Guowei Le. 2019. “Dietary Methionine Restriction Improves the Gut Microbiota and Reduces Intestinal Permeability and Inflammation in High-Fat-Fed Mice.” *Food & Function* 10 (9): 5952–68.
Yusufu, Ibrahim, Kehong Ding, Kathryn Smith, Umesh D. Wankhade, Bikash Sahay, G. Taylor Patterson, Rafal Pacholczyk, et al. 2021. “A Tryptophan-Deficient Diet Induces Gut Microbiota Dysbiosis and Increases Systemic Inflammation in Aged Mice.” *International Journal of Molecular Sciences* 22 (9). https://doi.org/10.3390/ijms22095005.
Zhang, Zhilin, Huan Zhang, Tian Chen, Lin Shi, Daorong Wang, and Dong Tang. 2022. “Regulatory Role of Short-Chain Fatty Acids in Inflammatory Bowel Disease.” *Cell Communication and Signaling: CCS* 20 (1): 64.
Zhao, Hui, Juanjuan Chen, Xiaoping Li, Qiang Sun, Panpan Qin, and Qi Wang. 2019. “Compositional and Functional Features of the Female Premenopausal and Postmenopausal Gut Microbiota.” *FEBS Letters* 593 (18): 2655–64.
