## Supplementary File 2 for "Microbes with higher metabolic independence are enriched in human gut microbiomes under stress"

[1]Biophysical Sciences Program, The University of Chicago, Chicago, IL 60637, USA; [2]Department of Medicine, The University of Chicago, Chicago, IL 60637, USA; [3]Data Science Institute and Department of Biomedical Data Science, Stanford University, Stanford, CA, 94305, USA; [4]Committee on Microbiology, The University of Chicago, Chicago, IL 60637, USA; [5]MARUM Center for Marine Environmental Sciences, University of Bremen, Bremen, Germany; [6]Department of Biostatistics, University of Washington, Seattle, WA, 98195, USA; [7]Toyota Technological Institute at Chicago, Chicago, IL 60605, USA; [8]Lundbeck Foundation GeoGenetics Centre, GLOBE Institute, University of Copenhagen, Copenhagen, Denmark; [9]Institute for Chemistry and Biology of the Marine Environment, University of Oldenburg, Oldenburg, Germany; [10]Marine 'Omics Bridging Group, Max Planck Institute for Marine Microbiology, 28359 Bremen, Germany; [11]Alfred Wegener Institute for Polar and Marine Research, Bremerhaven, Germany; [12]Helmholtz Institute for Functional Marine Biodiversity, 26129, Oldenburg, Germany

### Comparison of anvi-estimate-metabolism to existing tools for metabolism reconstruction

There are two main strategies for estimation of metabolic potential from sequencing data. The first is metabolic modeling, in which genome-scale metabolic models (GSMMs) are built to computationally represent the network of available metabolic reactions for a particular organism (Fang, Lloyd, and Palsson 2020; Gu et al. 2019). This strategy enables mathematical modeling of metabolic fluxes, typically with the linear programming technique known as flux-balance analysis (FBA) (Orth, Thiele, and Palsson 2010), which contextualizes the metabolic network within a set of constraints and thereby enables simulation of particular physiological conditions (Sen and Orešič 2019). The second strategy is pathway prediction, which estimates the presence/absence and/or completeness of metabolic pathways to produce a summary of the metabolic capacity encoded in the input sequences. This technique has received less attention than metabolic modeling, but its results are more readily interpretable than models, and it is critical for understanding microbial functional roles without the need for a parameterized, *in silico* environment (Zhou et al. 2022). Both methods can be integrated with auxiliary information such as gene expression data or growth kinetics for validation of predicted metabolisms (Gu et al. 2019).

A variety of software tools exist for both types of metabolism reconstruction. Two early examples with basic approaches are the web-based server platforms KAAS (Moriya et al. 2007) and RAST (Aziz et al. 2008). KAAS simply highlights annotated enzymes within pathway maps from the KEGG database (Kanehisa et al. 2006), without producing any quantitative estimates. RAST similarly produces a limited summary of metabolism by categorizing enzymes into metabolic ‘subsystems’, but is also able to produce a metabolic model using the SEED infrastructure (DeJongh et al. 2007). There are a plethora of more contemporary modeling tools that generate GSMMs, including ModelSEED (Henry et al. 2010), RAVEN (Agren et al. 2013), merlin (Dias et al. 2015), CarveMe (Machado et al. 2018), and AuReMe (Aite et al. 2018). For comprehensive reviews about these tools, we refer the reader to several previous publications (Faria et al. 2018; Mendoza et al. 2019; Gu et al. 2019).

Software for pathway prediction include MinPath, DRAM, METABOLIC, and metaPathPredict. MinPath (Ye and Doak 2009) uses integer programming to determine the minimum set of pathways that explain an input set of annotations. DRAM (Shaffer et al. 2020) and METABOLIC (Zhou et al. 2022) both integrate annotation of genes from various enzyme databases with estimation of pathway completeness; DRAM is specialized for working with metagenome-assembled genomes (MAGs) while METABOLIC focuses on biogeochemical cycles. The goal of metaPathPredict (Geller-McGrath et al. 2023) is to produce better estimations for incomplete genomes (especially MAGs reconstructed from environmental samples) using machine learning models trained on reference databases.

Though most of these tools specialize in one method of metabolism reconstruction, some software – such as Pathway Tools, KBase, gapseq, and KEMET – have the capacity for both reconstruction strategies. Pathway Tools (Karp et al. 2015) is a primarily web-based platform for numerous functional analyses based upon a custom ‘omics data format called a Pathway/Genome Database (PGDB), which can be used for both FBA and querying available metabolic capacity. KBase (Arkin et al. 2018) is an online workspace for hosting scientific analyses on ‘omics datasets, and it contains apps for running existing metabolism software (such as DRAM, ModelSeed and Rast) on uploaded data. Both gapseq (Zimmermann, Kaleta, and Waschina 2021) and KEMET (Palù et al. 2022) were designed to produce more accurate metabolic models by incorporating a gap-filling process into their model generation workflows, and their pathway prediction capabilities are a side effect of this strategy. Gapseq achieves this via a novel linear programming algorithm and by utilizing a highly-curated reaction database, while KEMET uses pathway prediction results for updating the metabolic models that it creates by internally running CarveMe (Machado et al. 2018).

Within the landscape of these current tools, `anvi-estimate-metabolism` represents a software for pathway prediction, providing quantitative predictions of metabolic capacity by computing completeness scores for a predefined set of metabolic pathways. Anvi-estimate-metabolism distinguishes itself from the existing pathway prediction tools in several important ways. Firstly, it is the only current tool that calculates a pathway redundancy metric, to the best of our knowledge. It achieves this primarily by computing pathway copy numbers, which is an essential strategy for community-level analysis of metabolic capabilities. This program also generates an alternative metric for pathway redundancy by providing gene-level coverage values for enzymes within each pathway through its integration with the wider anvi'o codebase and data structures, assuming that read-recruitment results are available. Secondly, `anvi-estimate-metabolism` offers two distinct interpretation strategies for metabolic pathway definitions – a 'pathwise' strategy which considers all possible enzyme combinations ('paths') that would yield a complete pathway, and a 'stepwise' strategy that equally weighs alternative enzymes for the same reaction step. In other words, the specific enzymes used for a given metabolic conversion matter for the 'pathwise' metrics, but not for the 'stepwise' metrics. Each strategy is suitable for a different type of analysis – for instance, 'pathwise' metrics can be advantageous for studies of individual genomes while 'stepwise' metrics are appropriate for metagenomic analysis. Thus, the combinations of pathway interpretation strategies and metric type allow for the application of this tool to a variety of different research questions and input data types. In contrast, most of the other pathway prediction tools explicitly target genomic data.

Finally, `anvi-estimate-metabolism` is one of the only tools that supports user-defined metabolic pathways rather than exclusively relying on reference pathways (i.e., from KEGG). The program enables users to create their own pathway files, using enzyme annotations from any functional annotation source (including from various standard databases such as NCBI COGs, Pfam, and CAZymes as well as from custom annotations imported by the user into their anvi'o databases). The only other software with a similar feature is DRAM, which offers estimation from 'custom distillate' files.

However, DRAM's custom pathways entirely rely on enzymes from the KEGG Ortholog database.

### Validation of per-population copy number (PPCN) approach on simulated metagenomic data

Our novel approach for normalizing metabolic pathway copy numbers by the estimated number of populations within a community to get per-population copy numbers (PPCNs) required validation. We used simulated metagenomes to test the robustness of our approach to the following common parameters of microbial communities that could potentially influence our comparison between healthy and IBD gut metagenomes: genome size, community size (e.g., number of distinct microbial populations within a metagenome), and diversity level (e.g., number of distinct phyla). We generated these synthetic metagenomes by randomly combining bacterial and archaeal representative genomes from different species clusters in the Genome Taxonomy Database (GTDB) v95 (Parks et al. 2022) according to which parameter we wanted to test (see Supplementary Methods). These representative genomes included both isolate genomes and metagenome-assembled genomes that are not necessarily complete or well-studied, but represent a wide diversity of microbial taxa. Most of the samples we created were synthetic assemblies, generated by concatenating the contig sequences from each selected genome's FASTA into one file. To validate the full process starting from assembly, we also generated a test case starting from synthetic short reads (see Supplementary Methods).

We applied our PPCN approach to the synthetic metagenome assemblies to mimic our analysis of pathway completeness in gut metagenomes. This approach included gene annotation with KEGG KOfams and microbial single-copy core genes (SCGs), estimation of KEGG module copy numbers with `anvi-estimate-metabolism`, estimation of the number of populations in the community based on SCGs, and calculation of the normalized PPCN values from the resulting data. We then analyzed the accuracy of the PPCN values and their correlation with sample parameters.

#### Two metrics for PPCN accuracy relative to genomic values

Validating the accuracy of the PPCN calculation required comparison of computed PPCN values to the true per-population copy number within a given metagenome. Obtaining this ‘true’ value for each synthetic community is difficult without expert knowledge of each microbe’s metabolic capacities and extensive manual calculation. Therefore, we approximated the ‘true’ PPCN in a high-throughput manner by averaging the genomic pathway metrics within a given sample. We used `anvi-estimate-metabolism` to compute the stepwise completeness or copy number for each metabolic pathway within each genome in the synthetic community, then averaged these values. Though our ability to predict metabolic capacity from individual genomes is in itself limited by genome (in)completeness and missing annotations, these values can serve as a reference point for how well we summarize community-level metabolism given our current genome-level knowledge.

We used both average genomic completeness and average genomic copy number values because each metric has advantages and limitations when approximating the true PPCN value. Average genomic copy number could be considered the most direct analog to metagenomic PPCN, yet the copy number calculation in `anvi-estimate-metabolism` very conservatively does not take into account partial copies of a pathway. Even when a pathway is highly complete in a given genome, a copy is not counted unless 100% of the steps are present; hence, the average genomic copy number value can underestimate the true PPCN. Genomic completeness scores can capture these partial versions of a pathway, making them a better approximation when the synthetic community harbors multiple incomplete genomes. However, completeness scores cannot resolve multiple copies of the same pathway encoded in an individual genome and can underestimate the true PPCN value, especially for short or simple pathways. Given these limitations, we assessed the accuracy of our computed PPCN values using both metrics individually as reference points. We computed ‘PPCN error’ by subtracting either average genomic completeness or average genomic copy number from the metagenomic PPCN value.

#### The PPCN calculation is generally accurate relative to genomic values but can slightly overestimate community-level metabolic capacity

Across all our simulated test cases, we observed that metagenomic PPCN was typically very close to the average genomic metrics. Relative to average completeness scores, the distribution of PPCN error was centered close to 0 and somewhat right-skewed, with a mean error ranging from -0.13 to -0.10 and a standard deviation ranging from 0.18 to 0.22. Relative to average copy number, PPCN error was centered closer to 0 and left-skewed, with a mean error ranging from 0.04 to 0.06 and a standard deviation ranging from 0.10 to 0.11 (Supplementary Table 6a, Supplementary Figure 8). The error range was limited in both cases, but was much smaller for error computed relative to average genomic copy number. Thus, while overall quite accurate, metagenomic PPCN has a slight tendency to underestimate average genomic completeness and overestimate average genomic copy number. For the subset of IBD-enriched pathways, the PPCN error distributions showed similar trends but were centered at a slightly higher point, with a mean of -0.06 to -0.02 relative to average completeness and a mean of 0.14 to 0.17 relative to average copy number.

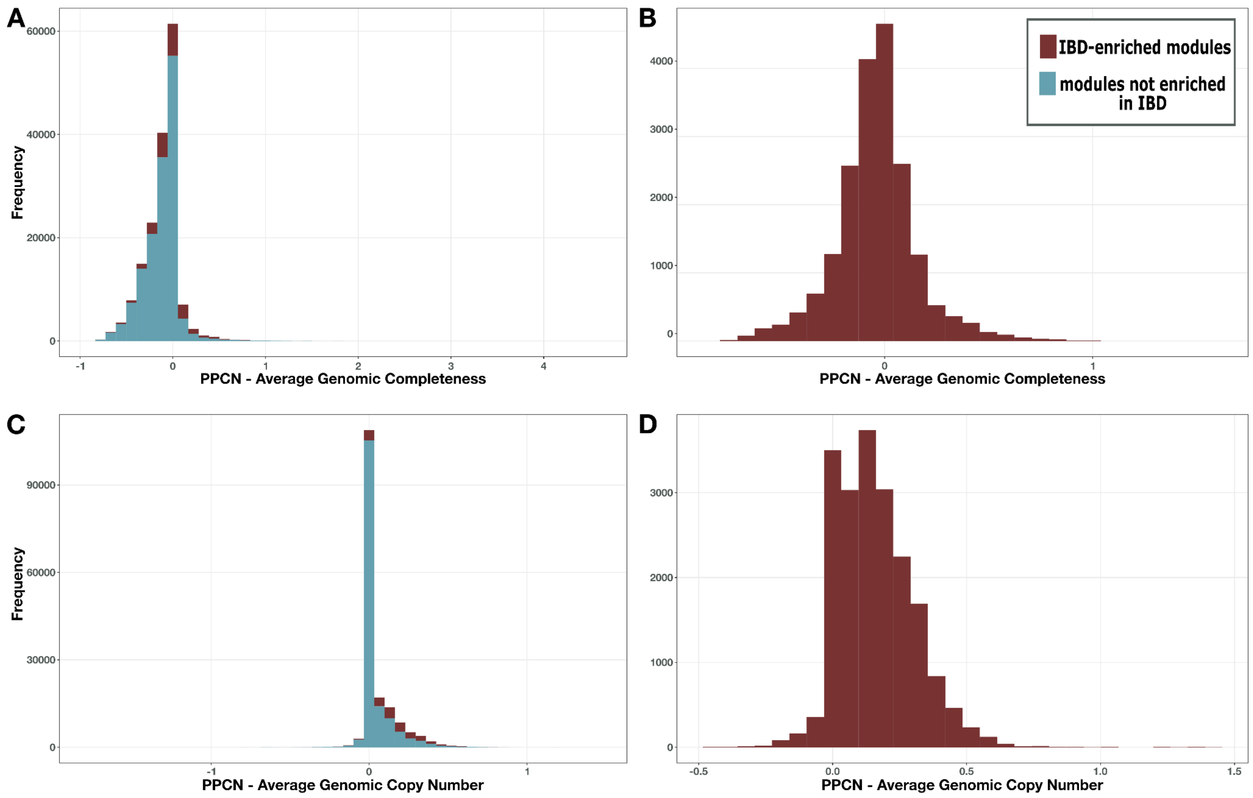


**Supplementary Figure 8.** Distribution of PPCN error relative to average genomic completeness (**A**, **B**) or average genomic copy number (**C**, **D**) for all modules (**A**, **C**) or just the IBD-enriched modules (**B**, **D**).

Examining the outliers in the underlying data revealed explanations for these trends. A primary reason for the overestimation of average genomic copy number was a phenomenon we term the ‘pathway complementarity effect’. When multiple members of a synthetic community contained partial yet complementary portions of a given metabolic pathway, these enzymes combined at the metagenomic level to produce additional ‘complete’ copies of the pathway. This effect was not observed when PPCN is compared to average genomic completeness scores because the latter metric takes partial copies of the pathway into account, thus reducing the observed error. Although cross-feeding is known to occur within microbial communities(Culp and Goodman 2023; Pacheco, Moel, and Segrè 2019) and in some cases pathway complementarity at the metagenome level could capture a legitimate biological signal, for the most part this is a technical artifact yielding a degree of error in the PPCN calculation, and is a natural outcome of the decision to consider the metagenome as one large pot of genes for the purposes of pathway prediction. That said, the magnitude of this error is typically small.

Metagenomic PPCN tends to slightly underestimate average genomic completeness due to the conservative nature of the copy number calculation. In cases when multiple members of a synthetic community have incomplete pathways (with non-zero completeness scores) and their respective portions of a given pathway are non-complementary, the average completeness scores will always be higher than the PPCN value, which does not count partial copies.

A few specific pathways had systematically overestimated PPCNs relative to one of the genomic metrics. For example, PPCN values for the beta-oxidation pathway (M00086) had the highest average error (0.946) relative to genomic completeness scores. This is an extremely short pathway, with only one reaction that can be catalyzed by one of two alternative enzymes. Thus, it represents an extreme case in which the copy number of the pathway is directly equivalent to the number of annotations for these two enzymes, which can lead to an extremely high copy number at the metagenome level. Within a given genome, the completeness of this pathway can never increase beyond 100% regardless of the number of annotations, and this limitation translates into a maximum average completeness score of 100% at the metagenome level. The systematic overestimation of PPCN for M00086 in this case is therefore due to the nature of the pathway itself and the limitation of average completeness score as a reference point. Relative to average genomic copy number, a module for glycolysis (M00001) had the highest average PPCN error (0.426) across all test cases. This is an extremely common metabolic pathway expected to occur in the majority of microbial genomes, which likely increases the pathway complementarity effect.

#### Accuracy of estimating the number of microbial populations within a metagenome

We also explored the accuracy of our method for estimating the number of populations from single-copy core genes. In general, this method has a slight tendency to underestimate the true number of populations, and errors are more likely when the actual community size is larger (Supplementary Figure 10). Regardless, the estimates were within 2 of the correct value over 90% of the time in all test cases for which we combined genomic contigs to create a synthetic metagenomic assembly. In the more realistic test case, when we generated synthetic short reads and assembled those reads *de novo*, the estimation accuracy dropped and was only within 2 of the correct value 67% of the time (Supplementary Table 6a). Accuracy increased with greater sequencing depth (Supplementary Figure 9), similar to what we observed in the gut metagenomes used in our main analysis (Supplementary Figure 1). This suggests that most errors in estimation were due to missing SCGs from incomplete genomes.

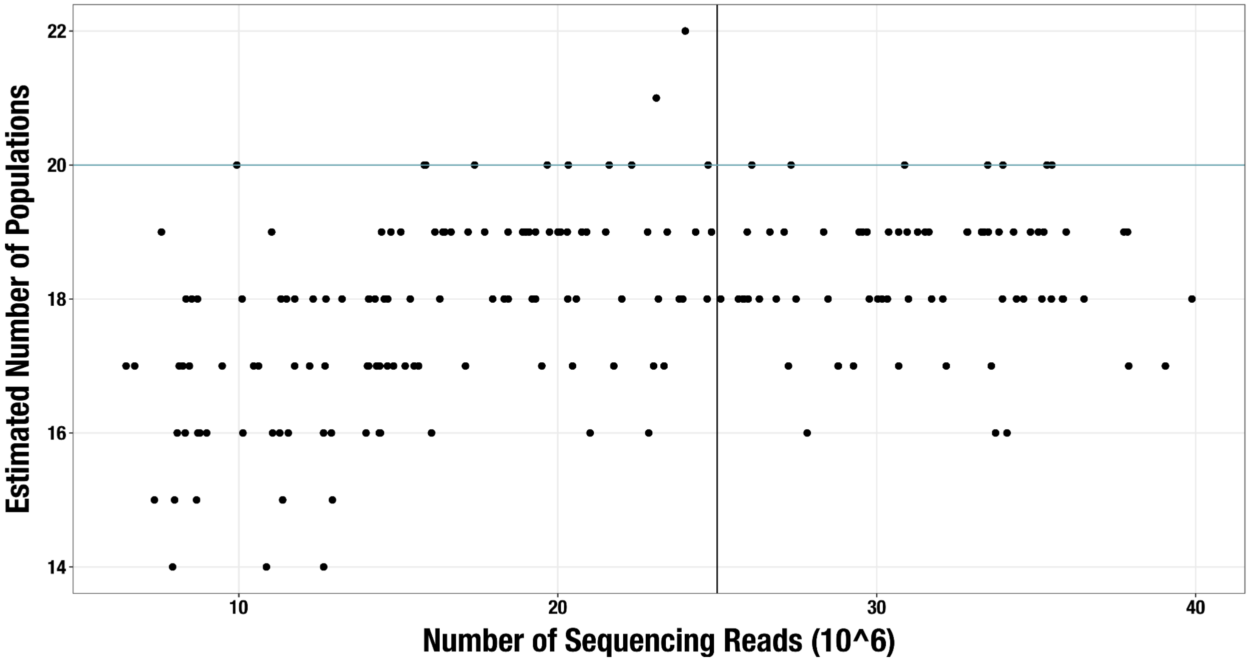


**Supplementary Figure 9. Scatterplot of sequencing depth vs estimated number of microbial populations in each of 189 'realistic' synthetic metagenome assemblies.** The blue line shows the actual number of genomes in each synthetic community (n = 20) and the black line shows the sequencing depth threshold used in our main analysis.

Since the estimated number of populations is the denominator in the PPCN calculation, underestimating these values can contribute to overestimation of the PPCN. This effect was not as strong as the pathway complementarity effect in the samples that we manually checked, which all belonged to the ideal test cases.

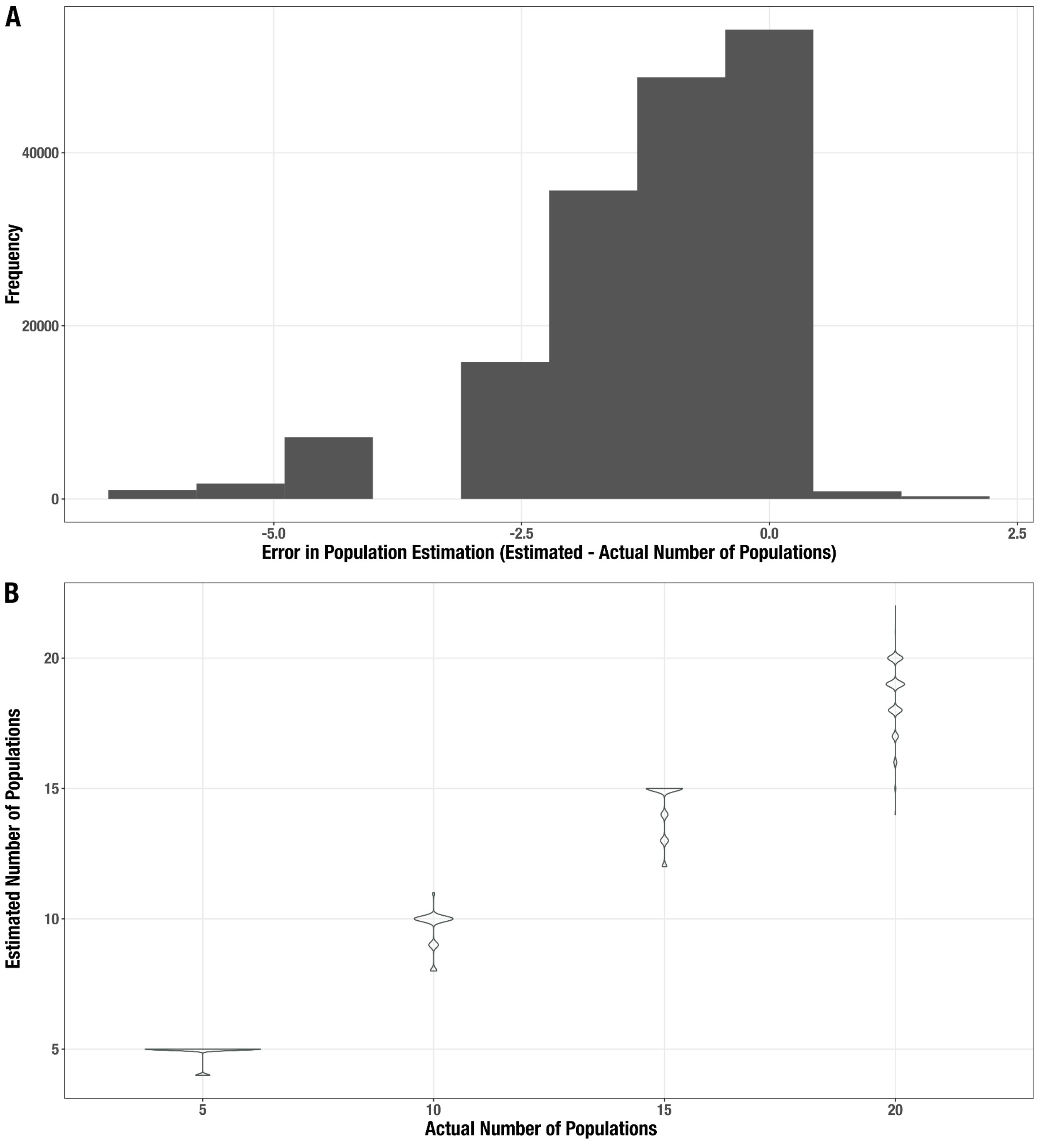


**Supplementary Figure 10. Distribution of error for the estimated number of populations in the synthetic metagenomes. A.** Histogram of the difference between estimated and actual community size. **B.** Distribution of estimates (y-axis) for each actual community size (x-axis).

#### The impact of genome size in an 'ideal' scenario

To explore how genome size influences the PPCN approach, we 'binned' genomes according to genome length to obtain the following size categories: small genomes (< 2 Mb), medium genomes (2 Mb up to 5 Mb), and large genomes (5 Mb up to 20 Mb). Genomes larger than 20 Mb in size were excluded. We then generated 189 random communities each containing 20 genomes from two size categories (S vs M, S vs L, and M vs L), such that each sample included a different proportion of genomes from each size category on a gradient from 0 to 1 (see Supplementary Methods). We independently analyzed each group of samples with the same size category pair with a Spearman's Correlation test to identify the relationship between genome size and PPCN, PPCN accuracy, and the estimated number of populations in the metagenome.

Across all pathways in the KEGG MODULE database, proportion of small genomes in a given sample had a weak negative correlation ($-0.09 < R < -0.02$) with PPCN values (Supplementary Figure 11, Supplementary Table 6b). The correlation became moderate ($-0.48 <= R <= -0.21$) for the subset of 33 IBD-enriched modules identified in our comparison of healthy and IBD gut metagenomes (Supplementary Figure 12, Supplementary Table 6b). All correlations were significant with a p-value threshold of $p < 0.05$, and the correlations were strongest for the S vs L group (Supplementary Table 6b). Thus, communities with more large genomes tend to have higher PPCN values, which makes sense considering that larger microbial genomes encode more genes and therefore more metabolic pathways. The stronger correlation for the subset of IBD-enriched modules mirrors our observation that IBD gut metagenomes harbor microbes with larger genomes with increased metabolic capacity (see 'Reference genomes with higher metabolic independence are over-represented in the gut metagenomes of individuals with IBD' in the main manuscript), and suggests that this subset of pathways is particularly likely to be found in large genomes regardless of environmental or taxonomic context.

The accuracy of the PPCN calculation had a weak correlation ($-0.02 < R < 0.07$) with proportion of small genomes, regardless of which genomic metric was used to approximate the error (Supplementary Figure 11, Supplementary Table 6b). The correlations became weakly negative for the subset of IBD-enriched pathways (-0.13 < R < 0.04), indicating that PPCN values are slightly more accurate for these pathways in communities of larger genomes, although several of the latter correlations were nonsignificant (Supplementary Figure 12, Supplementary Table 6b).

Proportion of small genomes had significant, moderate to strong negative correlations with the accuracy of community size estimates (-0.71 <= R <= -0.36; p < 1e-02), indicating that these estimates are more accurate when genomes in the community are larger (Supplementary Figure 13, Supplementary Table 6b). This might reflect a general tendency of larger genomes to contain more complete sets of SCGs.

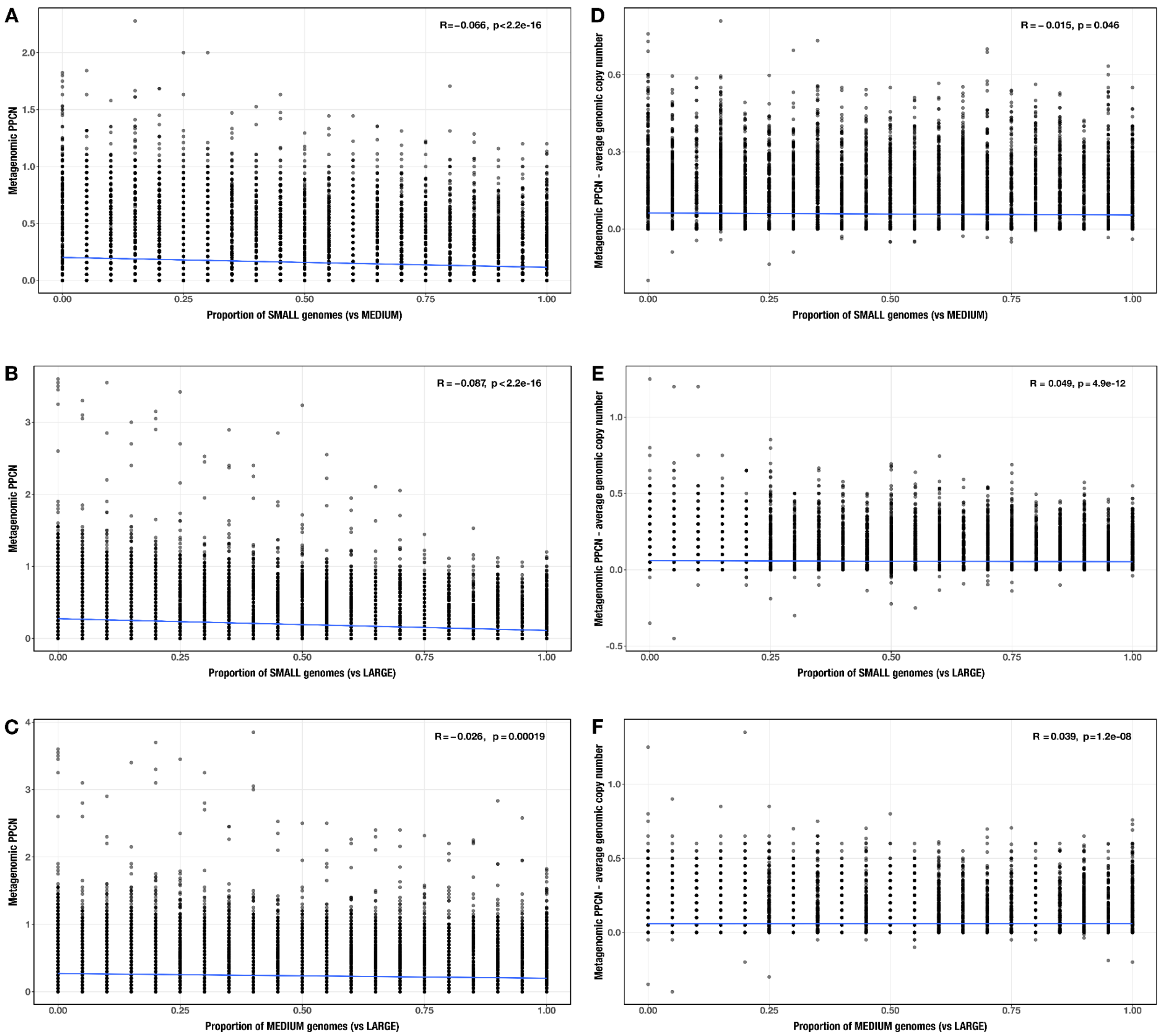


**Supplementary Figure 11.** Correlations between proportion of genomes in smaller size category and (**A – C**) PPCN or (**D – F**) PPCN error relative to average genomic copy number for each size category pair (**A**/**D**: small vs medium genomes; **B**/**E**: small vs large genomes; **C**/**F**: medium vs large genomes) across all modules. The Spearman's correlation coefficients and p-values are shown in the top-right corner of each plot, and regression lines are plotted in blue.

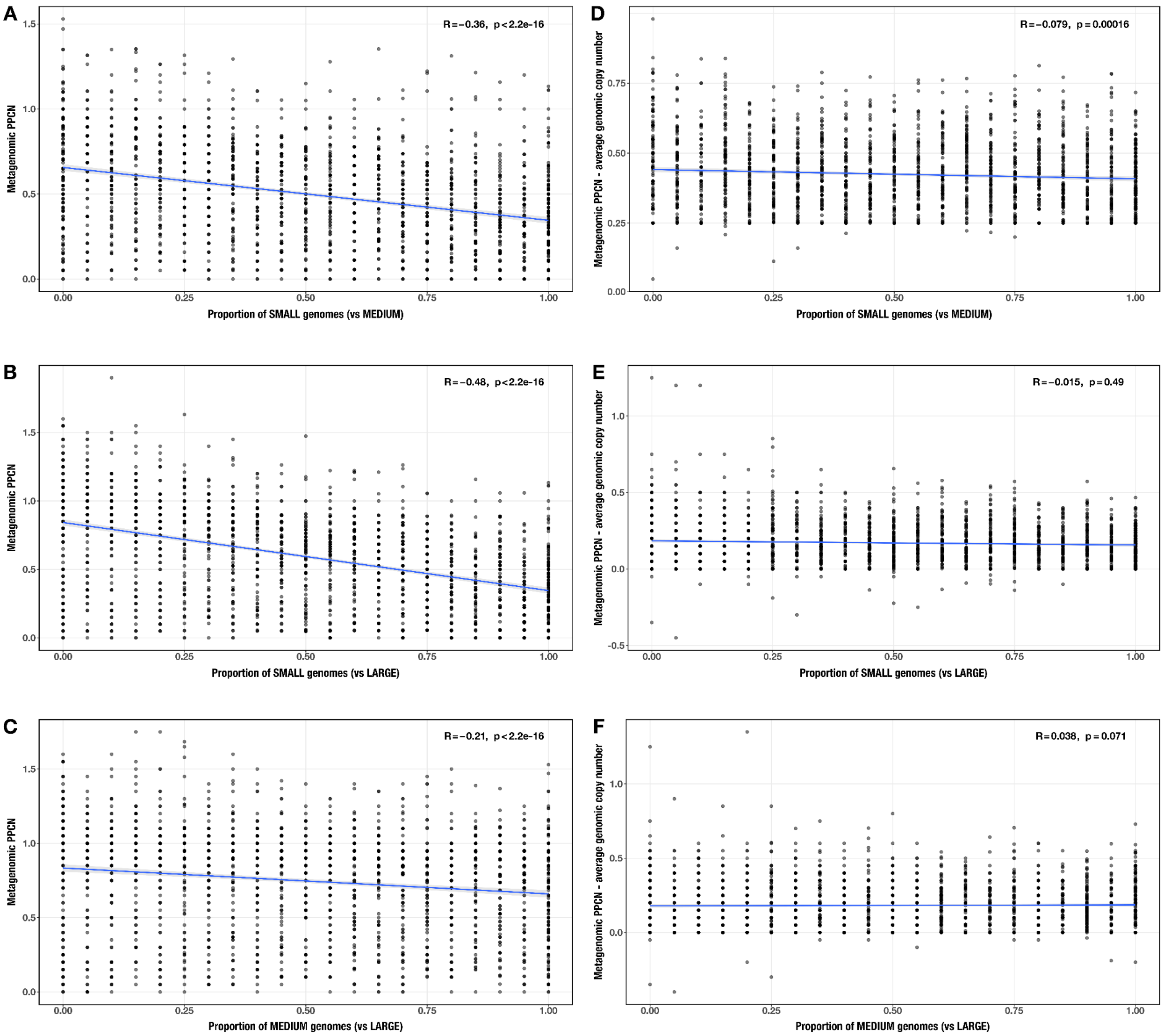


**Supplementary Figure 12.** Correlations between proportion of genomes in smaller size category and (**A – C**) PPCN or (**D – F**) PPCN error relative to average genomic copy number for each size category pair (**A**/**D**: small vs medium genomes; **B**/**E**: small vs large genomes; **C**/**F**: medium vs large genomes) across IBD-enriched modules (n=33). The Spearman's correlation coefficients and p-values are shown in the top-right corner of each plot, and regression lines are plotted in blue.

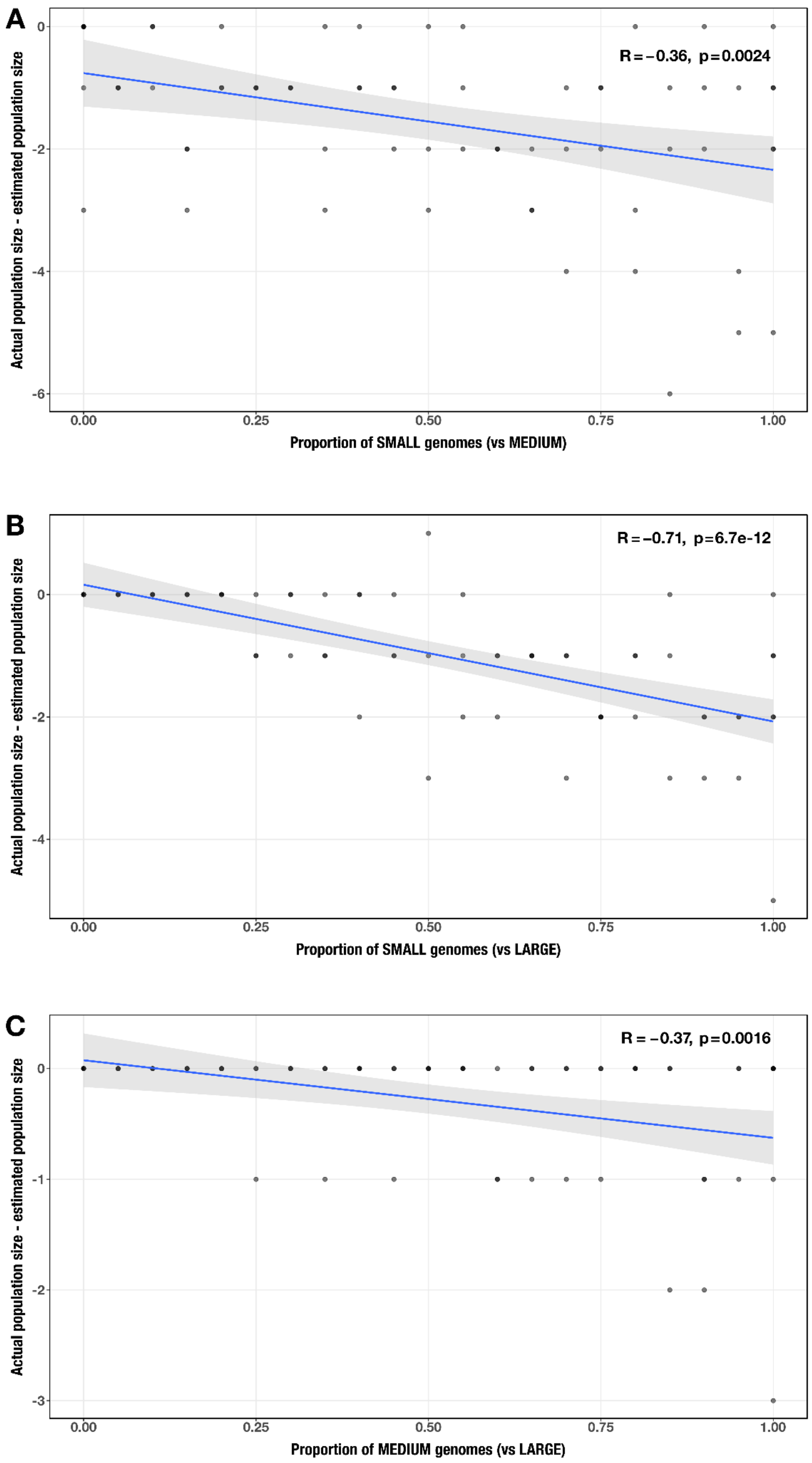


**Supplementary Figure 13.** Correlation between proportion of genomes in smaller size category and error in community size estimate (relative to actual community size) for each size category pair (**A**: small vs medium genomes; **B**: small vs large genomes; **C**: medium vs large genomes). The Spearman's correlation coefficients and p-values are shown in the top-right corner of each plot, and regression lines are plotted in blue.

#### The impact of genome size in a 'realistic' scenario

The prior test, which is based on the combination of pre-assembled genomic contigs into a synthetic metagenomic 'assembly', validates how our approach works for an 'ideal' scenario in which all community members can be assembled. However, a full metagenomic analysis workflow starts from highly fragmented sequencing reads, which must be assembled into contigs before gene annotation and other downstream analyses can be run. The assembly process can result in data loss if some microbial populations in the community cannot be fully assembled, which can impact downstream results. To simulate this entire process, we took the same 189 synthetic communities generated for the genome size test case and generated simulated short reads from each genome to create synthetic metagenome sequencing samples (see Supplementary Methods). We randomly assigned each genome in the community a relative abundance value from a normalized relative abundance curve of the top 20 most abundant populations in a healthy human gut metagenome (Supplementary Figure 14, Supplementary Table 6c, Supplementary Methods). We then converted the relative abundance values into coverage values ranging from 20x to 420x, and generated synthetic sequencing reads from each genome to produce its corresponding coverage value. Thus, each of the synthetic samples had the same coverage distribution across its 20 community members. After assembling the synthetic samples, we ran the PPCN workflow and performed the same validation described above.

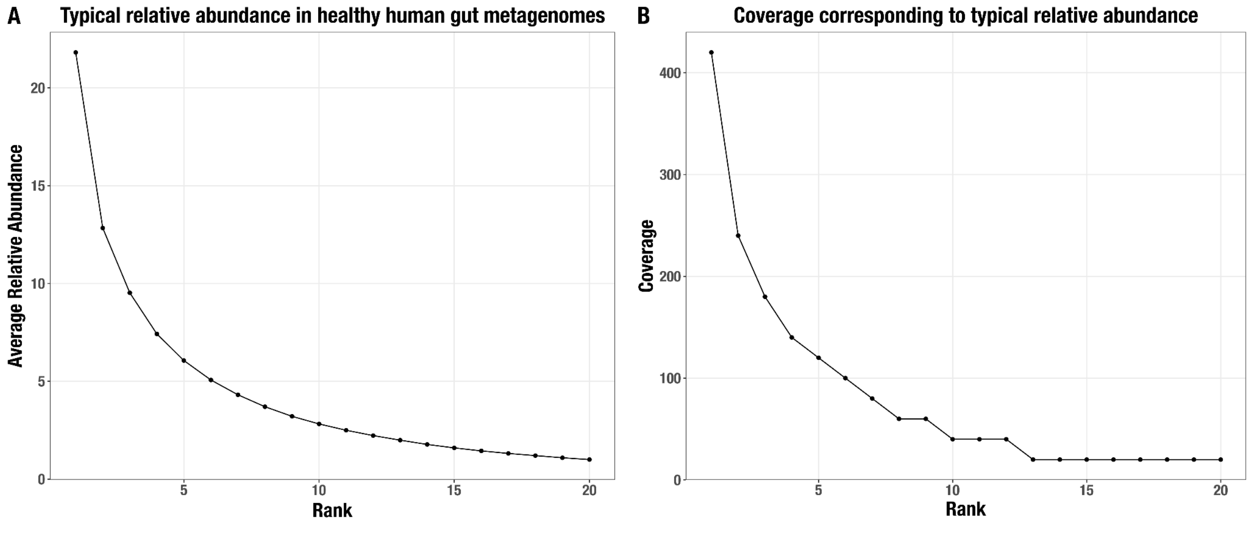

**Supplementary Figure 14.** Normalized average relative abundance curve **(A)** for the top 20 most abundant populations in a typical healthy human gut metagenome and **(B)** their corresponding coverage values in our synthetic metagenomes.

The validation results for this more realistic scenario showed largely the same trends as the ideal genome size case (Supplementary Figure 15, Supplementary Table 6ab) with two exceptions: (1) the accuracy of the population size estimates was lower, as mentioned previously. (2) The correlation between genome size proportion and population size estimation accuracy was weakly positive and nonsignificant for the M vs L group of genomes (Supplementary Figure 16, Supplementary Table 6ab). Given the similarity between the results for the ‘ideal’ case and the ‘realistic’ case, we decided to only test the less computationally-intensive ‘ideal’ case for the remaining parameters.

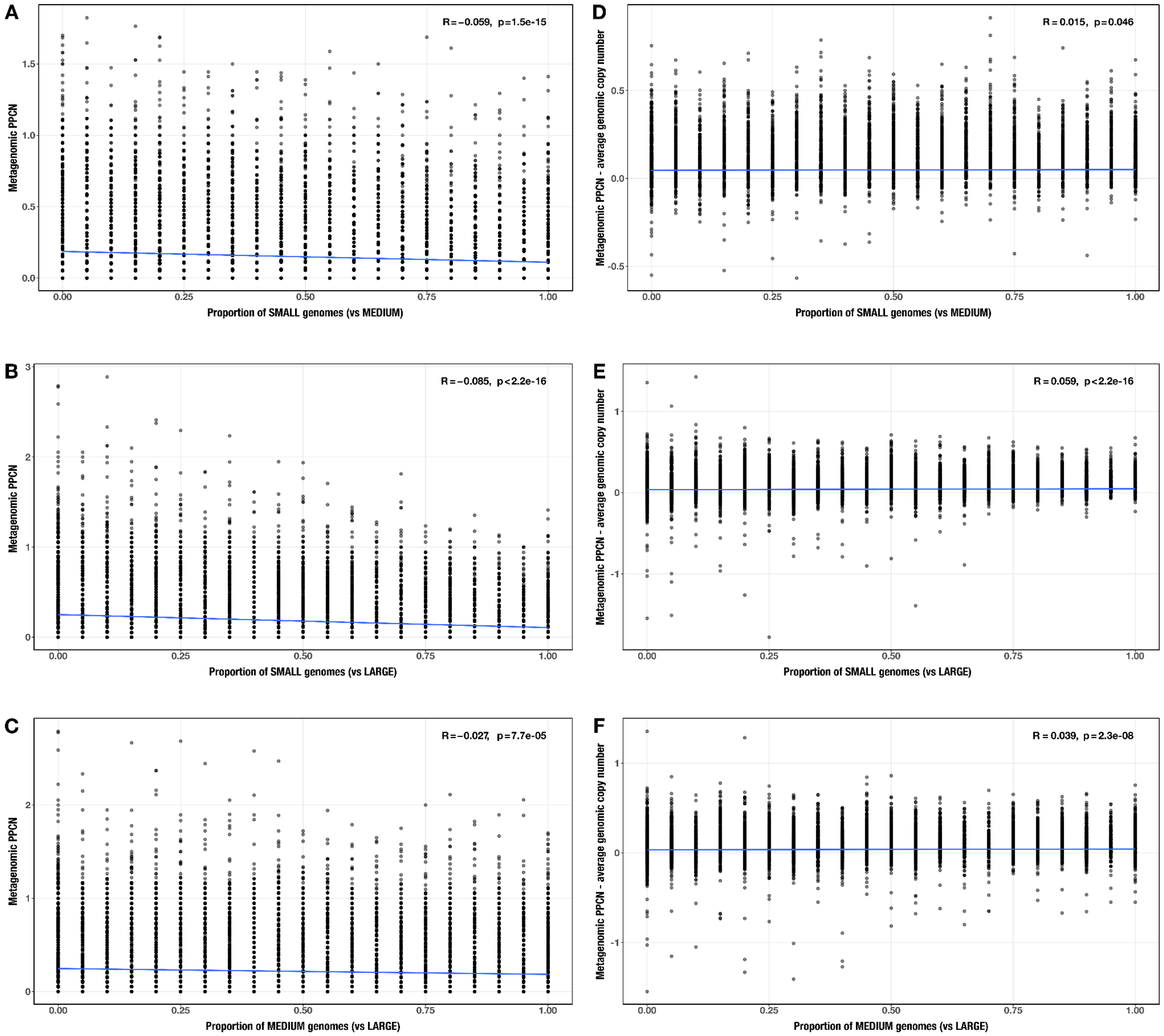

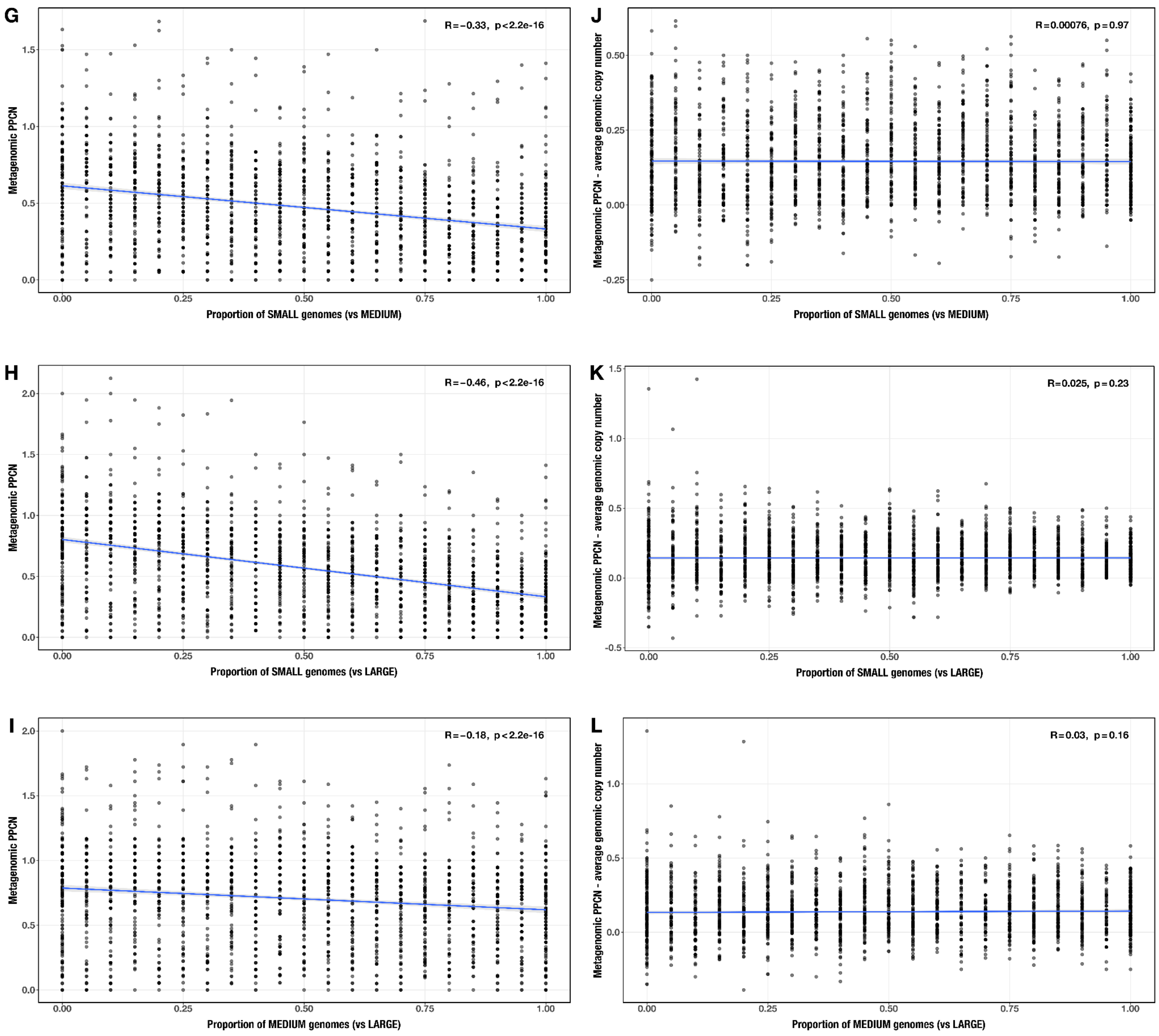


**Supplementary Figure 15.** Correlations between proportion of genomes in smaller size category and (**A** – **C**, **G** – **I**) PPCN or (**D** – **F**, **J** – **L**) PPCN error relative to average genomic copy number for each size category pair across all modules (**A** – **F**) or the subset of IBD-enriched modules (**G** – **L**) in the realistic genome size test case. The Spearman's correlation coefficients and p-values are shown in the top-right corner of each plot, and regression lines are plotted in blue.

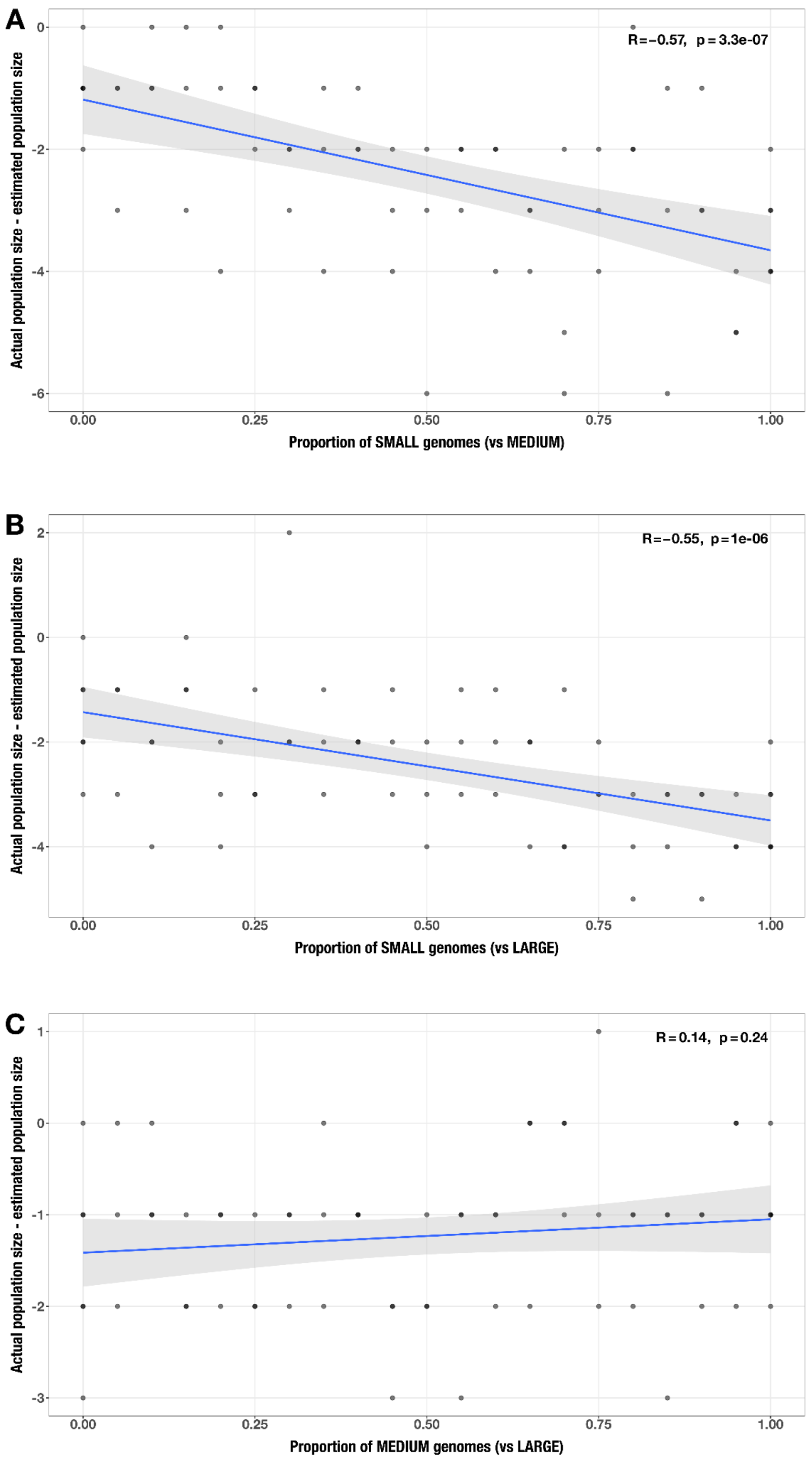


**Supplementary Figure 16.** Correlations between proportion of genomes in smaller size category and error in community size estimate (relative to actual community size) for each size category pair (**A**: small vs medium genomes; **B**: small vs large genomes; **C**: medium vs large genomes) in the realistic genome size test case. The Spearman's correlation coefficients and p-values are shown in the top-right corner of each plot, and regression lines are plotted in blue.

#### The impact of community size

To analyze the impact of community size on our approach, we generated 120 synthetic metagenome assemblies each containing 5, 10, 15, or 20 randomly-selected genomes from the same size category (S, M, or L; see Supplementary Methods). We independently analyzed each size category as described above.

As expected, we observed a significant positive correlation between metagenomic copy number (the numerator of PPCN) and community size in each group, likely driven by the increase in the copy number of core metabolic pathways in larger communities (Supplementary Figure 17). Interestingly, this correlation was much stronger for the subset of IBD-enriched pathways (0.49 <= R <= 0.67) than for all modules (0.12 <= R <= 0.13).

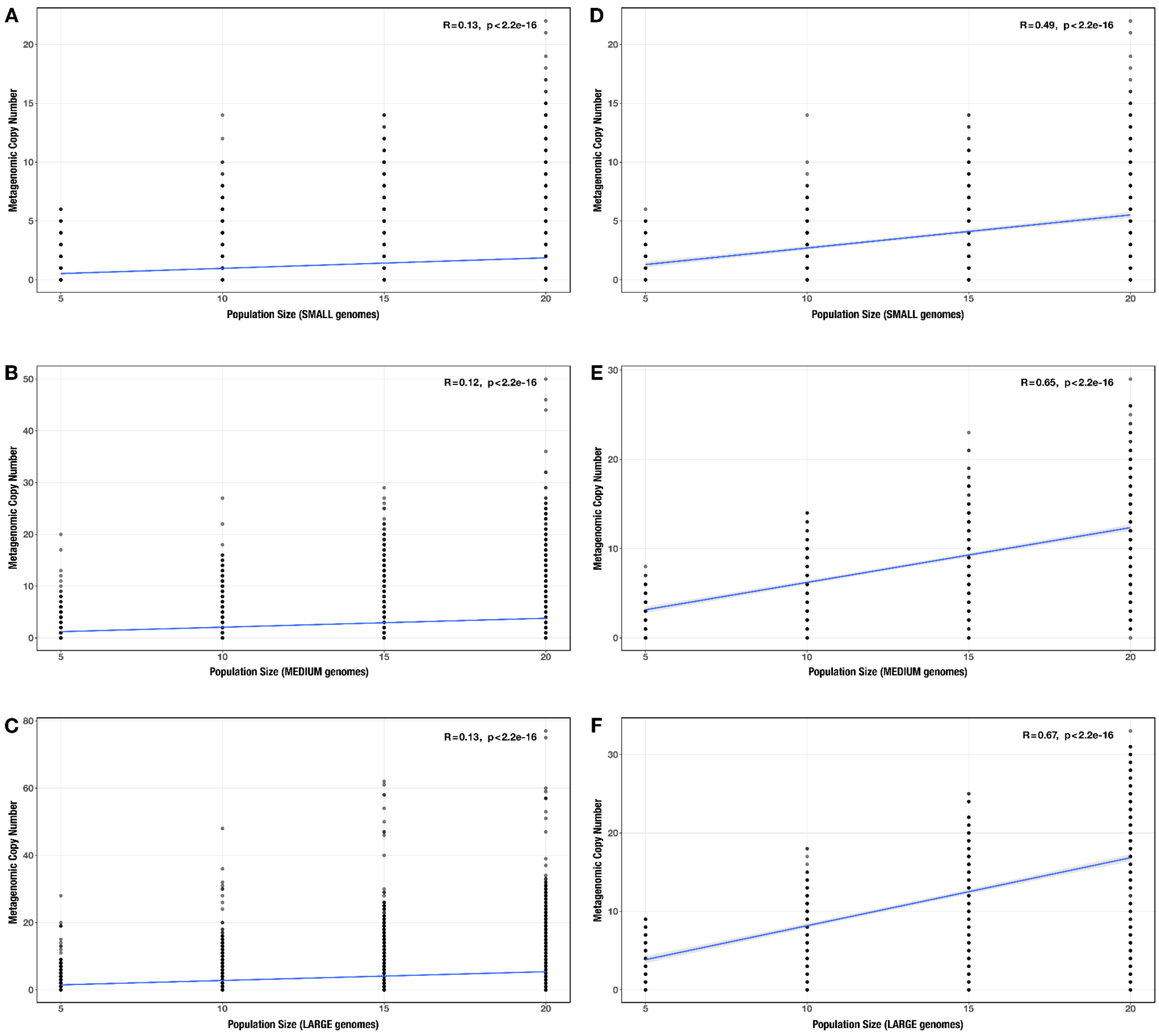


**Supplementary Figure 17.** Correlation between community size and metagenomic copy number across all modules (**A** – **C**) and across the subset of enriched modules (**D** – **F**) for each genome size category (**A**/**D**: small genomes; **B**/**E**: medium genomes; **C**/**F**: large genomes) in the community size test case. The Spearman's correlation coefficients and p-values are shown in the top-right corner of each plot, and regression lines are plotted in blue.

However, the correlation was much weaker and often nonsignificant for the normalized PPCN data in both groups of modules (all modules: $0.01 < R < 0.04$, enriched modules: $0.04 < R < 0.09$, Supplementary Table 6b, Supplementary Figure 18), which demonstrates the suitability of our normalization method to remove the effect of community size in comparisons of metagenome-level metabolic capacity.

The correlations between community size and PPCN accuracy were weakly yet significantly positive (0.07 < R <= 0.16), were slightly stronger for the subset of enriched modules (0.11 <= R <= 0.25), and indicate that PPCN values are slightly more accurate for smaller communities (Supplementary Table 6b, Supplementary Figure 18).

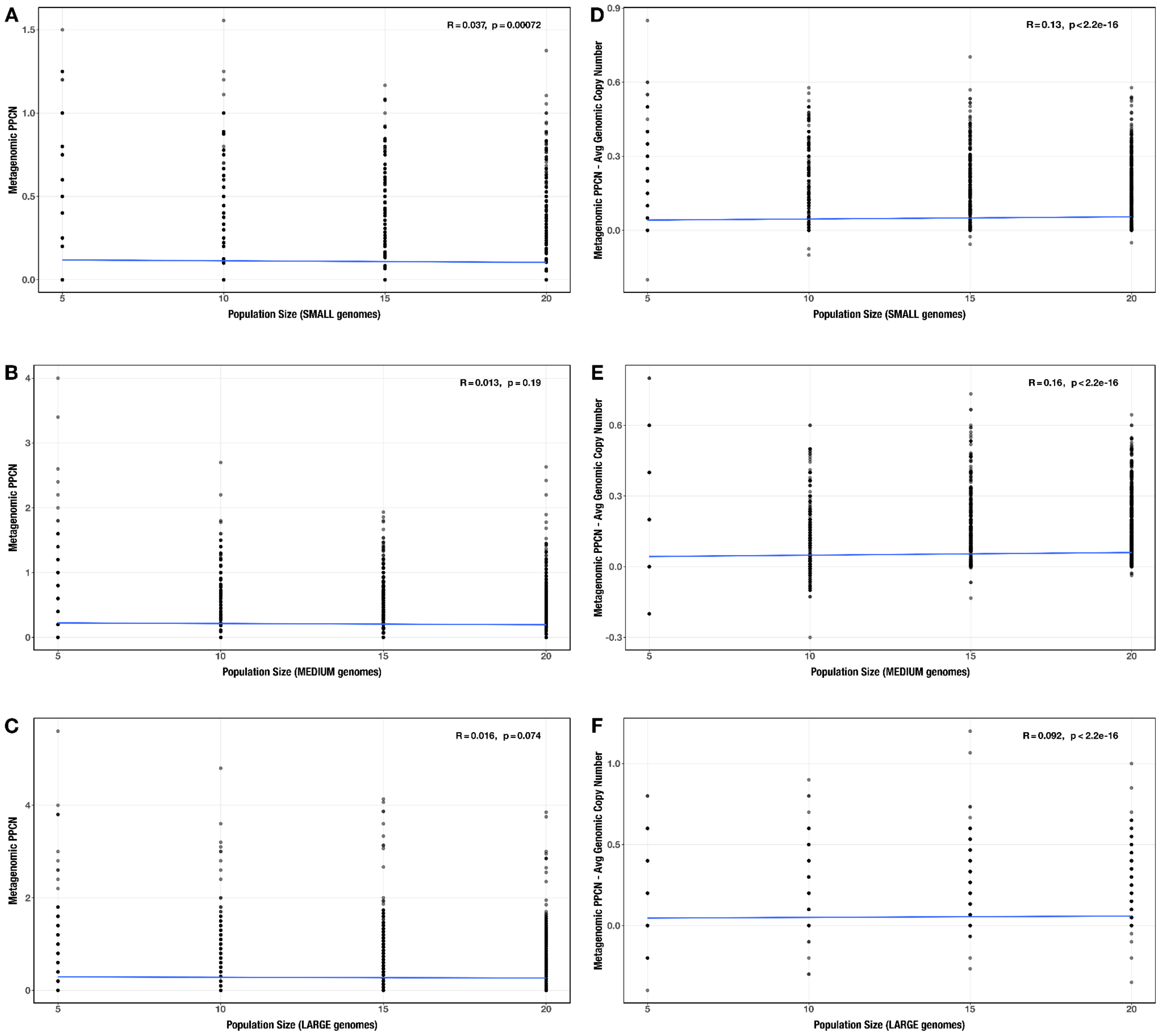

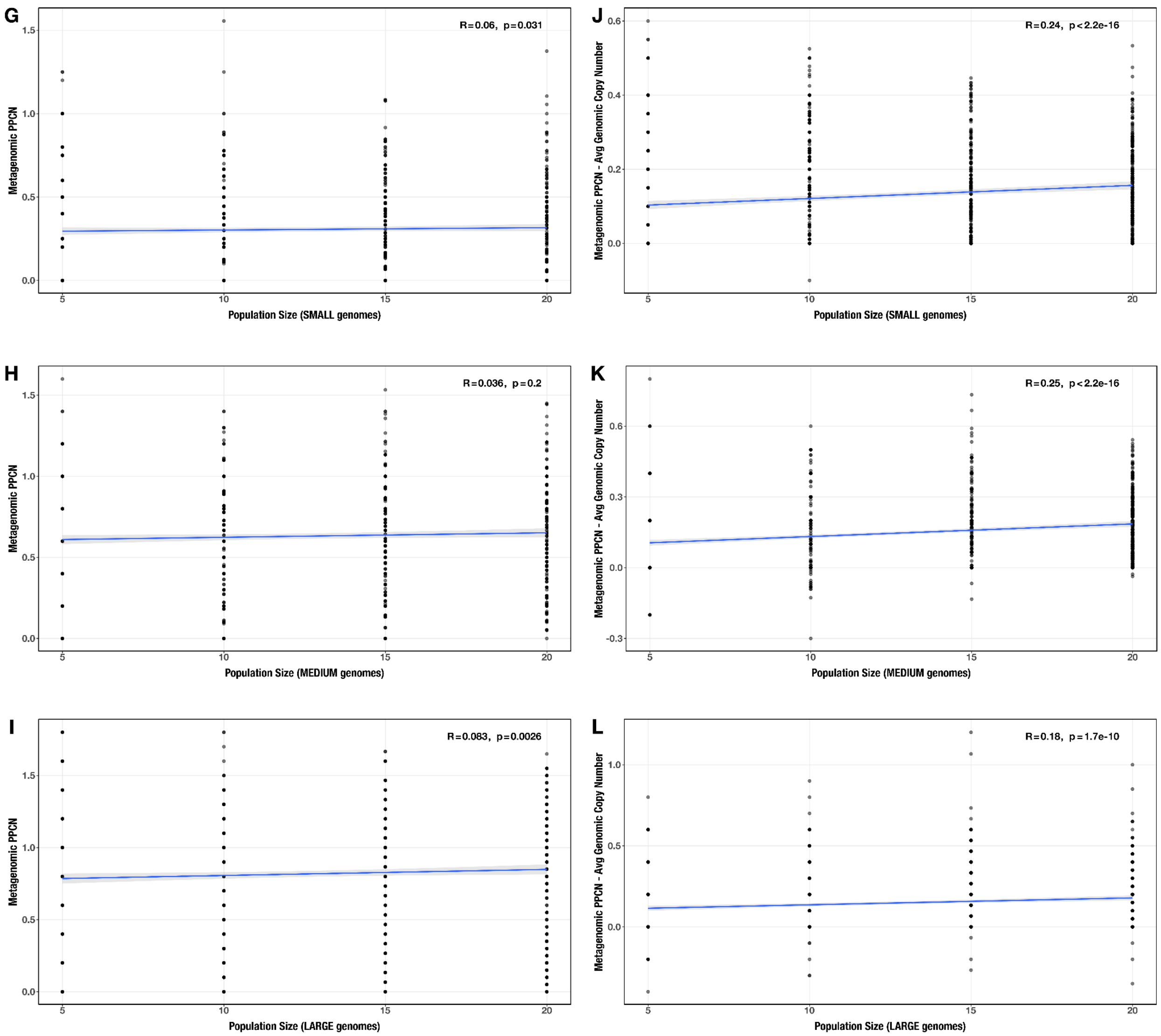


**Supplementary Figure 18**. Correlations between community size and (**A** – **C**, **G** – **I**) PPCN or (**D** – **F**, **J** – **L**) PPCN error relative to average genomic copy number for each genome size category (**A**/**D**/**G**/**J**: small genomes; **B**/**E**/**H**/**K**: medium genomes; **C**/**F**/**I**/**L**: large genomes), across all modules (**A** – **F**) or the subset of IBD-enriched modules (**G** – **L**) in the community size test case. The Spearman's correlation coefficients and p-values are shown in the top-right corner of each plot, and regression lines are plotted in blue.

Finally, the accuracy of estimating community size was negatively correlated with community size for communities of small and medium-sized genomes ($R <= -0.61$, $p < 1e-04$), meaning our method more accurately predicts the size of smaller communities. However, this trend disappeared for communities of large genomes, in which community sizes were predicted with 100% accuracy (Supplementary Figure 19). Similar to the observation in the genome size test case where the presence of more large genomes increased the accuracy of community size estimates, this trend might result from a tendency of larger genomes to contain more complete sets of SCGs.

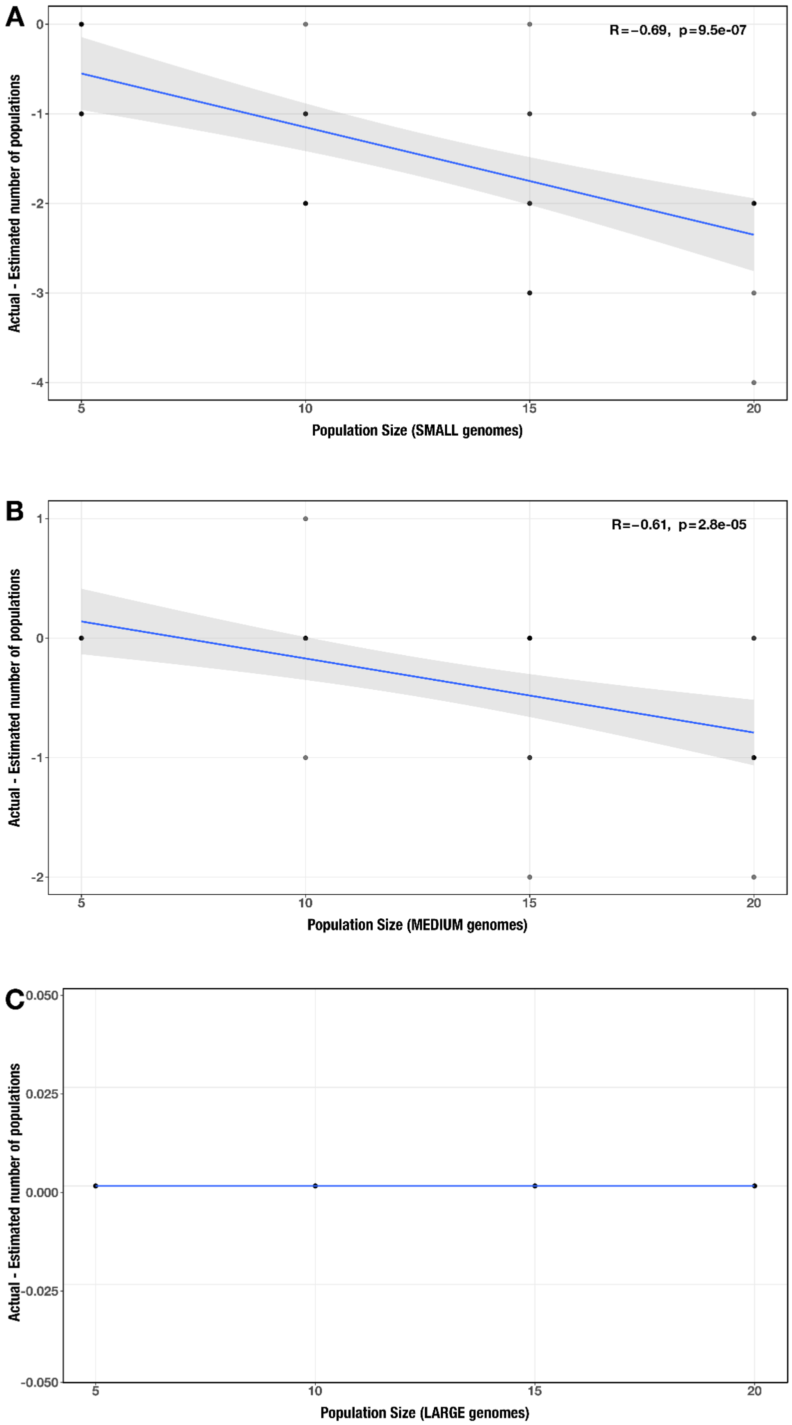


**Supplementary Figure 19.** Correlations between community size and error in community size estimate (relative to actual community size) for each genome size category (**A**: small; **B**: medium; **C**: large) in the community size test case. The Spearman's correlation coefficients and p-values are shown in the top-right corner of each plot, and regression lines are plotted in blue.

#### The impact of diversity

Our final test case assessed how diversity level influences the PPCN approach. We generated 100 synthetic metagenome assemblies, each containing 20 randomly-selected genomes from a different number of phyla (see Supplementary Methods). The number of genomes representing a given phylum was approximately equal in a given sample; however, genome size was not necessarily consistent. Due to the lack of genome size groups in this test case, we analyzed all 100 samples collectively.

The number of phyla represented in a given community was weakly correlated with both PPCN and PPCN accuracy ($0.02 < R < 0.04$, Supplementary Table 6b, Supplementary Figure 20), with only slightly stronger correlations for the subset of IBD-enriched modules ($0.04 < R < 0.07$, Supplementary Table 6b, Supplementary Figure 20). The accuracy of community size estimates were not significantly correlated with diversity level (Supplementary Figure 21).

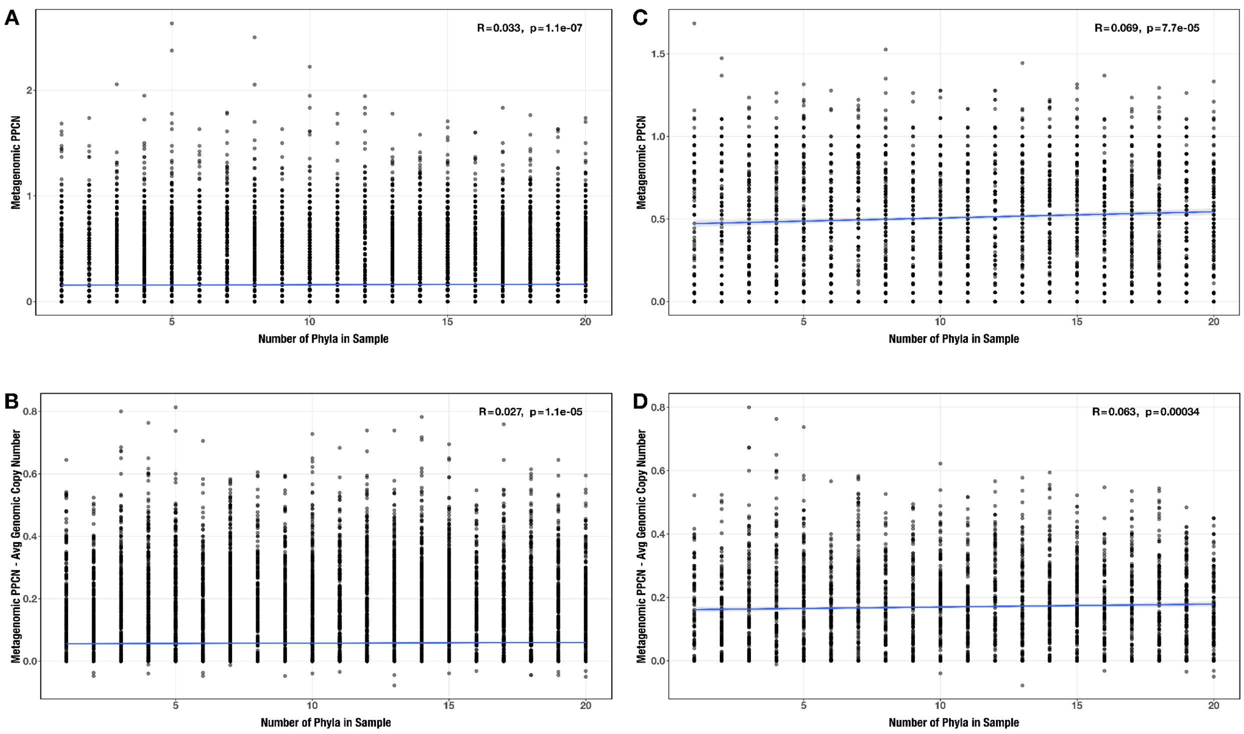


**Supplementary Figure 20.** Correlations between number of phyla and (**A**/**C**) PPCN or PPCN accuracy relative to average genomic copy number (**B**/**D**), for all modules (**A**/**B**) or the subset of IBD-enriched modules (**C**/**D**) in the diversity test case. The Spearman's correlation coefficients and p-values are shown in the top-right corner of each plot, and regression lines are plotted in blue.

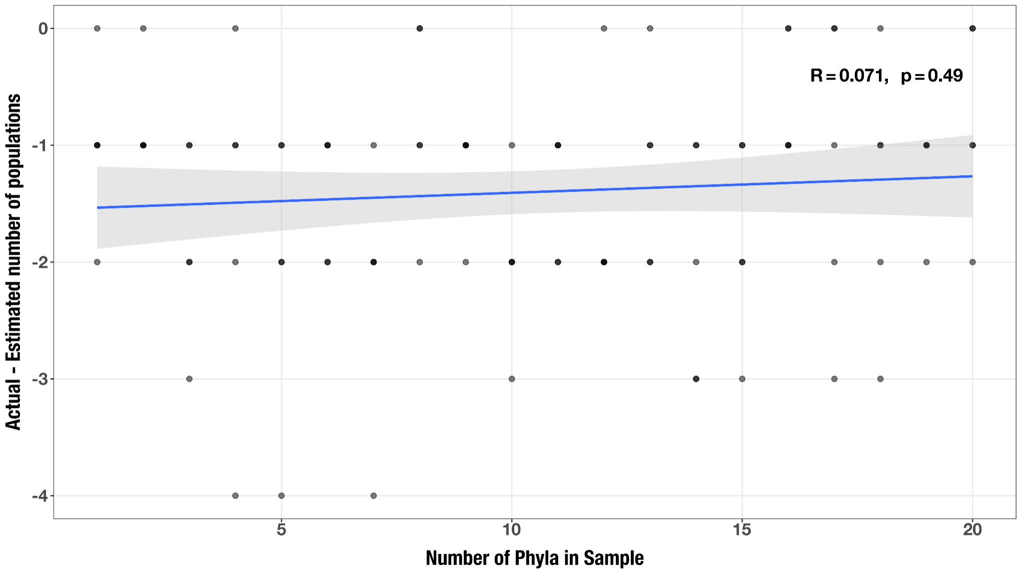


**Supplementary Figure 21.** Correlation between number of phyla and accuracy of community size estimates (relative to actual community size) in the diversity test case. The Spearman's correlation coefficient and p-value are shown in the top-right corner of each plot, and the regression line is plotted in blue.

#### Overall impact on the comparison between healthy and IBD gut metagenomes

In summary, our validation strategy revealed good accuracy at estimating metagenome-level metabolic capacity relative to our genome-level knowledge in the simulated data. While it often underestimated average genomic completeness by ignoring partial copies of metabolic pathways and often overestimated average genomic copy number due to the effect of pathway complementarity between different community members, the magnitude of error was overall limited in range and the error distributions were centered at or near 0. Furthermore, we observed these broad error trends in all cases we tested, and therefore we expect that they would also apply to both sample groups in our comparative analysis. Thus, we next considered how the PPCN approach might have influenced our analyses that considered metagenomes from healthy individuals and from those who have IBD – two groups that differed from one another with respect to some of the variables considered in our tests.

Most of the correlations between PPCN or PPCN accuracy and sample parameters were weak, yet significant (Table 1). They showed that community size and diversity level have limited influence on the PPCN calculation, while genome size does not influence its accuracy. The only exception was the moderate correlation between PPCN and genome size, particularly for the subset of IBD-enriched pathways. It was a negative correlation with the proportion of small genomes in a metagenome, indicating that PPCN values for these pathways are larger when there are more large genomes in the community and suggesting that these pathways tend to occur frequently in larger genomes. This is in line with our observation that IBD communities contain more large genomes and therefore confirms our interpretation that the populations surviving in the IBD gut microbiome are those with the genomic space to encode more metabolic capacities.

| Correlation? | PPCN (all) | PPCN Error (all) | PPCN (enriched) | PPCN Error (enriched) |
|---|---|---|---|---|
| Genome size (proportion of smaller genomes) | NO (-R < 0.09) | NO (+R ≤ 0.11) | YES (-R ≥ 0.48) | NO* (\|R\| < 0.13) |
| Community size (# of populations) | NO* (+R < 0.04) | SOME (+R ≤ 0.16) | NO* (+R < 0.085) | SOME (+R ≤ 0.25) |
| Diversity (# of phyla) | NO (+R = 0.033) | NO (+R < 0.04) | NO (+R = 0.069) | NO (+R < 0.065) |

**Table 1.** Summary of correlation relationships between PPCN, PPCN accuracy, and sample parameters from Supplementary Table 6b. We labeled each pair according to the strength of correlation indicated with the R value: NO, very weak with |R| ≤ 0.19; SOME, weak with 0.19 < |R| ≤ 0.39 ; YES, moderate to strong with |R| > 0.39. A '+' sign in front of the R value indicates that all R values were positive, a '-' sign indicates that all were negative, and the absolute value sign indicates that they had mixed signs. An asterisk (*) indicates that some of the correlations were non-significant (p > 0.05).

If we consider even the weak correlations, two of those relationships indicate that our approach would be more accurate for IBD metagenomes than for healthy metagenomes. For instance, PPCN accuracy was slightly higher for smaller communities (as in IBD samples), with a weakly positive correlation between PPCN error and community size. It was also slightly more accurate for less diverse communities (as in IBD samples), with a weakly positive correlation between PPCN error and number of phyla. The only opposing trend was the weakly positive correlation between PPCN error and proportion of smaller genomes, which favors higher accuracy in communities with smaller genomes (as in healthy samples). Given that our analysis focuses on the pathways enriched in IBD samples, an overall higher accuracy in IBD samples would increase the confidence in our enrichment results.

We also examined the accuracy of our method to predict the number of populations within a metagenome based on the distribution and frequency of single-copy core genes (i.e., the denominator in the calculation of PPCN). Our benchmarks show that the estimates are overall accurate, where most errors reflect a negligible amount of underestimations of the actual number of populations. Errors occurred more frequently for the realistic synthetic assemblies generated from simulated short read data than for the ideal synthetic assemblies generated from the combination of genomic contigs. The correlations between estimation accuracy and sample parameters indicated that the population estimates are more accurate for smaller communities and communities with more large genomes, as in IBD samples (Table 2). Thus, this method is more likely to underestimate the community size in healthy samples, and these errors could lead to overestimation of PPCN in healthy samples relative to IBD samples. Thus, the enrichment of a given pathway in the IBD samples would have to overcome its relative overestimation in the healthy sample group, making it more likely that we identified pathways that were truly enriched in the IBD communities.

| **Correlation?** | **Error in community size estimate** |
|---|---|
| Genome size<br>(proportion of smaller genomes) | YES* (0.14 < \|R\| < 0.72) |
| Community size<br>(# of populations) | YES (0.6 < -R < 0.7)<br>[except for large genomes] |
| Diversity<br>(# of phyla) | NO* (+R = 0.071) |

**Table 2.** Summary of correlation relationships between the accuracy of community size estimates and sample parameters from Supplementary Table 6b. We labeled each pair according to the strength of correlation indicated with the R value: NO, very weak with $|R| \leq 0.19$; SOME, weak with $0.19 < |R| \leq 0.39$ ; YES, moderate to strong with $|R| > 0.39$. A '+' sign in front of the R value indicates that all R values were positive, a '-' sign indicates that all were negative, and the absolute value sign indicates that they had mixed signs. An asterisk (*) indicates that some of the correlations were non-significant ($p > 0.05$).

Overall, the consideration of our simulations in the context of healthy vs IBD metagenomes suggest that slight biases in our estimates as a function of unequal diversity with sample groups should have driven PPCN calculations towards a conclusion that is opposite of our observations under neutral conditions. Thus, clear differences between healthy vs IBD metagenomes that overcome these biases suggest that biology, and not potential bioinformatics artifacts, is the primary driver of our observations.

### Supplementary Methods

Below are the methods used for validation of our approach on simulated metagenomic data. All custom scripts and auxiliary data files are available on Figshare at DOI:10.6084/m9.figshare.26038018. A reproducible workflow for this section is also available at https://merenlab.org/data/ibd-gut-metabolism/.

**Generation of synthetic communities.** We generated synthetic metagenomic communities using species cluster representative genomes from the Genome Taxonomy Database (GTDB) v95 (Parks et al. 2022), the same database utilized in our main analysis. We used genome length to group each genome into different size categories: 7,238 'small' genomes of < 2 Mbp; 18,416 'medium'-sized genomes of 2 Mbp to < 5 Mbp; and 6,254 'large' genomes of 5 Mbp to 20 Mbp. Two especially large cyanobacterial genomes with over 20 Mbp were excluded from further analysis.

We then used custom scripts to randomly combine these genomes into synthetic communities for each simulated test case. To test the robustness of our approach to genome size, we combined 20 genomes of two size categories (small and medium; small and large; medium and large) in different proportions in each community, varying the number of genomes from the first size group from 0 to 20 and varying the number from the second size group from 20 to 0. We randomly selected genomes from each size group without replacement, and generated 3 samples per proportion value for each size category pair. This produced a total of 189 synthetic communities representing a gradient of mixed genome sizes for the genome size test case. To test the robustness of our approach to community size, we combined 5, 10, 15, or 20 randomly-selected genomes (without replacement) of a single size category (small, medium, or large) in each community. We generated 40 synthetic communities (10 for each community size value) for each genome size category to obtain a total of 120 samples in the population size test case. Finally, to test the robustness of our approach to different diversity levels, we generated communities containing a total of 20 random bacterial genomes, varying the number of phyla represented in the community from 1 to 20 and ensuring that there was an approximately equal number of genomes from each phylum in the sample. Each phylum was selected randomly without replacement from the set of unique bacterial phyla represented in the GTDB. Phyla containing less than 20 genomes were excluded. We randomly selected genomes from each phylum without replacement, and we generated 5 synthetic communities for each diversity level to obtain 100 samples representing a gradient of diversity for the diversity test case.

**Generation of 'ideal' synthetic metagenomes.** The 'ideal' synthetic metagenomes represent the scenario in which all microbial populations in a metagenome are fully represented in the assembly (at least to the level of completion for each genome in the GTDB). For each of the aforementioned synthetic communities (in each test case), we concatenated the individual genome FASTA files into one FASTA containing the contig sequences of all genomes in the community. We then applied the approach (see main Methods) used in our main analysis to estimate population sizes, calculate metagenomic copy number, and compute per-population copy number for metabolic pathways in each synthetic metagenome. We used the same version of the KEGG database (from December 2020) as in our main analysis for consistency with those results.

**Generation of 'realistic' synthetic metagenomes.** We generated 'realistic' synthetic metagenomes for the 189 samples in the genome size test case by simulating paired-end short reads from genomic data. To ensure that the populations in these samples had realistic relative abundances, we first computed a relative abundance curve that is typical of healthy human gut metagenomes. For this task, we used the relative abundance data computed by MetaPhlAn 3 for publicly-available gut metagenomes (Beghini et al. 2021). We filtered this dataset to keep only species-level relative abundance values in 662 healthy control metagenomes, sorted the values in descending order, and kept only the top 20 relative abundances in each sample. We then scaled each sample's data to have the same maximum relative abundance value, averaged the values at each rank, scaled the resulting averages to obtain a minimum average relative abundance value of 1, rounded the averages into integer coverage values, and multiplied the coverages by 20 to obtain high enough sequencing depth for proper assembly of the synthetic metagenomes. This process yielded coverage values of 20x – 420x for individual populations within a 'typical' healthy human gut metagenome (Supplementary Figure 14).

We used a custom script to randomly select one of these typical coverage values for each genome in a given synthetic community. With the program `gen-paired-end-reads` (https://github.com/merenlab/reads-for-assembly), we generated synthetic short reads from each input genome with the specified target coverage value, only simulating from contigs with length ≥1000 bp. The simulation parameters for the paired-end reads were as follows: read length of 150, inner distances with a mean of 100 bp and standard deviation of 10 bp, and a 0.05% sequencing error rate. After simulating the short reads, we assembled each individual synthetic metagenome with MEGAHIT (Li et al. 2015), specifying `--min-contig-len 1000`. From there we followed the same approach used in our main analysis and for the 'ideal' synthetic metagenomes to obtain per-population copy numbers for metabolic pathways.

**Metabolism estimation for individual genomes.** To obtain genomic pathway prediction metrics for comparison to the metagenomic values, we ran `anvi-estimate-metabolism` on each genome used to generate the synthetic communities. We used the following parameters: `--include-zeros`, `--matrix-format`, and `--add-copy-number` (which were the same parameters used for estimating metabolism in the synthetic metagenomes).

**Analysis of estimation accuracy and robustness to sample parameters.** We wrote custom scripts to compare the PPCN values from metagenomes to the metabolism estimation data from individual genomes, and to compare the estimated number of populations to the true community size. We then used a custom R script for analysis and visualization of these comparisons, which include statistical analysis, Spearman's correlation with the sample parameters, and figure generation.

DeJongh, Matthew, Kevin Formsma, Paul Boillot, John Gould, Matthew Rycenga, and Aaron Best. 2007. "Toward the Automated Generation of Genome-Scale Metabolic Networks in the SEED." *BMC Bioinformatics* 8 (April): 139.

Dias, Oscar, Miguel Rocha, Eugénio C. Ferreira, and Isabel Rocha. 2015. "Reconstructing Genome-Scale Metabolic Models with Merlin." *Nucleic Acids Research* 43 (8): 3899–3910.

Fang, Xin, Colton J. Lloyd, and Bernhard O. Palsson. 2020. "Reconstructing Organisms in Silico: Genome-Scale Models and Their Emerging Applications." *Nature Reviews. Microbiology* 18 (12): 731–43.

Faria, José P., Miguel Rocha, Isabel Rocha, and Christopher S. Henry. 2018. "Methods for Automated Genome-Scale Metabolic Model Reconstruction." *Biochemical Society Transactions* 46 (4): 931–36.

Geller-McGrath, D., Kishori M. Konwar, V. P. Edgcomb, M. Pachiadaki, J. W. Roddy, T. J. Wheeler, and J. E. McDermott. 2023. "MetaPathPredict: A Machine Learning-Based Tool for Predicting Metabolic Modules in Incomplete Bacterial Genomes." *bioRxiv*. https://doi.org/10.1101/2022.12.21.521254.

Gu, Changdai, Gi Bae Kim, Won Jun Kim, Hyun Uk Kim, and Sang Yup Lee. 2019. "Current Status and Applications of Genome-Scale Metabolic Models." *Genome Biology* 20 (1): 121.

Henry, Christopher S., Matthew DeJongh, Aaron A. Best, Paul M. Frybarger, Ben Linsay, and Rick L. Stevens. 2010. "High-Throughput Generation, Optimization and Analysis of Genome-Scale Metabolic Models." *Nature Biotechnology* 28 (9): 977–82.

Kanehisa, Minoru, Susumu Goto, Masahiro Hattori, Kiyoko F. Aoki-Kinoshita, Masumi Itoh,

Shuichi Kawashima, Toshiaki Katayama, Michihiro Araki, and Mika Hirakawa. 2006. “From Genomics to Chemical Genomics: New Developments in KEGG.” *Nucleic Acids Research* 34 (Database issue): D354–57.
Karp, Peter D., Suzanne M. Paley, Peter E. Midford, Markus Krummenacker, Richard Billington, Anamika Kothari, Wai Kit Ong, Pallavi Subhraveti, Ingrid M. Keseler, and Ron Caspi. 2015. “Pathway Tools Version 24.0: Integrated Software for Pathway/genome Informatics and Systems Biology.” *arXiv [q-bio.GN]*. arXiv. http://arxiv.org/abs/1510.03964.
Li, Dinghua, Chi-Man Liu, Ruibang Luo, Kunihiko Sadakane, and Tak-Wah Lam. 2015. “MEGAHIT: An Ultra-Fast Single-Node Solution for Large and Complex Metagenomics Assembly via Succinct de Bruijn Graph.” *Bioinformatics* 31 (10): 1674–76.
Machado, Daniel, Sergej Andrejev, Melanie Tramontano, and Kiran Raosaheb Patil. 2018. “Fast Automated Reconstruction of Genome-Scale Metabolic Models for Microbial Species and Communities.” *Nucleic Acids Research* 46 (15): 7542–53.
Mendoza, Sebastián N., Brett G. Olivier, Douwe Molenaar, and Bas Teusink. 2019. “A Systematic Assessment of Current Genome-Scale Metabolic Reconstruction Tools.” *Genome Biology* 20 (1): 158.
Moriya, Yuki, Masumi Itoh, Shujiro Okuda, Akiyasu C. Yoshizawa, and Minoru Kanehisa. 2007. “KAAS: An Automatic Genome Annotation and Pathway Reconstruction Server.” *Nucleic Acids Research* 35 (Web Server issue): W182–85.
Orth, Jeffrey D., Ines Thiele, and Bernhard Ø. Palsson. 2010. “What Is Flux Balance Analysis?” *Nature Biotechnology* 28 (3): 245–48.
Pacheco, Alan R., Mauricio Moel, and Daniel Segrè. 2019. “Costless Metabolic Secretions as Drivers of Interspecies Interactions in Microbial Ecosystems.” *Nature Communications* 10 (1): 103.
Palù, Matteo, Arianna Basile, Guido Zampieri, Laura Treu, Alessandro Rossi, Maria Silvia Morlino, and Stefano Campanaro. 2022. “KEMET - A Python Tool for KEGG Module Evaluation and Microbial Genome Annotation Expansion.” *Computational and Structural Biotechnology Journal* 20 (March): 1481–86.
Parks, Donovan H., Maria Chuvochina, Christian Rinke, Aaron J. Mussig, Pierre-Alain Chaumeil, and Philip Hugenholtz. 2022. “GTDB: An Ongoing Census of Bacterial and Archaeal Diversity through a Phylogenetically Consistent, Rank Normalized and Complete Genome-Based Taxonomy.” *Nucleic Acids Research* 50 (D1): D785–94.
Sen, Partho, and Matej Orešič. 2019. “Metabolic Modeling of Human Gut Microbiota on a Genome Scale: An Overview.” *Metabolites* 9 (2). https://doi.org/10.3390/metabo9020022.
Shaffer, Michael, Mikayla A. Borton, Bridget B. McGivern, Ahmed A. Zayed, Sabina Leanti La Rosa, Lindsey M. Solden, Pengfei Liu, et al. 2020. “DRAM for Distilling Microbial Metabolism to Automate the Curation of Microbiome Function.” *Nucleic Acids Research* 48 (16): 8883–8900.
Ye, Yuzhen, and Thomas G. Doak. 2009. “A Parsimony Approach to Biological Pathway Reconstruction/inference for Genomes and Metagenomes.” *PLoS Computational Biology* 5 (8): e1000465.
Zhou, Zhichao, Patricia Q. Tran, Adam M. Breister, Yang Liu, Kristopher Kieft, Elise S. Cowley, Ulas Karaoz, and Karthik Anantharaman. 2022. “METABOLIC: High-Throughput Profiling of Microbial Genomes for Functional Traits, Metabolism, Biogeochemistry, and Community-Scale Functional Networks.” *Microbiome* 10 (1): 33.
Zimmermann, Johannes, Christoph Kaleta, and Silvio Waschina. 2021. “Gapseq: Informed Prediction of Bacterial Metabolic Pathways and Reconstruction of Accurate Metabolic Models.” *Genome Biology* 22 (1): 81.
